# TolRad: A model for predicting radiation tolerance using Pfam annotations identifies novel radiosensitive bacterial species from reference genomes and MAGs

**DOI:** 10.1101/2023.11.02.562514

**Authors:** Philip Sweet, Matthew Burroughs, Sungyeon Jang, Lydia Contreras

**Affiliations:** McKetta Dept. of Chemical Engineering, University of Texas at Austin

## Abstract

The trait of ionizing radiation (IR) tolerance is variable between bacterial species, with radiosensitive bacteria succumbing to acute doses around 100Gy and extremophiles able to survive doses exceeding 10,000Gy. While survival screens have identified multiple highly radioresistant bacteria, such systemic searches have not been conducted for radiosensitive bacteria. The taxonomy-level diversity of IR of intolerance across bacteria is poorly understood, as are genetic elements that influence IR sensitivity. Using the protein domain frequencies from 61 bacterial species with experimentally determined IR D10 values (the dose at which only 10% of the population survives) we trained TolRad, a random forest binary classifier, to distinguish between radiosensitive bacteria (D10 < 200Gy) and radiation tolerant bacteria (D10 > 200Gy). On the hidden species, TolRad had an accuracy of 0.900. We applied TolRad to 152 UniProt-hosted bacterial proteomes, including 37 strains from the ATCC Human Microbiome Collection, and classified 34 species as radiosensitive. Whereas IR intolerance (D10 < 200Gy) in the training dataset had been confined to the phylum *Proteobacterium*, this initial TolRad screen identified radiosensitive bacteria in 2 additional phyla. We experimentally validated the predicted radiosensitivity of a key species of the human microbiome from the *Bacteroidota* phyla. To demonstrate that TolRad can be applied to Metagenome-Assembled Genome (MAGs), we tested the accuracy of TolRad on Egg-NOG assembled proteomes (0.965) and partial proteomes. Finally, three collections of MAGs were screened using TolRad, identifying further phylum with radiosensitive species and suggesting that environmental conditions influence the abundance of radiosensitive bacteria.

**Importance:** Bacterial species have vast genetic diversity, allowing for life in extreme environments and the conduction of complex chemistry. The ability to harness the full potential of bacterial diversity is hampered by the lack of high-throughput experimental or bioinformatic methods for characterizing bacterial traits. Here, we present a computational model that uses *de novo* generated genome annotations to classify a bacterium as tolerant of ionizing radiation (IR) or as radiosensitive. This model allows for rapid screening of bacterial communities for low-tolerance species that are of interest for both mechanistic studies into bacterial sensitivity to IR and biomarkers of IR exposure.

## Introduction

Advances in sequencing technology and genome assembly algorithms have led to an exponential increase in the number and diversity of available bacterial genomes (1). Third-generation sequencing platforms and single-cell sequencing methods can capture an even greater genetic diversity of bacterial communities (1). These new methods do not require the ability to isolate and culture individual strains within a community--a barrier that had previously limited the diversity of bacteria that could be studied (2). While there are still challenges to assembling metagenomes (3), current methods have allowed for the description of thousands of new bacterial genomes, such as from ocean water (4), the human microbiome (5), soil (6) and even the international space station (7). This rise in genomic data highlights the need for computational tools for characterizing bacterial traits. Experimentally established traits, such as respiratory preference, gram stain, carbon source utilization, and antibiotic tolerance have historically been used to describe new bacterial species (8), however, these established methods are not suited for community-level analysis. Experimentally determining the traits of entire communities of bacteria is impractical; yet, to understand how complex communities of bacteria are interacting, knowing which traits are associated with which species is essential (3).

To aid in the interpretation of novel bacterial genomes, predictive algorithms have been developed that can generate genome annotations directly from genomic sequences. Genome annotation tools can identify protein-coding regions (Prodigal) (9), determine protein structure (AlphaFold) (10), and suggest possible protein functions (Pfam) (11). These tools allow for the description of individual proteins, but the connection between collections of proteins and specific phenotypic traits is still poorly understood. In response to these issues, statistical models that connect genome annotations to phenotypic traits have been developed. Models have been written to predict traits such as the nature of prophages (BACPHLIP) (12) (Hockenberry and Wilke, 2021), metabolic preference (13), virulence (14) and antibiotic tolerance (15) (16). Often, these models use Pfam domains, annotations assigned to protein sequences using a set of hidden Markov models (11), to inform the classification. The ability to predict bacterial tolerance for stress from such genome annotations is currently limited to antibiotic resistance (15, 16).

There is a renewed interest in understanding native bacterial tolerance for ionizing radiation (IR). IR is a complex stress that threatens the stability of the bacterial genome. A greater understanding of bacterial sensitivity to IR has implications for diverse research topics, including the hardening of bacteria against IR for bioremediation of radioactive sites (17), developing bacterial biomarkers of IR exposure (18) and understanding the risk posed to the human microbiome by space flight (19) and radiation therapy (20). For example, a recent metareview of the response of the human microbiome to radiation noted changes in species diversity and abundance after radiation exposure but highlighted the lack of clarity around the susceptibility of these species to IR (21).

Exposure of bacterial cells to IR causes both direct damage when ionized particles collide with macromolecules (22), and indirect damage generated by reactive oxygen species (ROS) resulting from radiolysis. DNA damage (23) and protein oxidation (24) have both been observed after the exposure of cells to IR. To compare IR tolerance between bacterial species, the **D**ose at which only **10**% of the exposed cells will produce viable colonies **(D10**) is used. The tolerance of bacteria for IR varies greatly between species (25). For instance, the extremophile *Deinococcus radiodurans (D. radiodurans*) has a D10 of 12,000Gy whereas the radiosensitive bacterium *Shewanella oneidensis* (*S. oneidensis*) has a D10 of 70Gy (26). Determining the D10 dose of a bacterium requires access to a powerful irradiation source and the ability to individually culture the bacterium in question. These demands have limited the number of bacteria for which D10 values are available. Historically, the identification of radiation-resistant bacteria has been advanced by the need to understand the dose of IR required for the sterilization of medical compounds (27), food (28), and wastewater (29). Work has also been done to identify extremophiles from off-world analog sites such as the Taklimakan desert (30). These research motives have favored the discovery of radioresistant bacteria over radiosensitive bacteria. And yet, when trying to understand the disruption of important bacterial communities, such as the gut microbiome (19) (21), that occurs after IR exposure, there is a limited understanding of which species are most sensitive to radiation, and what factors make certain bacteria more or less sensitive to IR. In part, this limitation is because the genetic diversity of radiosensitive bacteria is poorly understood, with generalizations about radiation tolerance being limited to observations, such as gram-negative bacteria are generally less tolerant than gram-positive bacteria (31). Studies of the mechanism behind bacterial sensitivity to IR are hindered by the small number of radiosensitive bacteria currently known. Experimental characterization has only discovered 14 bacteria with D10 values less than 200Gy (Sup. Table 1), and all these bacteria belong to the *Protobacterium* phylum.

The best-understood predictor of IR tolerance in bacteria is the intracellular ratio of manganese to iron (Mn/Fe) (23). Research comparing the radiosensitive bacterium *S. oneidensis* and extremely radioresistant bacterium *D. radiodurans* (26) (24) has found that the intracellular ratio of Mn/Fe is correlated with IR tolerance. This ratio has been found to generally be predictive of microorganism IR tolerance, including across five bacterial species (23). The relationship between the intracellular ratio of Mn/Fe and IR tolerance is explained by the opposing effect these ions have on the spread of ROS. Iron furthers the spread of ROS through Fenton chemistry, whereas manganese is an antioxidant, acting as a sponge of ROS. While determining the intracellular ratio of Mn/Fe does not require an IR source, it does require isolated culture growth. Complicating the interpretation of Mn/Fe ratios, the intracellular ratio of Mn/Fe has been shown to be influenced by growth media (24). To date, a correlation between gene or protein frequency and levels of intracellular ratios of Mn/Fe between bacteria has not been determined.

To identify novel radiosensitive bacterial species, we constructed a model capable of using the frequency of Pfam domains to identify bacterial species that haver a low survival threshold for IR. In this paper, we describe the construction and validation of the <u>Tol</u>erance for <u>Rad</u>iation (TolRad) model. We also describe the application of TolRad to pre-annotated reference proteomes and Metagenome Assembled Genomes (MAGs). TolRad is a random forest binary classifier that uses the relative frequency of Pfam annotations within a bacterial proteome to classify a genomic assembly as coming from a bacterium that is tolerant of (D10 > 200Gy) or sensitive to (D10 < 200Gy) IR exposure. To build TolRad, a diverse set of 61 bacteria, with associated Pfam annotations and experimentally determined D10 values, was split 70/30 into a Train Set and Test Set. The final TolRad model utilized 4 Pfam Domains (PF03466, PF07992, PF00300, and PF00849). The accuracy of TolRad on the Train Set was 0.875 Interestingly, it is the frequency of the Pfam domain PF07992 (pyridine nucleotide-disulfide oxidoreductase) that has the greatest contribution to the classification of species as radiosensitive. To validate the ability of TolRad to classify species it was not trained on, the model was applied to the Test Set, on which it was 0.900 accurate.

By applying TolRad to a collection of UniProt-assembled proteomes, 34 species were classified as putative radiosensitive. Of particular interest was the prediction that 19 of the 29 *Bacteroidetes* species, many which are abundant in the human gut, were classified as radiosensitive. One of the bacteria with predicted IR intolerance was the key human gut commensal *Bacteroides thetamicrone* (*B. thetamicrone*). As no members of the *Bacteroidota* phylum have yet been experimentally characterized as having a low survival threshold to IR, we validated that *B. thetamicrone* indeed had a D10 value below 200Gy. The ability of TolRad to correctly identify radiosensitive bacteria from *de novo* annotated Pfam domains was tested by reannotating the genomes of the Train/Test Set with EggNOGG-Mapper (32). TolRad suffered no decrease in accuracy (0.97) on the EggNOGG-Mapper (32) annotated genomes. We then applied TolRad to MAGs from three previously published datasets: a set of human microbiome bacteria (HMB) (Han et al., 2016), a set of bacteria collected from a glacial stream in the Canadian high Arctic (CHA) (Trivedi et al., 2020) and a collection from the deep ocean (33). Broadly speaking, TolRad predicted a greater ratio of putative radiosensitive bacteria from the deep ocean (30.7%) and human gut microbiome (10.9 %), compared with the CHA (6.9%) and human skin microbiome (0%). Screening these MAGs further expanded the diversity of putative radiosensitive species and suggests that environmental conditions can influence IR-tolerance.

In summary, we demonstrate that a random forest classifier, built using a dataset of previously published survival values and associated Pfam annotations, can be used to predict bacterial stress tolerance. A similar workflow, as we describe for the construction of TolRad, could be used to develop a predictive classifier for other environmental stresses. Specifically, using TolRad, we have identified novel species of radiosensitive bacteria from the human microbiome. Broadening the diversity of known radiosensitive bacteria will be vital for future studies into the mechanisms of radiation tolerance and for identifying bacterial biomarkers of radiation exposure.

## Results

### Collection of the Train/Test Set

TolRad was trained and tested on a manually curated dataset of 120 experimentally determined bacterial D10 values representing 61 species (**Sup. Table 1, Sup. Table 2**) and the associated frequency of Pfam domains, referred to as the Train/Test Set. Only D10s determined in liquid media or PBS and exposed at room temperature were included. When multiple D10s were found, the mean was used. The Train/Test Set contains a diversity of bacteria from 7 different phyla, species that are anaerobic, aerobic and, facultative anaerobic as well as gram-positive and negative species (**Sup. Table 1**).

### Establishing the IR Sensitivity Cut-off

Within the Train/Test Set, the D10 values varied from 60Gy to 15,000Gy, with a heavy rightward skew (**Figure 1A**). Bacteria previously described as extremely radiation-tolerant (D10 > 1500Gy) (**Sup. Table 1**) were found within the rightward tail. We defined the bacteria of the lower quarter of the distribution (**Figure 1A**) as radiosensitive, resulting in a cut-off of D10 < 200Gy. Since we were most interested in identifying radiosensitive species, we combined the “Moderately Tolerant” and “Extremely Tolerant” categories into a “Tolerant” category (D10 > 200Gy). Using a T-Test we demonstrated that, within the Train/Test Set, there was not a statistically significant difference (p< 0.05) in optimal growth temperature or the genome GC content between the radiosensitive and tolerant species (**Sup. Figure 1**). We observed that none of the gram-positive species were radiosensitive. At the phylogenetic level, all the radiosensitive species were *Proteobacteria.* The represented bacteria from the remaining phylum (*Actinobacteria*, *Aquificota*, *Firmicutes*, *Bacteroidetes* and *Deinococi*) were classified as tolerant of IR. TolRad was trained specifically to differentiate radiosensitive species (D10 < 200Gy) from tolerant species (D10 > 200Gy) and does not assign relative tolerance, only a binary classification.

**Figure 1:**
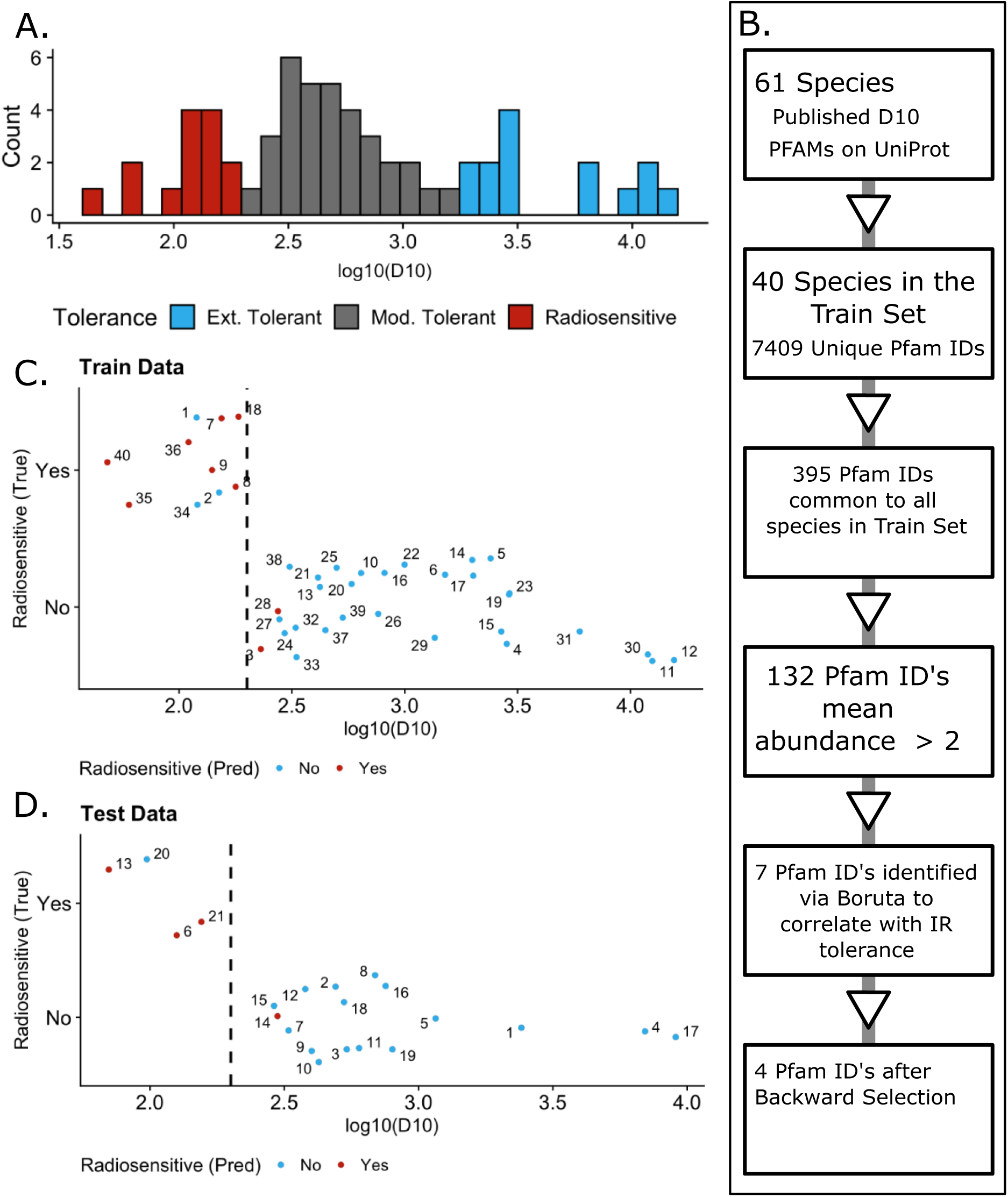
Construction of TolRad. A. Histogram distribution of the Train/Test set. Each count is the mean D10 of a unique species. A total of 61 species were used to train the model, representing 121 experimentally determined D10 values. B. Workflow used for training TolRad. C. Performance of TolRad on the Train Set. Numbers are a unique species. 1 *Acinetobacter calcaeceticus*, 2 *Aeromonas hydrophila*, 3 *Aeromonas salmonicida*, 4 *Aquifex pyrophilus*, 5 *Bacillus pumilus*, 6 *Bacillus sphaericus*, 7 Campylobacter coli, 8 *Campylobacter jejuni*, 9 *Campylobacter lari*, 10 *Coxiella burnetiid*, 11 *Deinococcus geothermalis*, 12 *Deinococcus radiodurans*, *13 Enterobacter sp. BIGb0383*, 14 *Enterococcus faecium*, 15 *Enterococcus faecalis*, 16 *Escherichia coli*, 17 *Kineococcus radiotolerans*, 18 *Klebsiella variicola*, *19 Kocuria rhizophila*, 20 Lactococcus lactis, 21 *Listeria monocytogenes*, 22 *Methylobacterium radiotoleran*, 23 *Micrococcus luteus*, 24 Morganella morganii, 25 *Mycobacterium smegmatis*, 26 *Mycobacterium tuberculosis*, 27 *Proteus vulgaris*, 28 *Pseudomonas putida*, 29 *Rhodococcus erythropolis*, 30 *Rubrobacter radiotolerans*, 31 *Rubrobacter xylanophilus,* 32 *Salmonella Senftenberg*, 33 Salmonella enterica subsp. enterica serovar Heidelberg, 34 Salmonella paratyphi, *35 Serratia marcescens*, 36 *Shewanella putrefaciens*, 37 *Staphylococcus aureus* 38 *Staphylococcus epidermidis*, 39 *Stenotrophomonas maltophilia*, 40 *Vibrio parahaemolyticus*. Graph IDs can also be found in Sup. Table 3. Color denotes the classification assigned by TolRad. Red denotes a species classified as radiosensitive. Blue denotes a species classified as tolerant. The dashed line at 200Gy is the cut-off used for defining radiosensitive vs. tolerant species. D. Same as C, but for the Train Set classifications. 1 *Acinetobacter radioresistens*, 2 *Bacillus cereus*, *3 Bifidobacterium breve*, 4 *Deinococcus ficus*, 5 *Lactobacillus acidophilus*, 6 *Neisseria gonorrhoeae*, 7 *Paenibacillus amylolyticus*, 8 *Pediococcus pentosaceus*, 9 *Priestia megaterium*, 10 Salmonella enterica, 11 Salmonella muenster, 12 *Salmonella typhimurium*, 13 *Shewanella oneidensis*, 14 *Shigella boydii*, 15 *Shigella flexneri*, 16 *Shigella sonnei*, 17 *Spirosoma radiotolerans*, 18 *Streptococcus thermophilus*, 19 *Thermus thermophilus*, 20 *Vibrio cholera O1*, 21 *Yersinia enterocolitica*

### Predictor Selection

Based on the success of previous models that have predicted bacterial traits using Pfam domains (12, 34), we started by identifying Pfam domains that correlated with IR tolerance. The Train/Test Set was then randomly split, 70/30, into a Train Set and a Test Set (**Table 1 and Sup. Table 1**). The Train set was used to select predictors of IR tolerance and train the model. The Test set was totally hidden from the construction of the model. Of the 7409 unique Pfam domains present in the 40 proteomes in the Train Set, 132 were abundant (more than 2 occurrences per genome). Using the Boruta feature selection algorithm (35), the relative frequency of 7 of these 132 Pfam annotations were found to correlate with IR tolerance classification. Step-wise removal was used to select the most parsimonious model, which resulted in the final selection of 4 Pfam Domains (**Table 2**). A visualization of the predictor selection pipelines is presented in **Figure 1B**. To provide context to the biological relevance of these predictor Pfam domains, we examined the *E.coli* proteins that contain these domains (summarized in **Table 2**). We also calculated the Mean Decrease in Accuracy (**Table 2**) for each predictor in the final model to determine which domains were most important for the model’s ability to correctly classify radiation tolerance. Further details about predictor selection are provided in the Methods section.

**Table 1:** Test/Train Dataset Summary.

| Table 1: Test/Train Dataset Summary |  |  |  |
| --- | --- | --- | --- |
| Set | Total | Tolerant | Radiosensitive |
| Train | 40 | 30 | 10 |
| Test | 21 | 17 | 4 |

**Table 2:** TolRad Predictors.

| Table 2: TolRad Predictors |  |  |  |  |
| --- | --- | --- | --- | --- |
| ID | Pfam Short Name | Mean Decrease in Accuracy | Mean Occurance per Proteomoe | Example from *E.coli* and Panther Function |
| PF00300 | Pyridine nucleotide-disulphide oxidoreductase | 7.58 | 15.52 | norW: Nitric oxide reductase<br>trxB: Thioredoxin reductase<br>ndh: Type II NADH:quinone oxidoreductase |
| PF07992 | Histidine phosphatase | 11.57 | 6.87 | ais: Lipopolysaccharide metabolic process<br>gpmA/B: Glycolytic process<br>sixA: Signal transduction |
| PF03466 | LysR substrate binding domain | 8.95 | 63.42 | cysB: TF regulagor of response to x-rays<br>oxyR: ROS-responsive TF<br>lysR: TF lysine biosynthesis |
| PF13411 | HTH family regulatory protein | 5.5 | 7.366197 | cueR: Copper-responsive TF<br>zntR: Zinc-responsive TF<br>mlrA: Regulator of biofilm formation |

### Model Training and Validation

The final model was built using the RandomForest function of the R package caret (36) via a 10-cross-validation on the Train Set. The accuracy of the final model was 0.875. The final model was unable to correctly classify 5 (12.5%) of the bacteria in the Train Set, making three false negative and two false positive error (**Figure 1C, Sup. Table 3**). Encouragingly, these misclassifications were of species with D10 values within 100Gy of the 200Gy cut-off, suggesting species classified as radiosensitive are still likely to have a low D10. When the classifier was applied to the Test Set, the accuracy was 0.900. As with the Train Set, the two misclassifications were of species with a D10 close to 200Gy (**Figure 1D, Sup. Table 3**).

### Using TolRad to screen species of the Human Microbiome reveals the radiosensitivity of the *Bacteroidota* phylum

To discover novel radiosensitive bacteria within the HMB, we applied TolRad to the proteomes of 152 bacterial strains that had previously been detected within samples originating from the human microbiome. This dataset included 37 species of the official Human Microbiome Project strain collection hosted by ATCC (**Sup. Table 2, Sup. Table 4**) (37), as well as 28 species identified from the Human Skin Microbiome (38, 39), 46 species from the Human Oral Cavity Microbiome (40), and 41 species from the Human Gut Microbiome (41). For this analysis, as for the construction of TolRad, we used the Pfam domain annotations hosted on UniProt for each species, selecting species with ATCC strain ID to support downstream experimental validation. In total, we identified 34 putative radiosensitive species (**Table 3, Sup. Table 5**), of which 10 belonged to the ATCC-hosted NIH Human Microbiome Project. A secondary literature search for D10 values associated with these 34 putative radiosensitive species uncovered support for the radiosensitive classification of *Klebsiella pneumoniae* (*42*). All Uniprot Proteome classification predictions are presented in **Supplemental Table 4**.

**Table 3:** UniProt Species Summary.

| Table 3: UniProt Species Summary |  |  |  |
| --- | --- | --- | --- |
| Set | Total | Phylums | Radiosensitive (predicted) |
| NCBI/ATCC Collection | 37 | 5 | 10 |
| Human Skin | 28 | 4 | 4 |
| Human Oral Cavity | 46 | 6 | 9 |
| Human Gut | 41 | 6 | 11 |
| Total | 152 |  | 34 |

We examined the phylum level diversity of the bacteria characterized as radiosensitive from the HMB set compared with that of Train/Test. While the Train/Test Set only contained radiosensitive bacteria from the *Proteobacteria* phylum (**Figure 2A**) TolRad predicted radiosensitive species from the HMB within the *Actinobacteria*, *Firmicutes,* and *Bacteroidetes* phylum (**Figure 2A**). Of particular interest was the prediction that 19 of the 29 *Bacteroidetes* species were characterized as radiosensitive (**Sup. Table 5**). Several species of this phylum are highly abundant within the human gut microbiome, including *Bacteroides thetaiotaomicron (B. thetaiotaomicron)* (43). To validate the ability of TolRad to predict the IR tolerance of bacteria beyond the phylum represented as radiosensitive in the Train/Test set, we experimentally determined the survival of *B. thetaiotaomicron* and found the D10 to be 110Gy (**Figure 2B**). We also validated the tolerant prediction of *Acinetobacter baumannii* (**Figure 2C**). In summary, TolRad identified 34 putative radiosensitive bacteria, including 10 from the ATCC Human Microbiome Project strain collection (37). Of these 34, one had previously been investigated for IR tolerance and was found to have D10 values in line with classification as radiosensitive, and one, from a phylum without radiosensitive examples in the Train/Test, was experimentally validated (**Figure 2B**). Through this finding, we demonstrate that TolRad can be applied to bacterial genomes to which Pfam domains have already been assigned, including the upward of 46,000 bacterial proteomes currently on UniProt (https://www.uniprot.org/).

**Figure 2:**
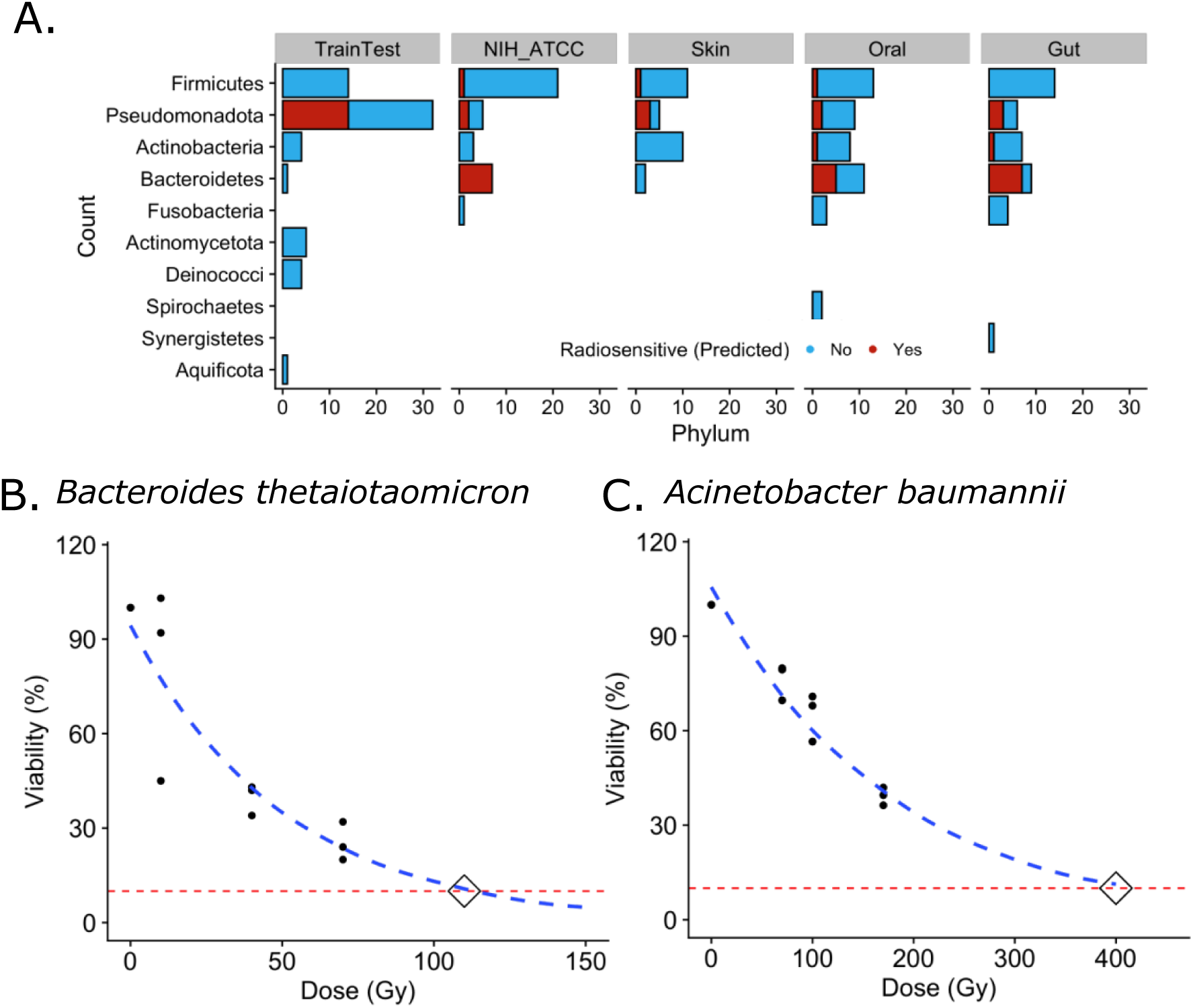
Apply TolRad to Bacteria Isolated from the Human Microbiome. A. Counts of bacteria species by phyla. Phylum diversity and tolerance predictions (by color) within the Train/Test Set, the ATCC-hosted NIH Human Microbiome Project, and additional species isolated from the Human Skin, Oral, and Gut Microbiome. B. Survival, determined via CFU, of *Bacteroides thetaiotaomicron* at 10, 40 and 70Gy. The D10 was determined to be 110Gy. Survival at 200Gy is predicted to be 1.8%. C. Survival, determined via CFU, of *Acinetobacter baumannii* at 10, 70, and 170Gy. The D10 was determined to be 400Gy.

### TolRad remains accurate when using *de novo* Pfam annotations assigned using EggNOG

We next sought to expand the utility of TolRad beyond pre-annotated UniProt Proteomes to Metagenome Assembled Genomes (MAGs). Since MAGs are constructed from community samples, they are unlikely to match with existing annotated genomes. For this reason, MAGs need to undergo both gene calling and Pfam annotation before TolRad can be applied. Additionally, MAGs, unlike UniProt Proteomes, are often only partial genome assembles (44).

To test the consistency of classifications made by TolRad on *de novo* Pfam annotations, the genome annotation pipeline EggNOG-Mapper (32) was used to assign coding regions and to annotate Pfam domains directly from the genome assemblies of the Train/Test Set. The workflow we used for processing and classifying MAGs is described in **Figure 3A**. TolRad returned the correct classification for 59 of the 61 (accuracy of 0.967%) EggNOG-mapper annotated genomes in the Train/Test Set (**Figure 3B, Sup. Table 3**).

**Figure 3:**
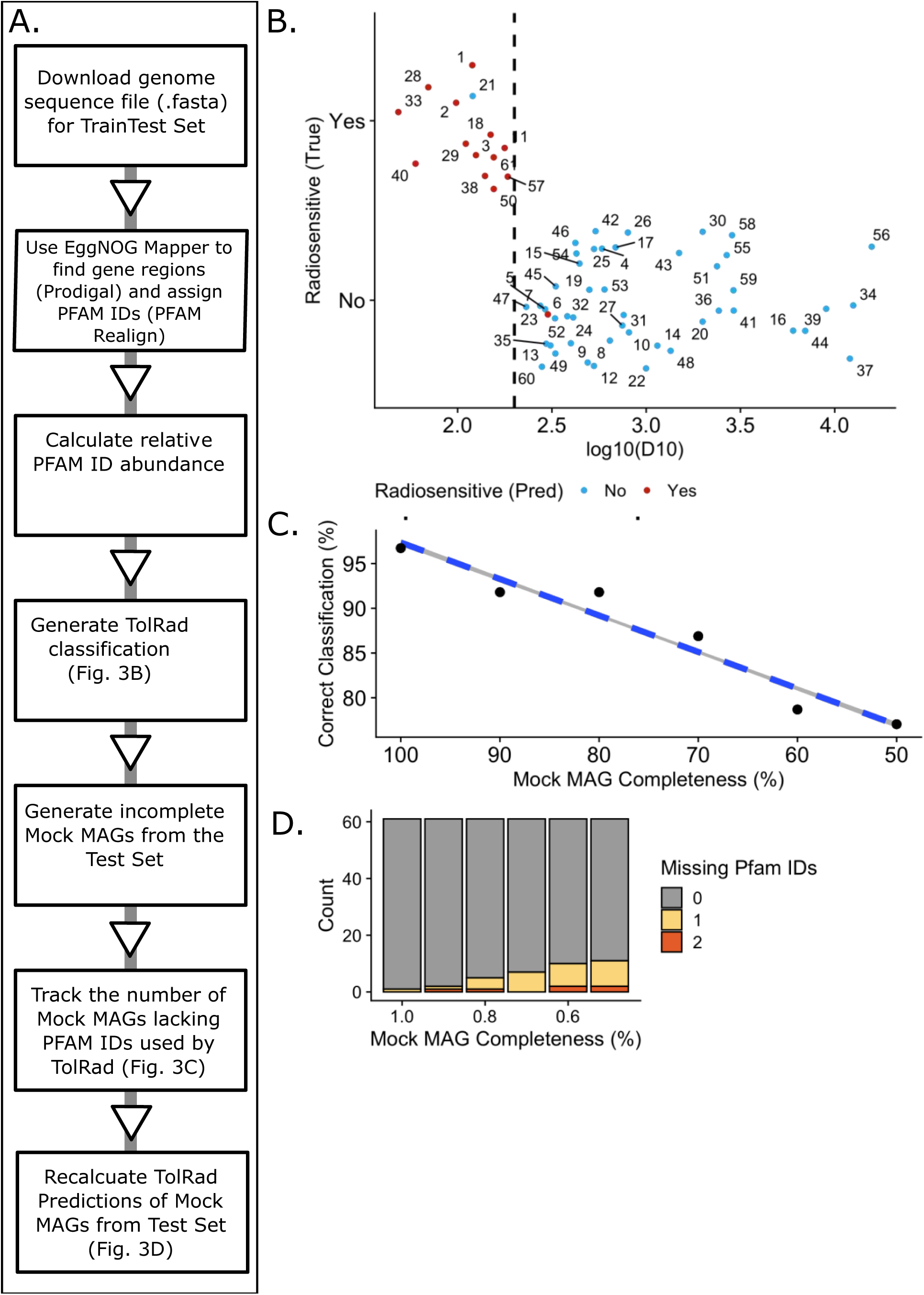
A. Workflow for generating TolRad predictions on *de novo* genome annotations. First, the genome sequence is downloaded. EggNOG Mapper is used to assign PFAMs. PFAM abundance is calculated. TolRad classifications are made. Then, PFAM domains were randomly removed, and the classification was determined again. B. *de novo* genome annotations were generated for the 61 species of the Train/Test Set. Species numbers (Graph ID) are in **Sup. Table 3**. These “mock MAGS” were classified using TolRad. Color denotes the classification assigned by TolRad. Red denotes a species classified as radiosensitive. Blue denotes a species classified as tolerant. The dashed line at 200Gy is the cut-off used for defining radiosensitive vs. tolerant species. C. The Mock MAGs were randomly degraded to 90, 80, 70, 60 and 50 %. The PFAM abundances were re-calculated along with the TolRad classification. The percent of correct classifications at each level is reported. D. The number of missing predictors at each level of degradation within the Mock MAGs from Fig.3C.

To test the ability of TolRad to handle incomplete genomes, a common occurrence in MAG datasets (44), we generated mock incomplete MAGs that ranged from 90-50% completion, in increments of 10%. This was done by randomly sampling the Pfam Domains generated from each EggNOG-mapper produced annotation files. After mock degradation, the frequencies of the predictor Pfam’s were recalculated, and the bacteria was reclassified by TolRad. The percent of the mock incomplete MAG’s correctly classified at each level of completeness were recorded. As shown in **Figure 3C**, the ability of TolRad to classify bacterial tolerance to IR worsened as the completeness of the mock MAGs decreased in a linear fashion; however, the classification rate stayed above 85% correct until 40% of the annotations had been removed. We also examined the relationship between mock incomplete MAGs and the rate at which predictor Pfam domains were lost (**Figure 3D**); however, even at 50% incomplete, less than 20% of the mock incomplete MAGs were missing a predictor Pfam domain. In summary, the use of Pfam domain frequency determined using the EggNOG-Mapper (32) genome annotation pipeline did not decrease the ability of TolRad to correctly classify bacterial tolerance for as IR and TolRad performance was reasonably robust on partial genomes.

### Applying TolRad to a collection of MAGs identified the deep sea and human gut as harboring a number of putative radiosensitive bacteria

To demonstrate the utility of TolRad for identifying radiosensitive species from within environmental samples and to gain insights into the ecological distribution of radiosensitive bacteria, we applied TolRad to three collections of previously assembled MAGs collected from diverse environments.

First, we examined a collection of MAGs originating from the human microbiome, representing the microbiome of the skin (HSM) and the gut (HGM) (45). The 91 high-quality MAGs (completeness > 60%) in this dataset had been previously assigned to four Phylum (*Firmicutes*, *Proteobacteria*, *Actinobacteriota*, and *Bacteroidota*) and had a mean completeness of 93.25%. Within this dataset, only 3 MAGs lacked one of the Pfam domains used by the model. TolRad characterized 7 (10.6%) as radiosensitive (**Table 4, Sup. Table 4**). All putative radiosensitive MAGs were members of either *Bacteroidota* or *Firmicutes* phylum (**Sup. Figure 2A**). This finding agreed with the radiosensitive predictions made on the HMB UniProt dataset (**Figure 2A**). Interestingly, all the MAGs predicted to be radiosensitive were collected from the HGM (17 out of 64, 26.56%), and no radiosensitive bacteria were predicted from the HSM-collected MAGs (**Figure 4A**). In the authors’ original analysis, the 91 MAGs were binned to 46 NCBI taxonomies, and 18 of these NCBI taxonomies included multiple MAGs. We expected that MAGs within the same NCBI Taxonomy would have the same tolerance for IR. TolRad had consistent tolerance predictions for 17 of 18 NCBI Taxonomies (**Sup. Figure 2B**).

**Figure 4:**
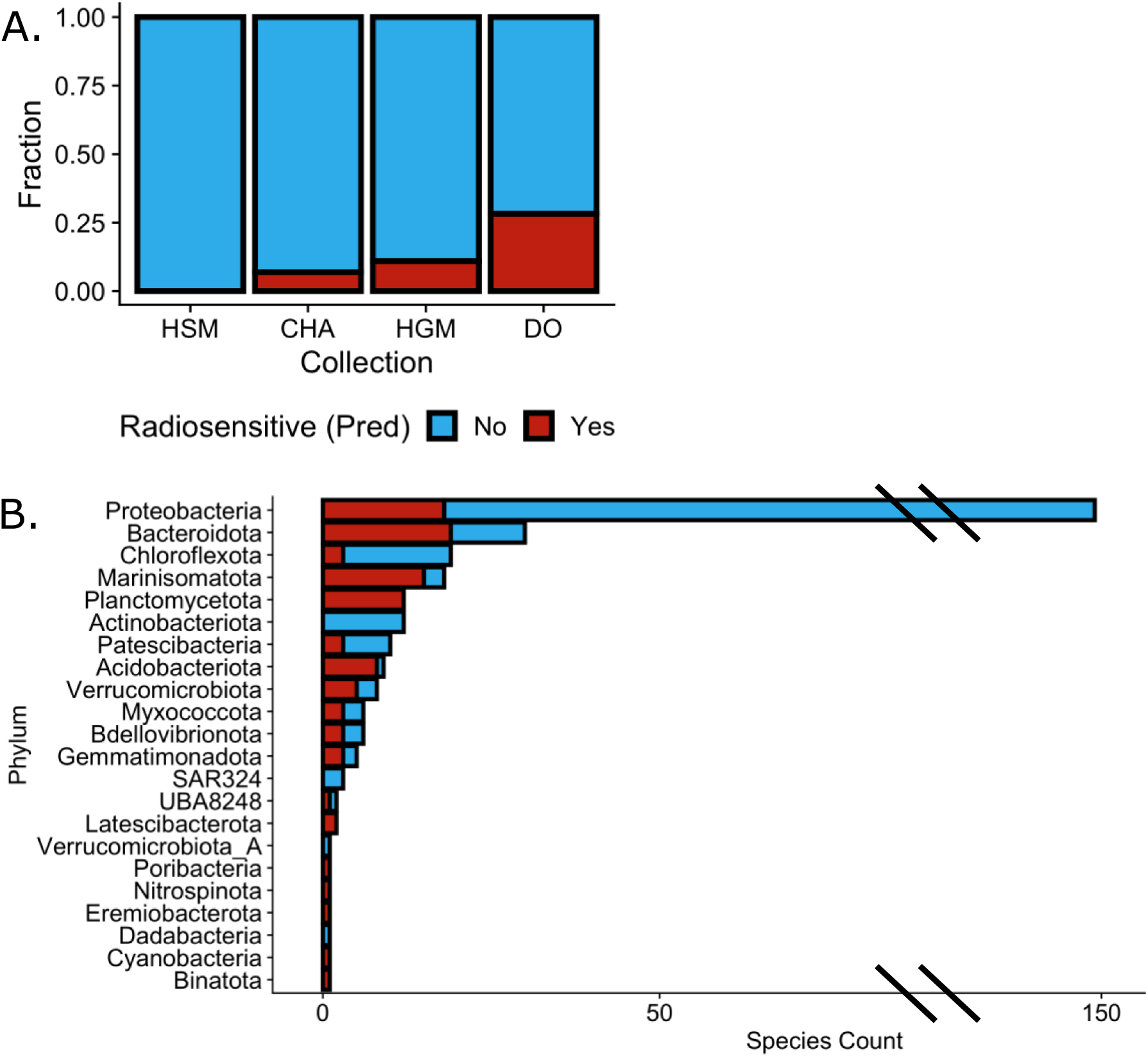
Application of TolRad to Three MAG Collections. A: As described in Figure 3, *de novo* annotations were assigned to previously published environmentally collected MAGs. The fraction of previously published MAGs classified as Radiosensitive (Red) or Tolerant (Blue) across MAGs collected from the Human Skin Microbiome (HSM), a glacial stream in the Canadian High Arctic (CHA), the Human Gut Microbiome (HGM) and the water column of the Deep Ocean (DO). B. Counts of MAGs, by phyla, from the Deep Ocean collection that were classified as Radiosensitive or Tolerant.

**Table 4:** MAG Summary.

| Table 4: MAG Summary |  |  |  |  |  |  |
| --- | --- | --- | --- | --- | --- | --- |
| Dataset | Total MAGs | Source | High Quality |  | Classification |  |
|  |  |  | MAGs | Phylum # | Radiosensitive | Tolerant |
| CHA | 31 | Trivedi et al. 2020 | 26 | 5 | 2 | 24 |
| Deep Ocean | 317 | Acinas et al. 2021 | 250 | 21 | 78 | 172 |
| HMB | 96 | Arikawa et al. 2021 | 91 | 4 | 7 | 84 |
| Total | 444 |  | 367 |  | 87 |  |

Next, we examined a set of MAGs collected from seasonal glacial surface streams in the Canadian High Arctic (CHA) (46). This MAG set represents an environment where, due to the high levels of solar radiation, we did not expect to identify radiosensitive bacteria (47). The High Arctic MAGs had a lower mean completeness (77.15%) compared with the other MAGs sets, although the number of MAGs lacking the predictor Pfam domains was minimal. As we expected, of the 26 MAGs with high-quality MAGs, only two (7.6%) were predicted to be radiosensitive and both were identified as *Proteobacteria* from the genus *Cellvibrio* and *Pseudomonas (***Sup. Figure 2C, Sup. Table 4**).

To discover a greater diversity of radiosensitive bacteria, we applied TolRad to a set of 312 MAGs collected from the Deep Ocean (33). The original publication assigned these MAGs to 26 phyla and the MAGs had a mean completeness of 84.2%. Due to the total lack of UV radiation in the deep sea, and the suspected role of ROS stresses, such as UV exposure, in the evolution of IR tolerance (25), we expected to identify a diversity of radiosensitive bacteria in this MAG dataset. Compared with the HMB dataset and the mock incomplete MAGs, the Deep Ocean collection had a greater number of MAGs that were missing predictor Pfam domains (**Sup. Table 3**) This finding may be due to the diversity of the Deep Ocean collection, which included many phyla not represented in the Train Set (**Figure 4B**). We observed that MAGs assigned to phyla not in the Train/Test set had a greater rate of missing predictor Pfam domains than the MAGs assigned to phylum in the Train/Test Set. Because we were unable to model the impact of missing predictor Pfam domains on TolRad accuracy, MAGs that were lacking 2 or more Pfam domains were excluded from classification. This left 250 MAGs from 21 Phylum. Of the MAGs examined, 78 were predicted to be radiosensitive (31.2%) (**Figure 4A, Sup. Table 4**), the highest percent of the MAG collections we examined. These MAGs came from 17 phyla, with many MAGs belonging to the *Acidobacteriota*, *Planctomycetota,* or *Marinisomatota* phyla (**Figure 4B**), suggesting that these phyla are ripe for further investigation as sources of IR biomarkers.

## Discussion

TolRad is a random forest binary classifier that uses protein annotations, which can be generated directly from the genome assembly, to predict if a bacteria is radiosensitive (D10 < 200Gy). TolRad is available as a stand-alone predictive tool and can be accessed at https://github.com/philipjsweet/TolRad and can be applied to both reference proteomes and MAGs. The ability of TolRad to identify radiosensitive bacteria species on which it had not been trained on, was demonstrated using a Test Set of bacteria (with known IR tolerances) that were excluded from the construction of TolRad (**Figure 1D**). The decrease in accuracy from the Train Set to Test Set was negligible (from 0.875 to 0.900). The ability of TolRad to make radiosensitive identifications for species from phylum without radiosensitive representation in the Train/Test Set was demonstrated by experimentally validating the radiosensitive classification of a *Bacteroidetes* species and tolerant classification of an *Acinetobacter* species (**Figure 2B**). The generalizability of TolRad was shown by applying TolRad to both pre-annotated proteomes from UniProt and *de novo* assembled proteomes to identify putative radiosensitive species from across 19 phyla. Additionally, TolRad can be used to understand the general IR tolerance of bacterial communities (**Figure 4A**). We also demonstrate that TolRad can be used to classify the tolerance of bacteria that have been well studied, such as those from the ATCC Human Microbiome Collection (37) (**Figure 2**), as well as Metagenome Assembled Genomes (MAGs), such as those previously assembled from the Deep Sea (46) (**Figure 4B**). Applying TolRad has allowed us to greatly expand the number and diversity of bacteria which are likely to have a low survival threshold for IR exposure. We have uploaded TolRad to GitHub allowing for integration into metagenomic analysis pipelines. In summary, we have created the first predictive model of bacterial tolerance for IR that relies exclusively on genomic annotations and demonstrated that this model can be widely deployed.

### Insights into the genetic traits of radiosensitivity

In addition to the construction of TolRad, this study also allows for a greater understanding of the bacterial traits that are associated with radiation tolerance. Previous work by the Daly Lab (23, 25, 26) has demonstrated that the intracellular ratio of Manganese to Iron (Mn/Fe) is a predictor of a species’ tolerance for IR. This correlation between radiation tolerance and the intracellular ratio of Mn/Fe was also observed in UV-C tolerant bacteria (48). While genetic explanations have been offered for the sensitivity of specific bacteria, such as a large number of proteins with heme (i.e. iron-binding) domains in the radiosensitive bacteria *S. oneidensis* (26) (49) a broadly applicable understanding of the genetic origins of radiation sensitivity has not been proposed. The construction of TolRad required the identification of Pfam domains informative of radiation tolerance; we present these findings, as well as the Mean Decrease in Accuracy and examples of *Escherichia coli* genes with these domains, in **Table 2**. We found that PF07992 *Pyridine nucleotide-disulphide oxidoreductase* domains are often found in reductases within known roles regulating intracellular ROS such as *norW, txrB* and *ndh*. The importance of PF07992 is in agreement with previous work demonstrating the importance of limiting ROS spread to ensure IR survival (26). PF00300 *Histidine phosphatase superfamily (branch 1)* was the second most important predictor in the model. These domains are found in a diverse set of proteins (**Table 2**), complicating its the interpretation of the contribution to IR tolerance. The contribution of the PF03466, *LysR substrate binding domain*, which is often found in ligand response transcription factors, is not immediately clear, however it has been previously noted that while the radiosensitive bacterium *S. oneidensis* has 52 proteins in the LysR Family, the radioresistant bacterium *D. radiodurans* only has two (50). Similarly, the connection of PF00849 *RNA pseudouridylate synthase*, a domain found only in RNA pseudouridylate synthase, to the classification of bacterial IR tolerance is unclear. A complication of interpreting random forest predictors is that the random forest model does not produce an equation, with coefficients that speak to the relationship between the predictor and the response variable. Future examinations of specific radiosensitive bacteria, such as those identified in this study, and focused studies into the proteins with the Pfam domains that we have correlated with IR tolerance will be required to fully understand the biological implications of these four Pfam domains for IR-tolerance.

### Applications of TolRad

There are several applications for TolRad, including providing context for metagenomic studies of bacterial communities after IR exposure (19), searching for bacterial sources of low-dose biomarkers (49), and as a guide for selecting against radiosensitive species for bioremediation development (17). Currently, the only way to determine the sensitivity of a bacteria species to IR is to conduct extensive experimentation, requiring both the ability to culture the bacteria of interest and a high-powered source of IR. While survival screens have enabled the isolation of highly radioresistant bacteria (30, 51) (52) no such discovery studies have been conducted for radiosensitive bacteria. As an example of applying TolRad to screen existing proteomes, we applied TolRad to a collection of 152 proteomes downloaded from UniPro, including the ATCC Human Microbiome strain collection (**Sup. Table 4**). TolRad identified 34 putative radiosensitive species, including several of the most abundant bacteria in the human gut. We then demonstrated experimentally that *B. thetaiotaomicron* is in fact radiosensitive, with a D10 of 110Gy. Excitingly, these predictions were correct, despite being made on species from phylum on which TolRad was not trained (**Figure 2C**).

As another example of applying TolRad for screening bacterial communities for radiosensitive species, we applied TolRad to MAGs to a collection of Deep Sea MAGs, and identified 78 candidate species, from 17 phyla as radiosensitive (**Figure 4B**). TolRad will allow other researchers to scan bacterial genomes rapidly and to determine which species from, a population is radiosensitive. These predictions will help guide future exploration of bacterial tolerance of IR.

### Taxonomy of Radiosensitive Species

We were also interested in the taxology of radiation tolerance, as when we began this study, bacteria with a low survival threshold for IR exposure had only been identified within the *Proteobacterium* phylum. To our knowledge, the dataset that we collected for TolRad is the most comprehensive set of bacterial D10 values published to date. We observed, within the Train/Test set, that intolerance for IR was limited to Proteobacteria; however, when we expanded our search to bacteria of unknown IR tolerances using TolRad, we found a diversity of putative radiosensitive. Across the UniProt proteomes and the three collections of MAGs, we applied TolRad to over 500 proteomes, representing 44 phyla, and identified 121 putative radiosensitive species across 21 phyla (**Sup. Table 3**). Since there is a minimal amount of naturally occurring IR on Earth, previous studies have suggested extreme radiation tolerance, observed in multiple phyla, (**Figure 1B** and **Sup. Table 1**), may have evolved as a response to bacteria living in environments with elevated ROS stress (31). Previous work with marine bacterial noted that that species collected from the subsurface had a lower tolerance for UV radiation and hydrogen peroxide, both ROS inducing stresses, than those collected at the ocean surface. The authors suggest that this may be due to a lower selective pressure from sunlight exposure on the subsurface species (47). This dynamic could also explain the difference in radiosensitive bacteria between the HSM and the HGM (**Figure 4A**). This idea is supported by the low number of radiosensitive bacteria that TolRad classified from the high UV (CHA) and high hydrogen peroxide (HSM) MAGs. Based on the predictions made by TolRad, we suggest that, in a similar way, IR intolerance could be prevalent in UV-sheltered environments, such as the deep sea (**Figure 4A**) and the human gut (**Figure 4A**). Identifying environments that are rich in radiosensitive species can aid in understanding the biological causes of IR intolerance.

## Limitations

The greatest limitation of TolRad is the limited size of the Test/Train Set. We were only able to collect D10 values from control (RT and in growth media or PBS) conditions for 61 bacteria (**Sup. Table 1**). These bacteria only represented 5 bacterial phyla, and radiosensitive bacteria were only found within one of those phyla. The Train/Test set was heavily skewed toward *Proteobacteria* (41.3%) and *Firmicutes* (30.0%) so it is possible that the model is more accurate on these phyla than on others. The experimental validation that we conducted on two representatives of *Bacteroidetes*, supports the use of TolRad beyond the phylum of the Train/Test set; however, similar initial validation experiments would be prudent for radiation intolerance predictions made on species from additional phylum. To build TolRad, only Pfam domains present in all the test set bacteria were used, and this selection worked well for the reference bacteria, the HMB MAGs, and the High Canadian Artic MAGs, all of which had a similar range of phyla as the Train Set. When examining the Deep Ocean MAG collection, we noted that some of the phyla had a much higher rate of proteomes that were missing predictor Pfam domains (**Sup. Table 4**). TolRad reports the number of missing predictor Pfam domains for each genome that is classified, and we only discuss classifications of putative radiosensitive species with at least three of the four predictors. Undoubtedly, TolRad will make misclassifications; however, the ease of incorporating additional D10 data (as it becomes available) into TolRad means that future findings can be incorporated into TolRad will serve to bolster the predictive power of this tool.

In summary, in this paper, we have described TolRad, the first strictly computationally based classifier of bacterial tolerance for IR. We have demonstrated the accuracy of TolRad beyond the bacteria species on which it was trained. We further demonstrate the ability of TolRad to identify novel bacteria with a low survival threshold for IR exposure. Additionally, we present the first experimental characterization of the tolerance of a *B. thetaiotaomicron*, an abundant member of the human microbiome. Further collection and validation of radiation phenotypic straits in bacteria should provide additional data upon which to continue to improve TolRad and other similar training models.

## Methods

### Collection of D10 Values: Supplemental Table 1

The species used in the Train/Test Set are presented in **Supplemental Table 1** and **Supplemental Table 2**. Only D10 values determined at room temperature and in a liquid matrix were considered for inclusion. For bacteria with multiple reported D10 values the mean was used to assign a tolerance classification to the bacteria. The UniProt Proteome from which Pfam domains were obtained is also provided. The lowest 20% of the D10 values were classified as radiosensitive. Bacteria were classified as Radiosensitive if the mean D10 was below 200Gy and as Tolerant if the mean D10 was above 200Gy.

### Selection of Pfam Domains and models construction

The Pfam domains for each bacteria in the Train/Test Set were acquired from UniProt. When possible, the “Reference” version of the proteome was used. Otherwise, the most complete proteome was used. No proteomes classified by UniProt as “low coverage” were used in the Train/Test Set. The UniProt proteome IDs are reported in **Supplemental Table 1**. The Train/Test Set was randomly split 70/30 into a Train Set (n=40) and a Test Set (n=21). The Train Set was used for the construction of TolRad. The raw counts of each Pfam Domain were calculated at the species level and across the Train Set. As shown in **Figure 1B**, we started with 7409 unique Pfam domains. We then selected domains that were universal to the Train Set, which left 395. Of these, we selected those with a mean occurrence per species greater than 2, this left 132. For each species, the relative frequency of each Pfam domain, against the total number of Pfam domains within each species, was calculated. The Boruta (35) feature selection algorithm (maxRuns = 500) was used to select 7 Pfam domains for which the frequency (out of the total number of Pfam domains for the species) was correlated with the species IR tolerance classification. The relative frequency of these domains was used to construct a random forest model using the RandomForest algorithm of the caret R library (https://cran.r-project.org/web/packages/caret/index.html) to bin bacteria as tolerant of IR or radiosensitive of IR using the relative frequency of each of the predictor Pfam domains. A 10X cross-validation was used to train the model. The Mean Decrease in Accuracy for each of the 7 predictor Pfam domains was determined and predictors were removed stepwise, starting with the domain with the lowest Mean Decrease in Accuracy. Of the 7 domains, 4 were determined to be required for correct classification and were retained for the final model (**Table 2**).

### Making Predictions using TolRad UniProt

For the classification of HMB UniProt species, Pfam annotations were downloaded from UniProt. In **Supplemental Table 4**, UniProt Species names along with the species name the classification, the total number of missing Pfam IDs and taxonomy as well as the source suggesting that the species is in the HMB.

### Processing MAGs

MAGs were downloaded preassembled, and the authors previously assigned taxonomy terms were used. The Train/Test Genome assemblies associated with the UniProt proteomes used to Train and Test TolRad were downloaded from EMBLE (https://www.ebi.ac.uk/CHA MAGs were downloaded from). The assembly IDs are in **Supplemental Table 4**. MAGs from the CHA (46) were downloaded, along with taxonomy assignments and completeness scores from https://figshare.com/articles/dataset/Borup_Fiord_Pass_-_Metagenome_Assembled_Genomes_MAGs_/9767564. Human Microbiome (45) samples were downloaded from SRA (https://www.ncbi.nlm.nih.gov/bioproject/) and taxonomy assignments and completeness scores were acquired from the supplemental figures. The Genome assembles Test/Train Set and the MAGs HMB and the CHA were processed using EggNOG-Mapper (32) (--pfam_realign realign --itype genome --genepred prodigal). The Deep Sea MAGs (33) were downloaded, with Pfam annotations, from https://malaspina-public.gitlab.io/malaspina-deep-ocean-microbiome/. Pfam domain frequency and TolRad predictions were generated as described above.

### Bacterial Strains and culture conditions

*Bacteroides thetaiotaomicron* was in grown Brain Heart Infusion Media (BHIM) with anaerobic conditions without shaking at 37°C. For all experiments, unless otherwise stated, cells were grown from single colonies in 10ml of BHIM in Hungate tubes. The morning of an exposure, 1.0ml of O/N culture was diluted into 9.0ml of fresh media to an OD_600_ of ∼0.1 in a Hungate Tube. Cells were grown for ∼5h until an OD_600_ of ∼0.4. All strains used are provided in Table 5.

**Table 5:**
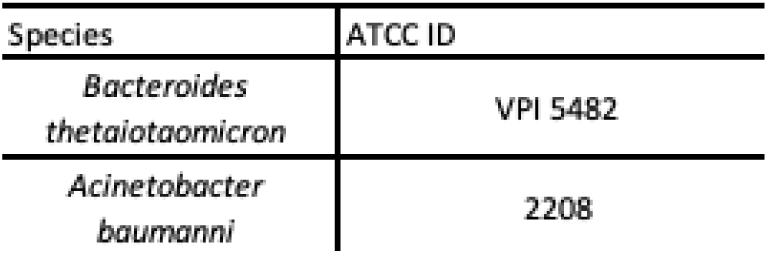
Strains used in this Study.

*Acinetobacter baumanni* was grown in LB media at 37 with shaking. For all experiments, unless otherwise stated, cells were grown from single colonies in 5ml of LB and grown on LB plates.

### X-Ray Exposures

When exposing cells to IR a Faxitron 225 MuliRad X-ray irradiator was used. The machine was set to 12mA, 220kV to deliver ∼5Gy/min. A 0.5mm aluminum filter ensured that only high-energy particles were delivered. Before exposure, cells were grown to exponential phase (OD_600_ of 0.6). 5ml of culture was sealed in de-gassed Nasco Whirl-Pak bags, sealed in anaerobic growth bags, and laid flat on the exposure tray to ensure even dosage across the sample. Exposures were conducted at room temperature (24°C).

### CFU Assays

For survival assays, three biological replicates were grown to mid-exponential phase, bagged, and exposed as described above. After exposure, cells were serially diluted and spread on BHIM or LB plates using beads. Three technical replicates were conducted per biological replicate. CFU were counted and the mean Sham plate count of the three technical replicates was used as the baseline against which the technical replicates of the doses of that biological replicate were compared to determine the surviving fraction. The significance of differences was determined using a t-test between the six survival fractions per dose against the sham, and the six survival fractions of the sham control.

## Acknowledgments

We would like to thank Daryl Barth (University of Texas at Austin) for assistance in running the EggNog-Mapper annotations.

This research was funded by a grant (FA9550-20-1-0131) from the Air Force Office of Scientific Research as well as a grant (HDTRA1-17-1-0025) from the Defense Threat Reduction Agency.

This work was also supported by a grant (W911NF22S0002) from the Intelligence Advanced Research Projects Activity.

---

**Supplemental Figure 1:**
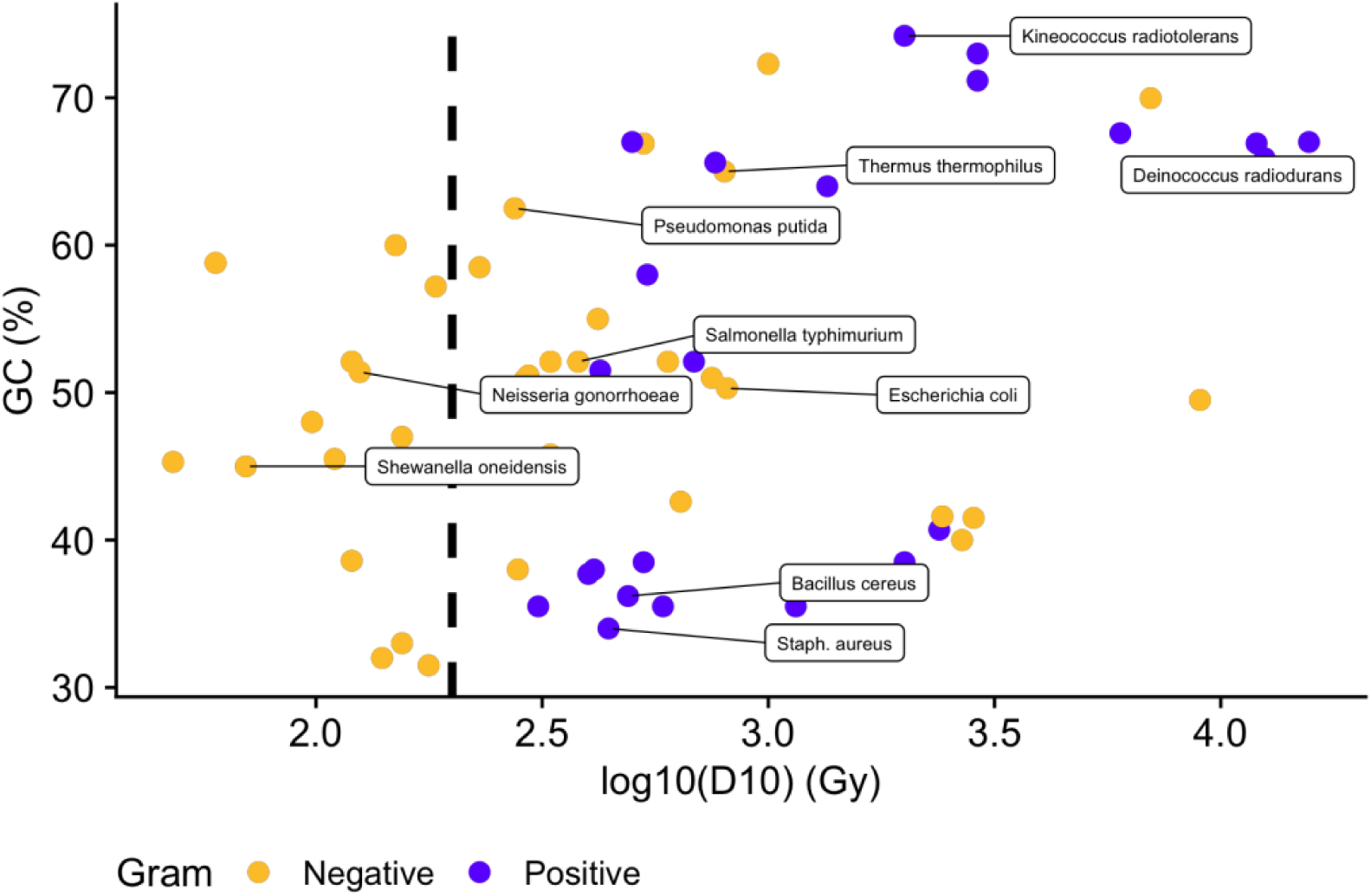
Traits of the Test/Train Set. Each point is a species D10 value. Species farther to the right have a higher D10. Species higher up have a higher GC content. Color denotes gram stain, purple dots are gram (+) species, and yellow dots are gram (-) species. All data is also found in Sup. Table 1. Species of interest are labeled. The dashed line denotes the 200Gy cutoff for radiosensitive species.

**Supplemental Figure 2:**
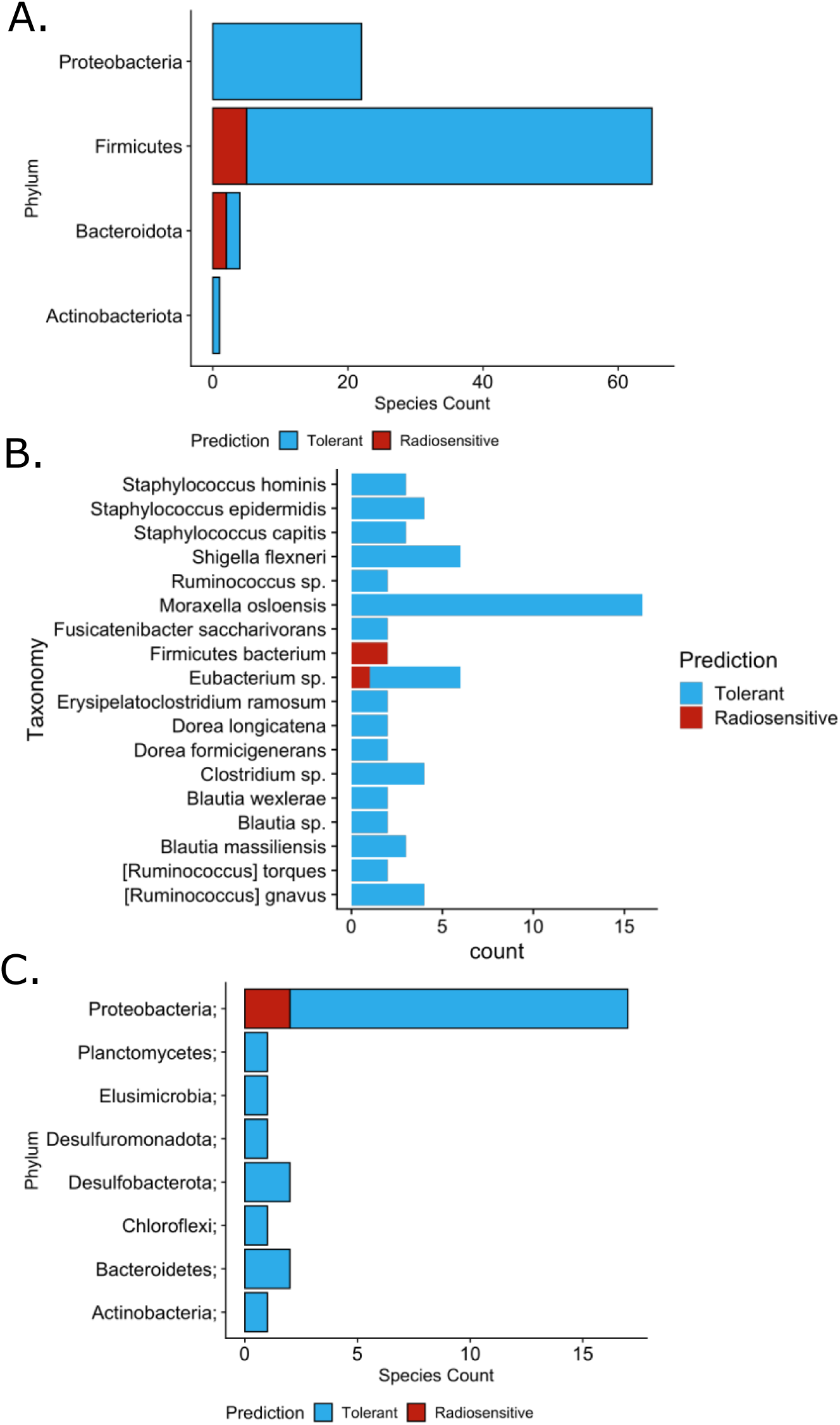
Taxonomy of MAGs. A. Counts of MAGs from the Human Microbiome by phyla. Color denotes if the MAG was identified as radiosensitive (Red) or Tolerant (Blue). B. Tolerance classifications for MAGs from the HMB collection belonging to the same NCBI Taxonomy. Color denotes TolRad predicted tolerance. C. Same as A, but for the Canadian High Arctic MAGs.

