## Supplementary material for "TolRad: A model for predicting radiation tolerance using Pfam annotations identifies novel radiosensitive bacterial species from reference genomes and MAGs": Sup. Tables

**Sup. Table 1: Mean D10 for all species in The Train/Test set with the UniProt straina and Genome fi**

| Species | Mean D10 | log10(D10) | Radiosensitive |
|---|---|---|---|
| Staphylococcus aureus | 443.3333333 | 2.646730386 | No |
| Lactococcus lactis | 585 | 2.767155866 | No |
| Staphylococcus epidermidis | 310 | 2.491361694 | No |
| Lactobacillus acidophilus | 1150 | 3.06069784 | No |
| Bacillus cereus | 489.6 | 2.689841409 | No |
| Bacillus sphaericus | 1500 | 3.176091259 | No |
| Listeria monocytogenes | 411.75 | 2.614633608 | No |
| Streptococcus thermophilus | 530 | 2.72427587 | No |
| Enterococcus faecium | 2000 | 3.301029996 | No |
| Enterococcus faecalis | 2680 | 3.428134794 | No |
| Bacillus pumilus | 2386.666667 | 3.377791768 | No |
| Coxiella burnetii | 640 | 2.806179974 | No |
| Escherichia coli | 810 | 2.908485019 | No |
| Shigella flexneri | 290 | 2.462397998 | No |
| Shigella sonnei | 750 | 2.875061263 | No |
| Morganella morganii | 295 | 2.469822016 | No |
| Salmonella enterica | 425 | 2.62838893 | No |
| Bifidobacterium breve | 540 | 2.73239376 | No |
| Aeromonas salmonicida | 230 | 2.361727836 | No |
| Pseudomonas putida | 275 | 2.439332694 | No |
| Rhodococcus erythropolis | 1350 | 3.130333768 | No |
| Thermus thermophilus | 800 | 2.903089987 | No |
| Mycobacterium tuberculosis | 763.3333333 | 2.882714228 | No |
| Deinococcus geothermalis | 12500 | 4.096910013 | No |
| Stenotrophomonas maltophilia | 530 | 2.72427587 | No |
| Rubrobacter radiotolerans | 12000 | 4.079181246 | No |
| Deinococcus radiodurans | 15666.66667 | 4.194976603 | No |
| Mycobacterium smegmatis | 500 | 2.698970004 | No |
| Rubrobacter xylanophilus | 6000 | 3.77815125 | No |
| Kocuria rhizophila | 2900 | 3.462397998 | No |
| Methylobacterium radiotoleran | 1000 | 3 | No |
| Micrococcus luteus | 2900 | 3.462397998 | No |
| Kineococcus radiotolerans | 2000 | 3.301029996 | No |
| Salmonella typhimurium | 380 | 2.579783597 | No |
| Pediococcus pentosaceus | 685 | 2.835690571 | No |
| Priestia megaterium | 400 | 2.602059991 | No |
| Acinetobacter radioresistens | 2425 | 3.384711743 | No |
| Spirosoma radiotolerans | 9000 | 3.954242509 | No |
| Deinococcus ficus | 7000 | 3.84509804 | No |
| Paenibacillus amylolyticus | 330 | 2.51851394 | No |
| Enterobacter sp. BIGb0383 | 420 | 2.62324929 | No |

| | | | |
|---|---|---|---|
| Salmonella enterica subsp. enterica s | 330 | 2.51851394 | No |
| Aquifex pyrophilus | 2842 | 3.453624074 | No |
| Proteus vulgaris | 279.5 | 2.446381812 | No |
| Salmonella muenster | 600 | 2.77815125 | No |
| Salmonella Senftenberg | 330 | 2.51851394 | No |
| Shigella boydii | 300 | 2.477121255 | No |
| Campylobacter jejuni | 177.5 | 2.249198357 | Yes |
| Campylobacter lari | 140 | 2.146128036 | Yes |
| Campylobacter coli | 155 | 2.190331698 | Yes |
| Shewanella oneidensis | 70 | 1.84509804 | Yes |
| Shewanella putrefaciens | 110 | 2.041392685 | Yes |
| Vibrio cholera O1 | 98 | 1.991226076 | Yes |
| Neisseria gonorrhoeae | 125 | 2.096910013 | Yes |
| Serratia marcescens | 60 | 1.77815125 | Yes |
| Aeromonas hydrophila | 150 | 2.176091259 | Yes |
| Acinetobacter calcaeceticus | 120 | 2.079181246 | Yes |
| Salmonella paratyphi | 120 | 2.079181246 | Yes |
| Yersinia enterocolitica | 155 | 2.190331698 | Yes |
| Klebsiella variicola | 184 | 2.264817823 | Yes |
| Vibrio parahaemolyticus | 48.33333333 | 1.684246747 | Yes |

**le, taxonomy data, and traits**

| UniProt Strain |
|---|
| Staphylococcus aureus (strain NCTC 8325 / PS 47) |
| Lactococcus lactis subsp. lactis (strain IL1403) (Streptococcus lactis) |
| Staphylococcus epidermidis (strain ATCC 35984 / RP62A) |
| Lactobacillus acidophilus (strain ATCC 700396 / NCK56 / N2 / NCFM) |
| Bacillus cereus (strain ATCC 14579 / DSM 31 / CCUG 7414 / JCM 2152 / NBRC 15305 |
| Lysinibacillus sphaericus (Bacillus sphaericus) |
| Listeria monocytogenes serovar 1/2a (strain ATCC BAA-679 / EGD-e) |
| Streptococcus thermophilus (strain ATCC BAA-250 / LMG 18311) |
| Enterococcus faecium (strain ATCC BAA-472 / TX0016 / DO) |
| Enterococcus faecalis (Streptococcus faecalis) |
| Bacillus pumilus (strain SAFR-032) |
| Coxiella burnetii (strain RSA 493 / Nine Mile phase I) |
| Escherichia coli O157:H7 |
| Shigella flexneri |
| Shigella sonnei (strain Ss046) |
| Morganella morganii subsp. morganii KT |
| Salmonella enteritidis |
| Bifidobacterium breve MCC 1340 |
| Aeromonas salmonicida |
| Pseudomonas putida (strain ATCC 47054 / DSM 6125 / CFBP 8728 / NCIMB 11950 / |
| Rhodococcus erythropolis (Arthrobacter picolinophilus) |
| Thermus thermophilus (strain ATCC 27634 / DSM 579 / HB8) |
| Mycobacterium tuberculosis (strain ATCC 25618 / H37Rv) |
| Deinococcus geothermalis (strain DSM 11300 / AG-3a) |
| Stenotrophomonas maltophilia (strain K279a) |
| Rubrobacter radiotolerans |
| Deinococcus radiodurans (strain ATCC 13939 / DSM 20539 / JCM 16871 / LMG 4051 |
| Mycolicibacterium smegmatis (strain ATCC 700084 / mc(2)155) (Mycobacterium sm |
| Rubrobacter xylanophilus (strain DSM 9941 / NBRC 16129 / PRD-1) |
| Kocuria rhizophila |
| Methylobacterium radiotolerans (strain ATCC 27329 / DSM 1819 / JCM 2831 / NBR( |
| Micrococcus luteus Mu201 |
| Kineococcus radiotolerans (strain ATCC BAA-149 / DSM 14245 / SRS30216) |
| Salmonella typhimurium (strain LT2 / SGSC1412 / ATCC 700720) |
| Pediococcus pentosaceus (strain ATCC 25745 / CCUG 21536 / LMG 10740 / 183-1w) |
| Priestia megaterium (strain ATCC 12872 / QMB1551) (Bacillus megaterium) |
| Acinetobacter radioresistens DSM 6976 = NBRC 102413 = CIP 103788 |
| Spirosoma radiotolerans |
| Deinococcus ficus |
| Paenibacillus amylolyticus |
| Enterobacter sp. BIGb0383 |

Salmonella enterica subsp. enterica serovar Heidelberg
Aquifex aeolicus (strain VF5)
Proteus vulgaris
Salmonella muenster
Salmonella senftenberg
Shigella boydii serotype 18 (strain CDC 3083-94 / BS512)
Campylobacter jejuni subsp. jejuni serotype O:2 (strain ATCC 700819 / NCTC 11168)
Campylobacter lari NCTC 11845
Campylobacter coli
Shewanella oneidensis (strain MR-1)
Shewanella putrefaciens (strain 200)
Vibrio cholerae serotype O1 (strain ATCC 39315 / El Tor Inaba N16961)
Neisseria gonorrhoeae (strain ATCC 700825 / FA 1090)
Serratia marcescens
Aeromonas hydrophila subsp. hydrophila (strain ATCC 7966 / DSM 30187 / BCRC 13
Acinetobacter calcoaceticus ANC 3680
Salmonella paratyphi C (strain RKS4594)
Yersinia enterocolitica subsp. enterocolitica
Klebsiella variicola
Vibrio parahaemolyticus serotype O3:K6 (strain RIMD 2210633)

| Genome | Domain | Phylum | Class |
|---|---|---|---|
| GCA_000013425.1 | Bacteria | Bacillota | Bacilli |
| GCA_000006865.1 | Bacteria | Bacillota | Bacilli |
| GCA_000011925.1 | Bacteria | Bacillota | Bacilli |
| GCA_000011985.1 | Bacteria | Bacillota | Bacilli |
| GCA_000007825.1 | Bacteria | Bacillota | Bacilli |
| GCA_002923795.1 | Bacteria | Bacillota | Bacilli |
| GCA_000196035.1 | Bacteria | Bacillota | Bacilli |
| GCA_000011825.1 | Bacteria | Bacillota | Bacilli |
| GCA_000174395.2 | Bacteria | Bacillota | Bacilli |
| GCA_009498155.3 | Bacteria | Bacillota | Bacilli |
| GCA_007679755.1 | Bacteria | Bacillota | Bacilli |
| GCA_000007765.2 | Bacteria | Pseudomonadota | Gammaproteobacteria |
| GCA_000008865.2 | Bacteria | Pseudomonadota | Enterobacterales |
| GCA_000006925.2 | Bacteria | Pseudomonadota | Gammaproteobacteria |
| GCA_000092525.1 | Bacteria | Pseudomonadota | Gammaproteobacteria |
| GCA_000286435.2 | Bacteria | Pseudomonadota | Gammaproteobacteria |
| GCA_008834165.1 | Bacteria | Pseudomonadota | Gammaproteobacteria |
| GCA_001264105.1 | Bacteria | Actinomycetota | Actinomycetia |
| GCA_003947395.1 | Bacteria | Pseudomonadota | Gammaproteobacteria |
| GCA_000007565.2 | Bacteria | Pseudomonadota | Pseudomonadales |
| GCA_003990875.1 | Bacteria | Actinobacteria | Rubrobacteria |
| GCA_000091545.1 | Bacteria | Deinococci | |
| GCA_000195955.2 | Bacteria | Actinobacteria | Actinomycetia |
| GCA_000196275.1 | Bacteria | Deinococci | |
| GCA_000072485.1 | Bacteria | Pseudomonadota | Xanthomonadales |
| GCA_000661895.1 | Bacteria | Actinobacteria | Rubrobacteria |
| GCA_011604825.1 | Bacteria | Deinococci | |
| GCA_000015005.1 | Bacteria | Actinomycetota | Actinomycetia |
| GCA_000014185.1 | Bacteria | Actinobacteria | Actinomycetia |
| GCA_001190985.2 | Bacteria | Actinomycetota | Actinomycetia |
| GCA_000019725.1 | Bacteria | Pseudomonadota | Hyphomicrobiales |
| GCA_900163885.1 | Bacteria | Actinomycetota | Actinomycetia |
| GCA_000017305.1 | Bacteria | Actinomycetota | Actinomycetia |
| GCA_000006945.2 | Bacteria | Pseudomonadota | Gammaproteobacteria |
| GCA_000014505.1 | Bacteria | Bacillota | Bacilli |
| GCA_000025825.1 | Bacteria | Bacillota | Bacilli |
| GCA_000368905.1 | Bacteria | Pseudomonadota | Gammaproteobacteria |
| GCA_000974425.1 | Bacteria | Bacteroidetes | Bacteroidetes |
| GCA_003444775.1 | Bacteria | Deinococci | |
| GCA_003719335.1 | Bacteria | Bacillota | Bacilli |
| GCA_003751845.1 | Bacteria | Pseudomonadota | Enterobacterales |

| | | | |
|---|---|---|---|
| GCA_008124065.1 | Bacteria | Pseudomonadota | Gammaproteobacteria |
| GCA_015661675.1 | Bacteria | Aquificota | Aquificales |
| GCA_900455275.1 | Bacteria | Pseudomonadota | Gammaproteobacteria |
| GCA_008480455.1 | Bacteria | Pseudomonadota | Gammaproteobacteria |
| GCA_004768585.1 | Bacteria | Pseudomonadota | Gammaproteobacteria |
| GCA_000020185.1 | Bacteria | Pseudomonadota | Gammaproteobacteria |
| GCA_000009085.1 | Bacteria | Pseudomonadota | Epsilonproteobacteria |
| GCA_000816365.1 | Bacteria | Pseudomonadota | Epsilonproteobacteria |
| GCA_004956825.1 | Bacteria | Pseudomonadota | Epsilonproteobacteria |
| GCA_000146165.2 | Bacteria | Pseudomonadota | Gammaproteobacteria |
| GCA_000169215.1 | Bacteria | Pseudomonadota | Gammaproteobacteria |
| GCA_000006745.1 | Bacteria | Pseudomonadota | Gammaproteobacteria |
| GCA_000006845.1 | Bacteria | Pseudomonadota | Betaproteobacteria |
| GCA_001051865.1 | Bacteria | Pseudomonadota | Gammaproteobacteria |
| GCA_000014805.1 | Bacteria | Pseudomonadota | Gammaproteobacteria |
| APQH01 | Bacteria | Pseudomonadota | Gammaproteobacteria |
| GCA_000018385.1 | Bacteria | Pseudomonadota | Gammaproteobacteria |
| GCA_901472495.1 | Bacteria | Pseudomonadota | Gammaproteobacteria |
| GCA_014489395.1 | Bacteria | Pseudomonadota | Gammaproteobacteria |
| GCA_000196095.1 | Bacteria | Pseudomonadota | Gammaproteobacteria |

| Gram | OxygenUse | GC | Temp_Opt | Temp_max |
|---|---|---|---|---|
| Gram-positive | Facultative Anaerobic | 34 | 33.5 | 45 |
| Gram-positive | Facultative Anaerobic | 35.5 | 30 | 39 |
| Gram-positive | Facultative Anaerobic | 35.5 | 33.5 | 45 |
| Gram-positive | Facultative Anaerobic | 35.5 | 37 | 45 |
| Gram-positive | Facultative Anaerobic | 36.2 | 37 | 42.5 |
| Gram-positive | Anaerobic | 37.2 | 37 | 45 |
| Gram-positive | Facultative Anaerobic | 38 | 33.54 | 45 |
| Gram-positive | Facultative Anaerobic | 38.5 | 37 | 65 |
| Gram-positive | Facultative Anaerobic | 38.5 | 36 | 60 |
| Gram-negative | Facultative Anaerobic | 40 | 35 | 60 |
| Gram-positive | Anaerobic | 40.7 | 37 | 50 |
| Gram-negative | Anaerobic | 42.6 | 37 | |
| Gram-negative | Facultative Anaerobic | 50.3 | 37 | 45 |
| Gram-negative | Facultative Anaerobic | 50.9 | 37 | 45 |
| Gram-negative | Facultative Anaerobic | 51 | 37 | 45 |
| Gram-negative | Facultative Anaerobic | 51.15 | 35 | |
| Gram-positive | Facultative Anaerobic | 51.5 | 37 | 46 |
| Gram-positive | Anaerobic | 58 | 39 | 44 |
| Gram-negative | Facultative Anaerobic | 58.5 | 25 | 34.5 |
| Gram-negative | Aerobic | 62.5 | 30 | 35 |
| Gram-positive | Aerobic | 64 | 30 | 40 |
| Gram-negative | Aerobic | 65 | 65 | 81 |
| Gram-positive | Facultative Anaerobic | 65.6 | 37 | 45 |
| Gram-positive | Aerobic | 65.9 | 30 | 55 |
| Gram-negative | Aerobic | 66.9 | 35 | 37 |
| Gram-positive | Aerobic | 66.91 | 45 | 55 |
| Gram-positive | Aerobic | 67 | 30 | 37 |
| Gram-positive | Aerobic | 67 | 37 | 45 |
| Gram-positive | Aerobic | 67.6 | 60 | 70 |
| Gram-positive | Aerobic | 71.16 | 28 | 40 |
| Gram-negative | Aerobic | 72.3 | 27.5 | |
| Gram-positive | Aerobic | 73 | 31 | 45 |
| Gram-positive | Facultative Anaerobic | 74.2 | 41 | |
| Gram-negative | Aerobic | 52.1 | 37 | 45 |
| Gram-positive | Facultative Anaerobic | 52.1 | 30 | 42 |
| Gram-positive | Facultative Anaerobic | 37.7 | 37 | 48 |
| Gram-negative | Aerobic | 41.6 | 37 | 42 |
| Gram-negative | Aerobic | 49.5 | 28 | |
| Gram-negative | Aerobic | 69.97 | 37 | 40 |
| Gram-negative | Aerobic | 45.8 | 30 | 40 |
| Gram-negative | Facultative Anaerobic | 55 | 37 | 45 |

| | | | | |
|---|---|---|---|---|
| Gram-negative | Facultative Anaerobic | 52.1 | 37 | |
| Gram-negative | Facultative Anaerobic | 41.5 | 85 | 95 |
| Gram-negative | Facultative Anaerobic | 38 | 25 | 43 |
| Gram-negative | Facultative Anaerobic | 52.1 | 37 | |
| Gram-negative | Facultative Anaerobic | 52.1 | 37 | 50 |
| Gram-negative | Facultative Anaerobic | 50.6 | 37 | 47 |
| Gram-negative | Microaerophile | 31.5 | 37 | 45 |
| Gram-negative | Microaerophile | 32 | 42 | 42 |
| Gram-negative | Microaerophile | 33 | 37 | 42 |
| Gram-negative | Facultative Anaerobic | 45 | 30 | 40 |
| Gram-negative | Facultative Anaerobic | 45.5 | 32.5 | 40 |
| Gram-negative | Facultative Anaerobic | 48 | | 40 |
| Gram-negative | Aerobic | 51.4 | 36 | |
| Gram-negative | Facultative Anaerobic | 58.8 | 37 | 40 |
| Gram-negative | Facultative Anaerobic | 60 | 29.5 | 40 |
| Gram-negative | Aerobic | 38.6 | 35 | 45 |
| Gram-negative | Facultative Anaerobic | 52.1 | 37 | 42 |
| Gram-negative | Facultative Anaerobic | 47 | 37 | 98 |
| Gram-negative | Facultative Anaerobic | 57.2 | 35 | 37 |
| Gram-negative | Facultative Anaerobic | 45.3 | 35 | 40 |

**Sup. Table 2: Sources for all D10 values used to arrive at r**

| **Species** | **D10** |
|---|---|
| Staphylococcus aureus | 330 |
| Rubrobacter radiotolerans | 12000 |
| Rubrobacter xylanophilus | 6000 |
| Bacillus cereus | 400 |
| Bacillus pumilus | 3000 |
| Bacillus pumilus | 2260 |
| Bacillus pumilus | 1900 |
| Campylobacter coli | 220 |
| Campylobacter jejuni | 230 |
| Escherichia coli | 350 |
| Staphylococcus epidermidis | 310 |
| Shewanella oneidensis | 70 |
| Micrococcus luteus | 4400 |
| Rhodococcus erythropolis | 1350 |
| Salmonella enterica | 600 |
| Listeria monocytogenes | 310 |
| Listeria monocytogenes | 350 |
| Listeria monocytogenes | 450 |
| Listeria monocytogenes | 500 |
| Enterococcus faecalis | 1700 |
| Escherichia coli | 300 |
| Listeria monocytogenes | 620 |
| Staphylococcus aureus | 500 |
| Yersinia enterocolitica | 200 |
| Serratia marcescens | 60 |
| Shewanella putrefaciens | 110 |
| Enterococcus faecalis | 6000 |
| Lactobacillus acidophilus | 1400 |
| Lactobacillus acidophilus | 900 |
| Pediococcus pentosaceus | 800 |
| Pediococcus pentosaceus | 570 |
| Stenotrophomonas maltophilia | 530 |
| Streptococcus thermophilus | 590 |
| Streptococcus thermophilus | 470 |
| Bifidobacterium breve | 820 |
| Bifidobacterium breve | 260 |
| Lactococcus lactis | 585 |
| Bacillus cereus | 1000 |
| Bacillus sphaericus | 1500 |
| Micrococcus luteus | 1400 |
| Coxiella burnetii | 640 |
| Aeromonas hydrophila | 50 |
| Vibrio parahaemolyticus | 75 |
| Listeria monocytogenes | 180 |

| | |
|---|---|
| Neisseria gonorrhoeae | 125 |
| Enterococcus faecalis | 700 |
| Mycobacterium tuberculosis | 290 |
| Spirosoma radiotolerans | 9000 |
| Enterococcus faecalis | 3500 |
| Escherichia coli | 2000 |
| Mycobacterium smegmatis | 500 |
| Proteus vulgaris | 250 |
| Salmonella paratyphi | 120 |
| Salmonella typhimurium | 320 |
| Salmonella typhimurium | 570 |
| Salmonella typhimurium | 700 |
| Salmonella typhimurium | 200 |
| Salmonella typhimurium | 210 |
| Priestia megaterium | 400 |
| Deinococcus geothermalis | 15000 |
| Deinococcus radiodurans | 15000 |
| Bacillus cereus | 140 |
| Aquifex pyrophilus | 2842 |
| Hymenobacter marinus | 754 |
| Deinococcus geothermalis | 10000 |
| Deinococcus radiodurans | 16000 |
| Enterococcus faecium | 2000 |
| Escherichia coli | 700 |
| Pseudomonas putida | 250 |
| Shewanella oneidensis | 70 |
| Acinetobacter radioresistens | 1300 |
| Acinetobacter radioresistens | 2200 |
| Enterococcus faecalis | 1500 |
| Salmonella muenster | 600 |
| Campylobacter coli | 120 |
| Campylobacter coli | 140 |
| Campylobacter coli | 140 |
| Campylobacter jejuni | 190 |
| Campylobacter jejuni | 160 |
| Campylobacter jejuni | 130 |
| Campylobacter lari | 140 |
| Deinococcus radiodurans | 16000 |
| Escherichia coli | 700 |
| Thermus thermophilus | 800 |
| Pseudomonas putida | 300 |
| Vibrio cholera O1 | 120 |
| Listeria monocytogenes | 324 |
| Salmonella enterica | 250 |
| Salmonella enterica subsp. enterica serovar | 330 |
| Salmonella Senftenberg | 330 |

| | |
|---|---|
| Salmonella typhimurium | 280 |
| Vibrio parahaemolyticus | 30 |
| Vibrio parahaemolyticus | 40 |
| Klebsiella variicola | 184 |
| Morganella morganii | 295 |
| Proteus vulgaris | 309 |
| Enterobacter sp. BIGb0383 | 360 |
| Enterobacter sp. BIGb0383 | 480 |
| Staphylococcus aureus | 500 |
| Vibrio cholera O1 | 76 |
| Acinetobacter radioresistens | 1200 |
| Bacillus cereus | 420 |
| Aeromonas hydrophila | 150 |
| Bacillus cereus | 488 |
| Paenibacillus amylolyticus | 330 |
| Kocuria rhizophila | 2900 |
| Acinetobacter radioresistens | 5000 |
| Mycobacterium tuberculosis | 200 |
| Mycobacterium tuberculosis | 1800 |
| Aeromonas hydrophila | 250 |
| Aeromonas salmonicida | 230 |
| Shigella boydii | 300 |
| Shigella flexneri | 290 |
| Shigella sonnei | 750 |
| Deinococcus ficus | 7000 |
| Kineococcus radiotolerans | 2000 |
| Massilia aerilata | 3000 |
| Methylobacterium radiotoleran | 1000 |
| Yersinia enterocolitica | 110 |
| Listeria monocytogenes | 560 |

**mean D10 in Sup. Table 1**

| **Author** |
|---|
| A Adu-Gyamfi |
| A C Ferreira 1, M F Nobre, E Moore, F A Rainey, J R Battista, M S da Costa Nina Gunde-Cimerman 5, Jocelyne DiRu |
| A C Ferreira 1, M F Nobre, E Moore, F A Rainey, J R Battista, M S da Costa Nina Gunde-Cimerman 5, Jocelyne DiRu |
| A K Kotiranta 1, H Ito, M P Haapasalo, K Lounatmaa |
| A N Parisi, A D Antoine |
| A. Parisi and A. D. Antoine |
| Al Rawi A.M.; Hall A.N |
| Amol D.Raut a, RavindranathShashidhar b, JayantR.Bandekar b,n, BaluP.Kapadnis a |
| Amol D.Raut a, RavindranathShashidhar b, JayantR.Bandekar b,n, BaluP.Kapadnis a |
| Andrej Trampuz, Kerryl E. Piper, James M. Steckelberg, Robin Patel |
| Andrej Trampuz, Kerryl E. Piper, James M. Steckelberg, Robin Patel |
| Ashley R Brown 1, Elon Correa 2, Yun Xu 2, Najla AlMasoud 2, Simon M Pimblott 3, Royston Goodacre 4, Jonathan |
| Beth J. Pitonzo, Penny S. Amy and Mark Rudin |
| Beth J. Pitonzo, Penny S. Amy and Mark Rudin |
| Bougle D.; Stahl V.; Union des Cooperatives Laitieres, Isigny-sur-Mer |
| Brandao Areal, H.; Charbonneau, R.; Thibault, C. |
| Brandao Areal, H.; Charbonneau, R.; Thibault, C. |
| Brandao Areal, H.; Charbonneau, R.; Thibault, C. |
| Brandao Areal, H.; Charbonneau, R.; Thibault, C. |
| C Emborg |
| Christopher H Sommers 1, Kathleen T Rajkowski |
| Christopher H Sommers 1, Kathleen T Rajkowski |
| Christopher H Sommers 1, Kathleen T Rajkowski |
| Christopher H Sommers 1, Kathleen T Rajkowski |
| D. L. Dewey |
| DONALD W. THAYER, GLENN BOYD |
| E A CHRISTENSEN |
| E-Nam Hwang, Sang-Mo Kang, Jae-Kyung Kim1 |
| E-Nam Hwang, Sang-Mo Kang, Jae-Kyung Kim1 |
| E-Nam Hwang, Sang-Mo Kang, Jae-Kyung Kim1 |
| E-Nam Hwang, Sang-Mo Kang, Jae-Kyung Kim1 |
| E-Nam Hwang, Sang-Mo Kang, Jae-Kyung Kim1 |
| E-Nam Hwang, Sang-Mo Kang, Jae-Kyung Kim1 |
| E-Nam Hwang, Sang-Mo Kang, Jae-Kyung Kim1 |
| E-Nam Hwang, Sang-Mo Kang, Jae-Kyung Kim1 , Ju-Woon Lee1 , and Jong-Heum Park1* |
| E-Nam Hwang, Sang-Mo Kang, Jae-Kyung Kim1 , Ju-Woon Lee1 , and Jong-Heum Park1* |
| E-Nam Hwang, Sang-Mo Kang, Jae-Kyung Kim1 , Ju-Woon Lee1 , and Jong-Heum Park1* |
| Farrag, H. A., El-Bazza, Z. E., El-Fouly, M. E. Z. and El-Tablawy, S. Y. M |
| Farrag, H. A., El-Bazza, Z. E., El-Fouly, M. E. Z. and El-Tablawy, S. Y. M |
| Farrag, H. A., El-Bazza, Z. E., El-Fouly, M. E. Z. and El-Tablawy, S. Y. M |
| G H Scott 1, T F McCaul, J C Williams |
| H.O. Rashid, H. Ito* and I. Ishigaki |
| H.O. Rashid, H. Ito* and I. Ishigaki |
| Harsojo; Banati, D.; Ito, H. |

Ian J. Mehr, H. Steven Seifert
J Schubert, H Stegeman, M Swildens
Jackson V Watkins, Justin Bell, Phillip Knabenbauer, Alexander Brandl,  View ORCID ProfileKaren M Dobos
Jae-Jin Lee, Sathiyaraj Srinivasan, Sangyong Lim, Minho Joe, Seonghun Im, So Il Bae, Kyoung Ryun Park, Ji-Hee Har
John K. Dyer, A. W. Anderson, and Pisawat Dutiyabodhi
John K. Dyer, A. W. Anderson, and Pisawat Dutiyabodhi
John K. Dyer, A. W. Anderson, and Pisawat Dutiyabodhi
John K. Dyer, A. W. Anderson, and Pisawat Dutiyabodhi
John K. Dyer, A. W. Anderson, and Pisawat Dutiyabodhi
John K. Dyer, A. W. Anderson, and Pisawat Dutiyabodhi
Kazi Shamsuzzaman, Marlene Goodwin, Ian George, Harwant Singh
Kazi Shamsuzzaman, Marlene Goodwin, Ian George, Harwant Singh
Kazi Shamsuzzaman, Marlene Goodwin, Ian George, Harwant Singh
Kazi Shamsuzzaman, Marlene Goodwin, Ian George, Harwant Singh
Kiortsis, M., Vrantzas, N.
Kira S. Makarova, 1 , * Marina V. Omelchenko, 1 Elena K. Gaidamakova, 2 Vera Y. Matrosova, 2 Alexander Vasilenl
Kira S. Makarova, 1 , * Marina V. Omelchenko, 1 Elena K. Gaidamakova, 2 Vera Y. Matrosova, 2 Alexander Vasilenl
Kirschner, N Citri, A Levitzki, M Anbar
Kristina Beblo, Thierry Douki, Gottfried Schmalz, Reinhard Rachel, Reinhard Wirth, Harald Huber, Günther Reitz &
Kristina Beblo, Thierry Douki, Gottfried Schmalz, Reinhard Rachel, Reinhard Wirth, Harald Huber, Günther Reitz &
M J Daly 1, E K Gaidamakova, V Y Matrosova, A Vasilenko, M Zhai, A Venkateswaran, M Hess, M V Omelchenko, H
M J Daly 1, E K Gaidamakova, V Y Matrosova, A Vasilenko, M Zhai, A Venkateswaran, M Hess, M V Omelchenko, H
M J Daly 1, E K Gaidamakova, V Y Matrosova, A Vasilenko, M Zhai, A Venkateswaran, M Hess, M V Omelchenko, H
M J Daly 1, E K Gaidamakova, V Y Matrosova, A Vasilenko, M Zhai, A Venkateswaran, M Hess, M V Omelchenko, H
M J Daly 1, E K Gaidamakova, V Y Matrosova, A Vasilenko, M Zhai, A Venkateswaran, M Hess, M V Omelchenko, H
M J Daly 1, E K Gaidamakova, V Y Matrosova, A Vasilenko, M Zhai, A Venkateswaran, M Hess, M V Omelchenko, H
M M Garcia 1, B W Brooks, R B Stewart, W Dion, J R Trudel, T Ouwerkerk
M M Garcia 1, B W Brooks, R B Stewart, W Dion, J R Trudel, T Ouwerkerk
M M Garcia 1, B W Brooks, R B Stewart, W Dion, J R Trudel, T Ouwerkerk
M M Garcia 1, B W Brooks, R B Stewart, W Dion, J R Trudel, T Ouwerkerk
M.F. Patterson
M.F. Patterson
M.F. Patterson
M.F. Patterson
M.F. Patterson
M.F. Patterson
M.F. Patterson
Marina V Omelchenko, Yuri I Wolf, Elena K Gaidamakova, Vera Y Matrosova, Alexander Vasilenko, Min Zhai, Micha
Marina V Omelchenko, Yuri I Wolf, Elena K Gaidamakova, Vera Y Matrosova, Alexander Vasilenko, Min Zhai, Micha
Marina V Omelchenko, Yuri I Wolf, Elena K Gaidamakova, Vera Y Matrosova, Alexander Vasilenko, Min Zhai, Micha
Michael J Daly 1
Moraes, I. R. de, Gelli, D. S., Jakabi, M., & Mastro, N. L. D
Muhammad Tanveer Munir1,* and Michel Federighi2,*
N. P. TIWARI, R. B. MAXCY
N. P. TIWARI, R. B. MAXCY
N. P. TIWARI, R. B. MAXCY

N. P. TIWARI, R. B. MAXCY
P Dion 1, R Charbonneau, C Thibault
P Dion 1, R Charbonneau, C Thibault
P. Ramakrishna Reddy, Sanath H. Kumar, P. Layana, Binaya Bhusan Na
P. Ramakrishna Reddy, Sanath H. Kumar, P. Layana, Binaya Bhusan Na
P. Ramakrishna Reddy, Sanath H. Kumar, P. Layana, Binaya Bhusan Na
R L Ward, J G Yeager, and C S Ashley
R L Ward, J G Yeager, and C S Ashley
R. B. Webb
Rubio, 1997
Sahadev Chandra Saha and Balu Anand Chopade
Samia Ayari 1, Dominic Dussault, El Akrem Hayouni, Khanh Dang Vu, Moktar Hamdi, Monique Lacroix
Samuel A. Palumbo, Ronald K. Jenkins, Robert L. Buchanan, Donald W. Thayer
Shiv Shankar a, Peter Follett b, Samia Ayari a c, Farah Hossain a, Stephane Salmieri a, Monique Lacroix a
Shiv Shankar a, Peter Follett b, Samia Ayari a c, Farah Hossain a, Stephane Salmieri a, Monique Lacroix a
Sihem Guesmi a b, Petar Pujic c k, Imen Nouioui d e, Audrey Dubost c, Afef Najjari f, Kais Ghedira g, José M. Igual h
Taeko Yoshinaka, Keiji Yano & Hikoyuki Yamaguchi
Teri A. Grieb a, Ren-Yo Forng a, Richard E. Stafford a, Jack Lin a, Jamie Almeida a, Simon Bogdansky b, Chad Ronhc
Teri A. Grieb a, Ren-Yo Forng a, Richard E. Stafford a, Jack Lin a, Jamie Almeida a, Simon Bogdansky b, Chad Ronhc
Vandan Nagar,* Lipika Pansare Godambe & Ravindranath Shashidhar
Vandan Nagar,* Lipika Pansare Godambe & Ravindranath Shashidhar
Vandan Nagar,* Lipika Pansare Godambe & Ravindranath Shashidhar
Vandan Nagar,* Lipika Pansare Godambe & Ravindranath Shashidhar
Vandan Nagar,* Lipika Pansare Godambe & Ravindranath Shashidhar
Vera Y Matrosova # 1 2, Elena K Gaidamakova # 1 2, Kira S Makarova 3, Olga Grichenko 1 2, Polina Klimenkova 1 2
Xuesong Luo, Xian-Chun Zeng, Zhancan He, Xiaolu Lu, Jie Yuan, Jingjing Shi, Ming Liu, Yunfan Pan & Yan-Xin Wang
Xuesong Luo, Xian-Chun Zeng, Zhancan He, Xiaolu Lu, Jie Yuan, Jingjing Shi, Ming Liu, Yunfan Pan & Yan-Xin Wang
Xuesong Luo, Xian-Chun Zeng, Zhancan He, Xiaolu Lu, Jie Yuan, Jingjing Shi, Ming Liu, Yunfan Pan & Yan-Xin Wang
Yehia A. El-Zawahry† and D. B. Rowley

| PMID | DOI |
|---|---|
| 10.4314/jast.v14i1-2.44317 | |
| ggiero 6, Igor Shuryak 7, Andrew Oza | 10.1007/s007920050121 |
| ggiero 6, Igor Shuryak 7, Andrew Oza | 10.1007/s007920050121 |
| 10518726 | 10.1111/j.1574-6968.1999.tb08738.x |
| 4844266 | 10.1128/am.28.1.41-46.1974 |
| 4844266 | 10.1128/am.28.1.41-46.1974 |
| te Buffer on the Radiation Susceptibility of Bacillus Pumilus Strain E601. International Atomic Energy Agency (IAE | |
| | https://doi.org/10.1016/j.radphyschem.2011.09.003 |
| | https://doi.org/10.1016/j.radphyschem.2011.09.003 |
| 16914659 | 10.1099/jmm.0.46488-0. |
| 16914659 | 10.1099/jmm.0.46488-0. |
| 26098880 | 10.1371/journal.pone.0131249 |
| 10381842 | |
| 10381842 | |
| ISSN : 0074-1884 | |
| ISSN/ISBN: 0240-8813 | |
| ISSN/ISBN: 0240-8813 | |
| ISSN/ISBN: 0240-8813 | |
| ISSN/ISBN: 0240-8813 | |
| 4626006 | 10.1111/j.1699-0463.1972.tb00048.x |
| 21477481 | 10.4315/0362-028X.JFP-10-419 |
| 21477481 | 10.4315/0362-028X.JFP-10-419 |
| 21477481 | 10.4315/0362-028X.JFP-10-419 |
| 21477481 | 10.4315/0362-028X.JFP-10-419 |
| | https://doi.org/10.2307/3571410 |
| https://doi.org/10.1111/j.1745-4565.1996.tb00156.x | |
| 14164896 | 10.1111/apm.1964.61.3.483 |
| | https://doi.org/10.5851/kosfa.2013.33.3.335 |
| | https://doi.org/10.5851/kosfa.2013.33.3.335 |
| | https://doi.org/10.5851/kosfa.2013.33.3.335 |
| | https://doi.org/10.5851/kosfa.2013.33.3.335 |
| | https://doi.org/10.5851/kosfa.2013.33.3.335 |
| | https://doi.org/10.5851/kosfa.2013.33.3.335 |
| | https://doi.org/10.5851/kosfa.2013.33.3.335 |
| | DOI http://dx.do.org/10.5851/kosfa.2013.33.3.335 |
| | DOI http://dx.do.org/10.5851/kosfa.2013.33.3.335 |
| | https://doi.org/10.5851/kosfa.2013.33.3.335 |
| lora of Nigella sativa seeds and on its oil constituents. Acta Pharm. 50, 195-207. | |
| lora of Nigella sativa seeds and on its oil constituents. Acta Pharm. 50, 195-207. | |
| lora of Nigella sativa seeds and on its oil constituents. Acta Pharm. 50, 195-207. | |
| 2636260 | 10.1099/00221287-135-12-3263 |
| 24425563 | doi: 10.1007/BF01201947. |
| 24425563 | 10.1007/BF01201947 |
| ARN: JP1998005288 | |

| | | |
|---|---|---|
| 10094619 | | 10.1046/j.1365-2958.1998.01089.x |
| 6792101 | | 10.1080/09553008114551011 |
| | | 10.1101/2021.03.02.433681 |
| 24748440 | | 10.1128/am.14.1.92-97.1966 |
| 4958149 | | 10.1128/am.14.1.92-97.1966 |
| 4958149 | | 10.1128/am.14.1.92-97.1966 |
| 4958149 | | 10.1128/am.14.1.92-97.1966 |
| 4958149 | | 10.1128/am.14.1.92-97.1966 |
| 4958149 | | 10.1128/am.14.1.92-97.1966 |
| 4958149 | | 10.1128/am.14.1.92-97.1966 |
| | | https://doi.org/10.1016/1359-0197(89)90339-1 |
| | | https://doi.org/10.1016/1359-0197(89)90339-2 |
| | | https://doi.org/10.1016/1359-0197(89)90339-3 |
| | | https://doi.org/10.1016/1359-0197(89)90339-4 |
| | | https://doi.org/10.1007/BF01921611 |
| | 17895995 | 10.1371/journal.pone.0000955 |
| | 17895995 | 10.1371/journal.pone.0000955 |
| 4984594 | | 10.1080/09553007014550081 |
| 21638055 | | 10.1007/s00203-011-0718-5 |
| 21638055 | | 10.1007/s00203-011-0718-5 |
| | 15459345 | 10.1126/science.1103185 |
| | 15459345 | 10.1126/science.1103185 |
| | 15459345 | 10.1126/science.1103185 |
| | 15459345 | 10.1126/science.1103185 |
| | 15459345 | 10.1126/science.1103185 |
| | 15459345 | 10.1126/science.1103185 |
| 3651881 | | NA |
| 3651881 | | NA |
| 3651881 | | NA |
| 3651881 | | NA |
| 7786498 | | DOI: 10.1111/j.1472-765X.1995.tb01315.x |
| 7786499 | | DOI: 10.1111/j.1472-765X.1995.tb01315.x |
| 7786500 | | DOI: 10.1111/j.1472-765X.1995.tb01315.x |
| 7786498 | | DOI: 10.1111/j.1472-765X.1995.tb01315.x |
| 7786499 | | DOI: 10.1111/j.1472-765X.1995.tb01315.x |
| 7786500 | | DOI: 10.1111/j.1472-765X.1995.tb01315.x |
| 7786501 | | DOI: 10.1111/j.1472-765X.1995.tb01315.x |
| 16242020 | | 10.1186/1471-2148-5-57 |
| 16242020 | | 10.1186/1471-2148-5-57 |
| 16242020 | | 10.1186/1471-2148-5-57. |
| 22112864 | | 10.1016/j.dnarep.2011.10.024 |
| | | https://doi.org/10.1590/S1413-95961998000600001 |
| 32635407 | | 10.3390/foods9070878 |
| | | https://doi.org/10.1111/j.1365-2621.1972.tb02669.x |
| | | https://doi.org/10.1111/j.1365-2621.1972.tb02669.x |
| | | https://doi.org/10.1111/j.1365-2621.1972.tb02669.x |

| | |
|---|---|
| | https://doi.org/10.1111/j.1365-2621.1972.tb02669.x |
| 8069779 | 10.1139/m94-060 |
| 8069779 | 10.1139/m94-060 |
| | https://doi.org/10.4315/JFP-19-511 |
| | https://doi.org/10.4315/JFP-19-511 |
| | https://doi.org/10.4315/JFP-19-511 |
| 6789765 | 10.1128/aem.41.5.1123-1127.1981 |
| 6789765 | 10.1128/aem.41.5.1123-1127.1981 |
| | https://doi.org/10.2307/3571402 |
| 20120776.00 | DOI: 10.3329/bmrcb.v35.i2.2490 |
| 22986202 | 10.1016/j.fm.2012.07.016 |
| 30959721 | 10.4315/0362-028X-49.3.189 |
| 32147410 | 10.1016/j.micpath.2020.104118 |
| 32147410 | 10.1016/j.micpath.2020.104118 |
| ı, Guylaine Miotello i, Ameur Cherif j | https://doi.org/10.1016/j.ygeno.2020.11.029 |
| | https://doi.org/10.1080/00021369.1973.10861003 |
| 15576177 | 10.1016/j.biomaterials.2004.06.028 |
| 15576177 | 10.1016/j.biomaterials.2004.06.028 |
| | 10.1016/j.radphyschem.2011.04.002 |
| | 10.1016/j.radphyschem.2011.04.002 |
| | DOI: 10.1111/ijfs.13277 |
| | DOI: 10.1111/ijfs.13277 |
| | DOI: 10.1111/ijfs.13277 |
| 28775794 | 10.1186/s40793-017-0258-y |
| 25182517 | 10.1007/s10646-014-1325-4 |
| 25182517 | 10.1007/s10646-014-1325-4 |
| 25182517 | 10.1007/s10646-014-1325-4 |
| 243400 | 10.1128/aem.37.1.50-54.1979 |
| | https://www.anses.fr/en/system/files/BIORISK2016SA0081FiEN.pdf |

EA): IAEA.

**Sup. Table 3: TolRad Classifications on the Train/Test Sets**

**Train Set**

| **Species** | **Graph ID** | **Classification Success** |
|---|---|---|
| Acinetobacter calcaeceticus | 1 | Incorrect |
| Aeromonas hydrophila | 2 | Incorrect |
| Aeromonas salmonicida | 3 | Incorrect |
| Aquifex pyrophilus | 4 | Correct |
| Bacillus pumilus | 5 | Correct |
| Bacillus sphaericus | 6 | Correct |
| Campylobacter coli | 7 | Correct |
| Campylobacter jejuni | 8 | Correct |
| Campylobacter lari | 9 | Correct |
| Coxiella burnetii | 10 | Correct |
| Deinococcus geothermalis | 11 | Correct |
| Deinococcus radiodurans | 12 | Correct |
| Enterobacter sp. BIGb0383 | 13 | Correct |
| Enterococcus faecium | 14 | Correct |
| Enterococcus faecalis | 15 | Correct |
| Escherichia coli | 16 | Correct |
| Kineococcus radiotolerans | 17 | Correct |
| Klebsiella variicola | 18 | Correct |
| Kocuria rhizophila | 19 | Correct |
| Lactococcus lactis | 20 | Correct |
| Listeria monocytogenes | 21 | Correct |
| Methylobacterium radiotoleran | 22 | Correct |
| Micrococcus luteus | 23 | Correct |
| Morganella morganii | 24 | Correct |
| Mycobacterium smegmatis | 25 | Correct |
| Mycobacterium tuberculosis | 26 | Correct |
| Proteus vulgaris | 27 | Correct |
| Pseudomonas putida | 28 | Incorrect |
| Rhodococcus erythropolis | 29 | Correct |
| Rubrobacter radiotolerans | 30 | Correct |
| Rubrobacter xylanophilus | 31 | Correct |
| Salmonella Senftenberg | 32 | Correct |
| Salmonella enterica subsp. enteric | 33 | Correct |
| Salmonella paratyphi | 34 | Incorrect |
| Serratia marcescens | 35 | Correct |
| Shewanella putrefaciens | 36 | Correct |
| Staphylococcus aureus | 37 | Correct |
| Staphylococcus epidermidis | 38 | Correct |
| Stenotrophomonas maltophilia | 39 | Correct |
| Vibrio parahaemolyticus | 40 | Correct |

**Test Set**

| Species | Graph ID | Classification Success |
|---|---|---|
| Acinetobacter radioresistens | 1 | Correct |
| Bacillus cereus | 2 | Correct |
| Bifidobacterium breve | 3 | Correct |
| Deinococcus ficus | 4 | Correct |
| Lactobacillus acidophilus | 5 | Correct |
| Neisseria gonorrhoeae | 6 | Correct |
| Paenibacillus amylolyticus | 7 | Correct |
| Pediococcus pentosaceus | 8 | Correct |
| Priestia megaterium | 9 | Correct |
| Salmonella enterica | 10 | Correct |
| Salmonella muenster | 11 | Correct |
| Salmonella typhimurium | 12 | Correct |
| Shewanella oneidensis | 13 | Correct |
| Shigella boydii | 14 | Incorrect |
| Shigella flexneri | 15 | Correct |
| Shigella sonnei | 16 | Correct |
| Spirosoma radiotolerans | 17 | Correct |
| Streptococcus thermophilus | 18 | Correct |
| Thermus thermophilus | 19 | Correct |
| Vibrio cholera O1 | 20 | Incorrect |
| Yersinia enterocolitica | 21 | Correct |

**EggNogMapper**

| Species | Graph ID | Classification Success |
|---|---|---|
| Acinetobacter calcaeceticus | 1 | Correct |
| Vibrio cholera O1 | 2 | Correct |
| Neisseria gonorrhoeae | 3 | Correct |
| Lactococcus lactis | 4 | Correct |
| Shigella flexneri | 5 | Correct |
| Salmonella typhimurium | 6 | Correct |
| Pseudomonas putida | 7 | Correct |
| Coxiella burnetii | 8 | Correct |
| Bacillus cereus | 9 | Correct |
| Escherichia coli | 10 | Correct |
| Campylobacter jejuni | 11 | Correct |
| Streptococcus thermophilus | 12 | Correct |
| Staphylococcus epidermidis | 13 | Correct |
| Lactobacillus acidophilus | 14 | Correct |
| Staphylococcus aureus | 15 | Correct |
| Rubrobacter xylanophilus | 16 | Correct |
| Pediococcus pentosaceus | 17 | Correct |
| Aeromonas hydrophila | 18 | Correct |

| | | |
|---|---|---|
| Mycobacterium smegmatis | 19 | Correct |
| Kineococcus radiotolerans | 20 | Correct |
| Salmonella paratyphi | 21 | Incorrect |
| Methylobacterium radiotoleran | 22 | Correct |
| Shigella boydii | 23 | Incorrect |
| Priestia megaterium | 24 | Correct |
| Stenotrophomonas maltophilia | 25 | Correct |
| Thermus thermophilus | 26 | Correct |
| Shigella sonnei | 27 | Correct |
| Shewanella oneidensis | 28 | Correct |
| Shewanella putrefaciens | 29 | Correct |
| Enterococcus faecium | 30 | Correct |
| Mycobacterium tuberculosis | 31 | Correct |
| Listeria monocytogenes | 32 | Correct |
| Vibrio parahaemolyticus | 33 | Correct |
| Deinococcus geothermalis | 34 | Correct |
| Morganella morganii | 35 | Correct |
| Acinetobacter radioresistens | 36 | Correct |
| Rubrobacter radiotolerans | 37 | Correct |
| Campylobacter lari | 38 | Correct |
| Spirosoma radiotolerans | 39 | Correct |
| Serratia marcescens | 40 | Correct |
| Kocuria rhizophila | 41 | Correct |
| Bifidobacterium breve | 42 | Correct |
| Bacillus sphaericus | 43 | Correct |
| Deinococcus ficus | 44 | Correct |
| Paenibacillus amylolyticus | 45 | Correct |
| Enterobacter sp. BIGb0383 | 46 | Correct |
| Aeromonas salmonicida | 47 | Correct |
| Rhodococcus erythropolis | 48 | Correct |
| Salmonella Senftenberg | 49 | Correct |
| Campylobacter coli | 50 | Correct |
| Bacillus pumilus | 51 | Correct |
| Salmonella enterica subsp. enteric | 52 | Correct |
| Salmonella muenster | 53 | Correct |
| Salmonella enterica | 54 | Correct |
| Enterococcus faecalis | 55 | Correct |
| Deinococcus radiodurans | 56 | Correct |
| Klebsiella variicola | 57 | Correct |
| Aquifex pyrophilus | 58 | Correct |
| Micrococcus luteus | 59 | Correct |
| Proteus vulgaris | 60 | Correct |
| Yersinia enterocolitica | 61 | Correct |

**Sup. Table 4: All TolRad classifications**

| ATCC_Name | ATCC_Strain | UniProt_name | Phylum | ID |
|---|---|---|---|---|
| **NIH ATCC Dataset** | | | | |
| Alistipes putredinis¬ | VPI 3293 [1Ando, CCU | Alistipes putredini | Bacteroidetes | ATCC_2 |
| Bacteroides caccae¬† | VPI 3452A [CIP 10420 | Bacteroides cacca | Bacteroidetes | ATCC_7 |
| Bacteroides ovatus¬† | [NCTC 11153] | Bacteroides ovatu | Bacteroidetes | ATCC_8 |
| Bacteroides stercoris | VPI B5-21 [CIP 10420 | Bacteroides sterco | Bacteroidetes | ATCC_9 |
| Bacteroides uniformi | Bacteroides thetaiota | Bacteroides unifor | Bacteroidetes | ATCC_10 |
| Capnocytophaga sputigena¬† | Leadbetter et | Capnocytophaga s | Bacteroidetes | ATCC_12 |
| Eikenella corrodens¬ | 333/54-55 | Eikenella corroder | Pseudomonadota | ATCC_21 |
| Eubacterium siraeum | VPI T9-50-2 [DSM 157 | Eubacterium sirae | Bacillota | ATCC_23 |
| Helicobacter canade | NLEP 16143 [98-5491, | Helicobacter cana | Pseudomonadota | ATCC_28 |
| Parabacteroides mer | VPI T4-1 [CIP 104202T | Parabacteroides m | Bacteroidetes | ATCC_32 |
| Abiotrophia defectiva | SC10 [CIP 103242] | Abiotrophia defec | Bacillota | ATCC_1 |
| Anaerococcus hydrog | JCM 7635 [DSM 7454, | Anaerococcus hyd | Bacillota | ATCC_3 |
| Anaerococcus lactoly | JCM 8140 [CIP 10372 | Anaerococcus lact | Bacillota | ATCC_4 |
| Anaerococcus tetradi | GIFU 7672 [CCM 3634 | Anaerococcus tetr | Bacillota | ATCC_5 |
| Anaerofustis stercori | WAL 14563 [CCUG 47 | Anaerofustis sterc | Bacillota | ATCC_6 |
| Bifidobacterium den | B757 | Bifidobacterium d | Actinobacteria | ATCC_11 |
| Clostridium bolteae | WAL 16351 [CCUG 46 | Enterocloster bolt | Bacillota | ATCC_13 |
| Clostridium leptum¬ | VPI T7-24-2 [DSM 753 | Clostridium leptur | Bacillota | ATCC_14 |
| Clostridium scindens | VPI 13733 | Clostridium scinde | Bacillota | ATCC_15 |
| Clostridium spiroforn | [NCTC 11211, VPI C28 | [Clostridium] spirc | Bacillota | ATCC_16 |
| Clostridium sporogenes¬† | (Metchnikoff) Be | Clostridium sporo | Bacillota | ATCC_17 |
| Collinsella aerofacier | VPI 1003 [DSM 3979, | Collinsella aerofac | Actinobacteria | ATCC_18 |
| Coprococcus eutactu | VPI C33-22 | Coprococcus euta | Bacillota | ATCC_19 |
| Dorea formicigenera | VPI C8-13 [JCM 9500] | Dorea formicigene | Bacillota | ATCC_20 |
| Eubacterium dolichu | VPI C9-20 [DSM 3991] | Eubacterium dolic | Bacillota | ATCC_22 |
| Eubacterium ventrios | VPI 1013B | Eubacterium vent | Bacillota | ATCC_24 |
| Fusobacterium ulcer | NCTC 12111 | Fusobacterium ulc | Fusobacteria | ATCC_25 |
| Gardnerella vaginalis | 317 | Gardnerella vagin | Actinobacteria | ATCC_26 |
| Helicobacter bilis¬†F | SLH-38264 | Helicobacter bilis | Pseudomonadota | ATCC_27 |
| Helicobacter wingha | NLEP 97-1090 | Helicobacter wing | Pseudomonadota | ATCC_29 |
| Lactobacillus buchneri¬† | (Henneberg) Berge | Lentilactobacillus | Bacillota | ATCC_30 |
| Lactobacillus salivari | HO66 | Ligilactobacillus sa | Bacillota | ATCC_31 |
| Parvimonas micra¬† | VPI 5464 [3119B, DSM | Parvimonas micra | Bacillota | ATCC_33 |
| Providencia stuartii¬ | NCDC 3646-51 | Providencia stuart | Pseudomonadota | ATCC_34 |
| Pseudoflavonifractor | VPI R2-29-1 | Pseudoflavonifrac | Bacillota | ATCC_35 |
| Ruminococcus gnavu | VPI C7-9 | [Ruminococcus] g | Bacillota | ATCC_36 |
| Ruminococcus torqu | VPI B2-51 | [Ruminococcus] tc | Bacillota | ATCC_37 |
| **UniProt Human Oral Dataset** | | | | |
| Campylobacter gracilis RM3268 | | | Proteobacteria | Oral_7 |
| Capnocytophaga granulosa | | | Bacteroidetes | Oral_9 |

| | | |
|---|---|---|
| Capnocytophaga granulosa (strain ATCC 51502 / DSM 11449 / | Bacteroidetes | Oral_10 |
| Cryptobacterium curtum (strain ATCC 700683 / DSM 15641 / ( | Actinobacteria | Oral_15 |
| Eikenella corrodens CC92I | Proteobacteria | Oral_16 |
| Prevotella denticola | Bacteroidetes | Oral_32 |
| Prevotella nigrescens ATCC 33563 | Bacteroidetes | Oral_34 |
| Tannerella forsythia (strain ATCC 43037 / JCM 10827 / CCUG 2 | Bacteroidetes | Oral_42 |
| Veillonella parvula ATCC 17745 | Firmicutes | Oral_46 |
| Filifactor alocis (strain ATCC 35896 / D40 B5) (Fusobacterium a | Firmicutes | Oral_1 |
| Abiotrophia defectiva ATCC 49176 | Firmicutes | Oral_2 |
| Actinomyces graevenitzii C83 | Actinobacteria | Oral_3 |
| Actinomyces massiliensis F0489 | Actinobacteria | Oral_4 |
| Anaeroglobus geminatus F0357 | Firmicutes | Oral_5 |
| Bulleidia extructa W1219 | Firmicutes | Oral_6 |
| Capnocytophaga gingivalis | Bacteroidetes | Oral_8 |
| Cardiobacterium hominis (strain ATCC 15826 / DSM 8339 / NC | Proteobacteria | Oral_11 |
| Cardiobacterium valvarum | Proteobacteria | Oral_12 |
| Corynebacterium durum F0235 | Actinobacteria | Oral_13 |
| Corynebacterium matruchotii ATCC 14266 | Actinobacteria | Oral_14 |
| Elizabethkingia meningoseptica (Chryseobacterium meningose | Bacteroidetes | Oral_17 |
| Eubacterium brachy ATCC 33089 | Firmicutes | Oral_18 |
| Haemophilus parahaemolyticus | Proteobacteria | Oral_19 |
| Kingella oralis ATCC 51147 | Proteobacteria | Oral_20 |
| Lancefieldella parvula (strain ATCC 33793 / DSM 20469 / CCU( | Actinobacteria | Oral_21 |
| Lautropia mirabilis ATCC 51599 | Proteobacteria | Oral_22 |
| Leptotrichia buccalis (strain ATCC 14201 / DSM 1135 / JCM 12 | Fusobacteria | Oral_23 |
| Leptotrichia hofstadii F0254 | Fusobacteria | Oral_24 |
| Megasphaera micronuciformis F0359 | Firmicutes | Oral_25 |
| Neisseria bacilliformis ATCC BAA-1200 | Proteobacteria | Oral_26 |
| Neisseria elongata subsp. glycolytica ATCC 29315 | Proteobacteria | Oral_27 |
| Oribacterium sinus F0268 | Firmicutes | Oral_28 |
| Porphyromonas gingivalis (strain ATCC BAA-308 / W83) | Bacteroidetes | Oral_29 |
| Porphyromonas catoniae ATCC 51270 | Bacteroidetes | Oral_30 |
| Porphyromonas endodontalis (strain ATCC 35406 / BCRC 1449 | Bacteroidetes | Oral_31 |
| Prevotella loescheii DSM 19665 = JCM 12249 = ATCC 15930 | Bacteroidetes | Oral_33 |
| Rothia aeria | Actinobacteria | Oral_35 |
| Rothia dentocariosa | Actinobacteria | Oral_36 |
| Selenomonas infelix ATCC 43532 | Firmicutes | Oral_37 |
| Selenomonas sputigena (strain ATCC 35185 / DSM 20758 / VP | Firmicutes | Oral_38 |
| Streptococcus intermedius B196 | Firmicutes | Oral_39 |
| Streptococcus mutans serotype c (strain ATCC 700610 / UA15 | Firmicutes | Oral_40 |
| Streptococcus sanguinis (strain SK36) | Firmicutes | Oral_41 |
| Treponema maltophilum ATCC 51939 | Spirochaetes | Oral_43 |
| Treponema socranskii subsp. socranskii VPI DR56BR1116 = AT | Spirochaetes | Oral_44 |

| | | | |
|---|---|---|---|
| Fusobacterium nucleatum subsp. vincentii ATCC 49256 | | Fusobacteria | Oral_45 |
| **UniProt Human Skin Dataset** | | | |
| Neisseria cinerea | Neisseria cinerea | Proteobacteria | Skin_14 |
| Neisseria flavescens NRL30031/H210 | Neisseria Flavesce | Proteobacteria | Skin_15 |
| Pseudomonas aeruginosa (strain ATCC 156! | pseudomonas fluc | Proteobacteria | Skin_17 |
| Veillonella parvula (strain ATCC 10790 / DSI | Veillonella parvula | Firmicutes | Skin_27 |
| Acinetobacter lwoffii NIPH 715 | Acinetobacter Lwc | Proteobacteria | Skin_1 |
| Corynebacterium amycolatum | Corynebacterium a | Actinobacteria | Skin_2 |
| Corynebacterium kroppenstedtii (strain DSI | Corynebacterium k | Actinobacteria | Skin_3 |
| Corynebacterium minutissimum | Corynebacterium r | Actinobacteria | Skin_4 |
| Corynebacterium resistens (strain DSM 451 | Corynebacterium r | Actinobacteria | Skin_5 |
| Corynebacterium simulans | Corynebacterium s | Actinobacteria | Skin_6 |
| Corynebacterium tuberculostearicum SK14 | Corynebacterium t | Actinobacteria | Skin_7 |
| Cutibacterium acnes (strain DSM 16379 / K | Propionibacterium | Actinobacteria | Skin_8 |
| Enhydrobacter aerosaccus | Enhydrobacter aer | Proteobacteria | Skin_9 |
| Gardnerella vaginalis 6420B | Gardmerella vagin | Actinobacteria | Skin_10 |
| Gordonia bronchialis (strain ATCC 25592 / I | Gordonia bronchia | Actinobacteria | Skin_11 |
| Lactobacillus crispatus | lactobacillus crispa | Firmicutes | Skin_12 |
| Micrococcus luteus (strain ATCC 4698 / DSM | Micrococcus luteu | Actinobacteria | Skin_13 |
| Porphyromonas asaccharolytica (strain ATC | porphyromonas as | Bacteroidetes | Skin_16 |
| Staphylococcus capitis | Staphylococcus ca | Firmicutes | Skin_18 |
| Staphylococcus hominis | Staphylococcus ho | Firmicutes | Skin_19 |
| Staphylococcus warneri | Staphylococcus wa | Firmicutes | Skin_20 |
| Streptococcus mitis SK1073 | Streptococcus mit | Firmicutes | Skin_21 |
| Streptococcus oralis subsp. oralis | Streptococcus ora | Firmicutes | Skin_22 |
| Streptococcus pseudopneumoniae | Streptococcus pse | Firmicutes | Skin_23 |
| Streptococcus pyogenes serotype M1 | Streptococcus pyo | Firmicutes | Skin_24 |
| Streptococcus salivarius | Streptococcus sali | Firmicutes | Skin_25 |
| Streptococcus sanguinis (strain SK36) | Streptococcus san | Firmicutes | Skin_26 |
| Weeksella virosa (strain ATCC 43766 / DSM | weeksella virosa | Bacteroidetes | Skin_28 |
| **UniProt Human Gut Dataset** | | | |
| Alcaligenes xylosoxydans xylosoxydans (Achromobacter xylosc | | Proteobacteria | Gut_2 |
| Alistipes shahii WAL 8301 | | Bacteroidetes | Gut_3 |
| Bacteroides fragilis (strain ATCC 25285 / DSM 2151 / CCUG 48 | | Bacteroidetes | Gut_5 |
| Bacteroides stercoris | | Bacteroidetes | Gut_6 |
| Bacteroides thetaiotaomicron (strain ATCC 29148 / DSM 2079 | | Bacteroidetes | Gut_7 |
| Gordonibacter pamelaeae 7-10-1-b | | Actinobacteria | Gut_25 |
| Helicobacter pylori (strain ATCC 700392 / 26695) (Campylobac | | Proteobacteria | Gut_26 |
| Klebsiella pneumoniae subsp. pneumoniae (strain HS11286) | | Proteobacteria | Gut_29 |
| Odoribacter splanchnicus (strain ATCC 29572 / DSM 20712 / C | | Bacteroidetes | Gut_34 |
| Parabacteroides merdae ATCC 43184 | | Bacteroidetes | Gut_36 |
| Prevotella copri DSM 18205 | | Bacteroidetes | Gut_37 |
| Acinetobacter baumannii | | Proteobacteria | Gut_1 |

| | | |
|---|---|---|
| Bacteroides eggerthii | Bacteroidetes | Gut_4 |
| Bifidobacterium adolescentis (strain ATCC 15703 / DSM 2008 | Actinobacteria | Gut_8 |
| Bifidobacterium bifidum (strain PRL2010) | Actinobacteria | Gut_9 |
| Bifidobacterium longum (strain NCC 2705) | Actinobacteria | Gut_10 |
| Bifidobacterium pseudocatenulatum | Actinobacteria | Gut_11 |
| Cloacibacillus evryensis DSM 19522 | Synergistetes | Gut_12 |
| Clostridium beijerinckii (Clostridium MP) | Firmicutes | Gut_13 |
| Collinsella aerofaciens | Actinobacteria | Gut_14 |
| Coprococcus catus | Firmicutes | Gut_15 |
| Coprococcus comes ATCC 27758 | Firmicutes | Gut_16 |
| Dorea formicigenerans CAG:28 | Firmicutes | Gut_17 |
| Eggerthella lenta (strain ATCC 25559 / DSM 2243 / CCUG 173 | Actinobacteria | Gut_18 |
| Erysipelatoclostridium ramosum DSM 1402 | Firmicutes | Gut_19 |
| Faecalibacterium prausnitzii M21/2 | Firmicutes | Gut_20 |
| Fusobacterium gonidiaformans 3-1-5R | Fusobacteria | Gut_21 |
| Fusobacterium mortiferum ATCC 9817 | Fusobacteria | Gut_22 |
| Fusobacterium nucleatum subsp. animalis ATCC 51191 | Fusobacteria | Gut_23 |
| Fusobacterium varium ATCC 27725 | Fusobacteria | Gut_24 |
| Holdemanella biformis DSM 3989 | Firmicutes | Gut_27 |
| Holdemania filiformis DSM 12042 | Firmicutes | Gut_28 |
| Lactobacillus amylovorus DSM 16698 | Firmicutes | Gut_30 |
| Lactobacillus gasseri (strain ATCC 33323 / DSM 20243 / BCRC | Firmicutes | Gut_31 |
| Megamonas funiformis YIT 11815 | Firmicutes | Gut_32 |
| Mitsuokella jalaludinii | Firmicutes | Gut_33 |
| Parabacteroides distasonis (strain ATCC 8503 / DSM 20701 / | Bacteroidetes | Gut_35 |
| Proteus mirabilis | Proteobacteria | Gut_38 |
| Roseburia intestinalis CAG:13 | Firmicutes | Gut_39 |
| [Ruminococcus] gnavus CC55_001C | Firmicutes | Gut_40 |
| Shigella sonnei (strain Ss046) | Proteobacteria | Gut_41 |

| **Canadian High Arctic MAG Dataset** | **Phylum** | **ID** |
|---|---|---|
| | Proteobacteria; | Glacial4_7 |
| | Proteobacteria; | Glacial4_25 |
| | Proteobacteria; | Glacial4_1 |
| | Proteobacteria; | Glacial4_2 |
| | Desulfobacterota; | Glacial4_3 |
| | Bacteroidetes; | Glacial4_4 |
| | Proteobacteria; | Glacial4_5 |
| | Proteobacteria; | Glacial4_6 |
| | Chloroflexi; | Glacial4_8 |
| | Bacteroidetes; | Glacial4_9 |
| | Proteobacteria; | Glacial4_10 |

| | |
|---|---|
| Proteobacteria; | Glacial4_11 |
| Proteobacteria; | Glacial4_12 |
| Desulfuromonadota; | Glacial4_13 |
| Elusimicrobia; | Glacial4_14 |
| Proteobacteria; | Glacial4_15 |
| Proteobacteria; | Glacial4_16 |
| Proteobacteria; | Glacial4_17 |
| Planctomycetes; | Glacial4_18 |
| Proteobacteria; | Glacial4_19 |
| Deinococci; | Glacial4_20 |
| Proteobacteria; | Glacial4_21 |
| Proteobacteria; | Glacial4_22 |
| Firmicutes; | Glacial4_23 |
| Bacteroidetes; | Glacial4_24 |
| Proteobacteria; | Glacial4_26 |
| Proteobacteria; | Glacial4_27 |
| Desulfobacterota; | Glacial4_28 |
| Actinobacteria; | Glacial4_29 |
| Proteobacteria; | Glacial4_30 |

**Human Microbiome MAG Dataset**

| | |
|---|---|
| Firmicutes | HumanMB_3 |
| Firmicutes | HumanMB_3 |
| Firmicutes | HumanMB_5 |
| Bacteroidota | HumanMB_5 |
| Firmicutes | HumanMB_5 |
| Bacteroidota | HumanMB_7 |
| Firmicutes | HumanMB_7 |
| Firmicutes | HumanMB_1 |
| Proteobacteria | HumanMB_2 |
| Firmicutes | HumanMB_3 |
| Firmicutes | HumanMB_4 |
| Proteobacteria | HumanMB_5 |
| Firmicutes | HumanMB_6 |
| Proteobacteria | HumanMB_7 |
| Proteobacteria | HumanMB_8 |
| Firmicutes | HumanMB_9 |
| Proteobacteria | HumanMB_1 |
| Proteobacteria | HumanMB_1 |
| Proteobacteria | HumanMB_1 |
| Firmicutes | HumanMB_1 |
| Firmicutes | HumanMB_1 |
| Proteobacteria | HumanMB_1 |
| Proteobacteria | HumanMB_1 |

| | |
|---|---|
| Firmicutes | HumanMB_1 |
| Proteobacteria | HumanMB_1 |
| Proteobacteria | HumanMB_1 |
| Actinobacteriota | HumanMB_2 |
| Firmicutes | HumanMB_2 |
| Proteobacteria | HumanMB_2 |
| Proteobacteria | HumanMB_2 |
| Proteobacteria | HumanMB_2 |
| Firmicutes | HumanMB_2 |
| Firmicutes | HumanMB_2 |
| Proteobacteria | HumanMB_2 |
| Proteobacteria | HumanMB_2 |
| Firmicutes | HumanMB_2 |
| Proteobacteria | HumanMB_3 |
| Firmicutes | HumanMB_3 |
| Firmicutes | HumanMB_3 |
| Firmicutes | HumanMB_3 |
| Firmicutes | HumanMB_3 |
| Firmicutes | HumanMB_3 |
| Firmicutes | HumanMB_3 |
| Firmicutes | HumanMB_3 |
| Firmicutes | HumanMB_4 |
| Proteobacteria | HumanMB_4 |
| Firmicutes | HumanMB_4 |
| Firmicutes | HumanMB_4 |
| Firmicutes | HumanMB_4 |
| Firmicutes | HumanMB_4 |
| Firmicutes | HumanMB_4 |
| Firmicutes | HumanMB_4 |
| Firmicutes | HumanMB_4 |
| Firmicutes | HumanMB_4 |
| Firmicutes | HumanMB_5 |
| Firmicutes | HumanMB_5 |
| Firmicutes | HumanMB_5 |
| Firmicutes | HumanMB_5 |
| Firmicutes | HumanMB_5 |
| Firmicutes | HumanMB_5 |
| Firmicutes | HumanMB_5 |
| Firmicutes | HumanMB_6 |
| Firmicutes | HumanMB_6 |
| Firmicutes | HumanMB_6 |
| Firmicutes | HumanMB_6 |
| Firmicutes | HumanMB_6 |

| | |
|---|---|
| Firmicutes | HumanMB_6 |
| Firmicutes | HumanMB_6 |
| Firmicutes | HumanMB_6 |
| Bacteroidota | HumanMB_6 |
| Firmicutes | HumanMB_6 |
| Firmicutes | HumanMB_7 |
| Firmicutes | HumanMB_7 |
| Firmicutes | HumanMB_7 |
| Firmicutes | HumanMB_7 |
| Firmicutes | HumanMB_7 |
| Firmicutes | HumanMB_7 |
| Firmicutes | HumanMB_7 |
| Firmicutes | HumanMB_7 |
| Firmicutes | HumanMB_8 |
| Firmicutes | HumanMB_8 |
| Firmicutes | HumanMB_8 |
| Firmicutes | HumanMB_8 |
| Bacteroidota | HumanMB_8 |
| Proteobacteria | HumanMB_8 |
| Firmicutes | HumanMB_8 |
| Firmicutes | HumanMB_8 |
| Firmicutes | HumanMB_8 |
| Firmicutes | HumanMB_8 |
| Firmicutes | HumanMB_9 |
| Firmicutes | HumanMB_9 |
| Firmicutes | HumanMB_9 |
| Firmicutes | HumanMB_9 |
| Proteobacteria | HumanMB_9 |
| Proteobacteria | HumanMB_9 |
| Proteobacteria | HumanMB_9 |

**Deep Ocean MAG Dataset**

| | |
|---|---|
| Planctomycetota | DeepOcean4_ |
| Acidobacteriota | DeepOcean4_ |
| Proteobacteria | DeepOcean4_ |
| Proteobacteria | DeepOcean4_ |
| Proteobacteria | DeepOcean4_ |
| Bacteroidota | DeepOcean4_ |
| Proteobacteria | DeepOcean4_ |
| Bdellovibrionota | DeepOcean4_ |
| Planctomycetota | DeepOcean4_ |
| Bacteroidota | DeepOcean4_ |
| Marinisomatota | DeepOcean4_ |
| Bacteroidota | DeepOcean4_ |

| | |
|---|---|
| Chloroflexota | DeepOcean4_ |
| Marinisomatota | DeepOcean4_ |
| Bacteroidota | DeepOcean4_ |
| Crenarchaeota | DeepOcean4_ |
| Marinisomatota | DeepOcean4_ |
| Proteobacteria | DeepOcean4_ |
| Planctomycetota | DeepOcean4_ |
| Chloroflexota | DeepOcean4_ |
| UBA8248 | DeepOcean4_ |
| Thermoplasmatota | DeepOcean4_ |
| Planctomycetota | DeepOcean4_ |
| Proteobacteria | DeepOcean4_ |
| Bdellovibrionota | DeepOcean4_ |
| Nitrospinota | DeepOcean4_ |
| Proteobacteria | DeepOcean4_ |
| Crenarchaeota | DeepOcean4_ |
| Marinisomatota | DeepOcean4_ |
| Bacteroidota | DeepOcean4_ |
| Proteobacteria | DeepOcean4_ |
| Crenarchaeota | DeepOcean4_ |
| Marinisomatota | DeepOcean4_ |
| Marinisomatota | DeepOcean4_ |
| Proteobacteria | DeepOcean4_ |
| Bacteroidota | DeepOcean4_ |
| Latescibacterota | DeepOcean4_ |
| Verrucomicrobiota | DeepOcean4_ |
| Marinisomatota | DeepOcean4_ |
| Acidobacteriota | DeepOcean4_ |
| Planctomycetota | DeepOcean4_ |
| Marinisomatota | DeepOcean4_ |
| Nanoarchaeota | DeepOcean4_ |
| Patescibacteria | DeepOcean4_ |
| Planctomycetota | DeepOcean4_ |
| Thermoplasmatota | DeepOcean4_ |
| Chloroflexota | DeepOcean4_ |
| Latescibacterota | DeepOcean4_ |
| Thermoplasmatota | DeepOcean4_ |
| Thermoplasmatota | DeepOcean4_ |
| Proteobacteria | DeepOcean4_ |
| Thermoplasmatota | DeepOcean4_ |
| Acidobacteriota | DeepOcean4_ |
| Proteobacteria | DeepOcean4_ |
| Proteobacteria | DeepOcean4_ |

| | |
|---|---|
| Planctomycetota | DeepOcean4_ |
| Marinisomatota | DeepOcean4_ |
| Acidobacteriota | DeepOcean4_ |
| Myxococcota | DeepOcean4_ |
| Planctomycetota | DeepOcean4_ |
| Marinisomatota | DeepOcean4_ |
| Proteobacteria | DeepOcean4_ |
| Proteobacteria | DeepOcean4_ |
| Marinisomatota | DeepOcean4_ |
| Planctomycetota | DeepOcean4_ |
| Acidobacteriota | DeepOcean4_ |
| Bacteroidota | DeepOcean4_ |
| Proteobacteria | DeepOcean4_ |
| Verrucomicrobiota | DeepOcean4_ |
| Gemmatimonadota | DeepOcean4_ |
| Proteobacteria | DeepOcean4_ |
| Verrucomicrobiota | DeepOcean4_ |
| Bacteroidota | DeepOcean4_ |
| Myxococcota | DeepOcean4_ |
| Acidobacteriota | DeepOcean4_ |
| Cyanobacteria | DeepOcean4_ |
| Bacteroidota | DeepOcean4_ |
| Marinisomatota | DeepOcean4_ |
| Crenarchaeota | DeepOcean4_ |
| Marinisomatota | DeepOcean4_ |
| Planctomycetota | DeepOcean4_ |
| Eremiobacterota | DeepOcean4_ |
| Nanoarchaeota | DeepOcean4_ |
| Binatota | DeepOcean4_ |
| Poribacteria | DeepOcean4_ |
| Bacteroidota | DeepOcean4_ |
| Bacteroidota | DeepOcean4_ |
| Acidobacteriota | DeepOcean4_ |
| Planctomycetota | DeepOcean4_ |
| Planctomycetota | DeepOcean4_ |
| Proteobacteria | DeepOcean4_ |
| Verrucomicrobiota | DeepOcean4_ |
| Acidobacteriota | DeepOcean4_ |
| Thermoplasmatota | DeepOcean4_ |
| Nanoarchaeota | DeepOcean4_ |
| Myxococcota | DeepOcean4_ |
| Proteobacteria | DeepOcean4_ |
| Thermoplasmatota | DeepOcean4_ |

| | |
|---|---|
| Patescibacteria | DeepOcean4_ |
| Bdellovibrionota | DeepOcean4_ |
| Patescibacteria | DeepOcean4_ |
| Bacteroidota | DeepOcean4_ |
| Bacteroidota | DeepOcean4_ |
| Bacteroidota | DeepOcean4_ |
| Bacteroidota | DeepOcean4_ |
| Gemmatimonadota | DeepOcean4_ |
| Gemmatimonadota | DeepOcean4_ |
| Bacteroidota | DeepOcean4_ |
| Bacteroidota | DeepOcean4_ |
| Bacteroidota | DeepOcean4_ |
| Bacteroidota | DeepOcean4_ |
| Marinisomatota | DeepOcean4_ |
| Verrucomicrobiota | DeepOcean4_ |
| Marinisomatota | DeepOcean4_ |
| Proteobacteria | DeepOcean4_ |
| Proteobacteria | DeepOcean4_ |
| Proteobacteria | DeepOcean4_ |
| Chloroflexota | DeepOcean4_ |
| Proteobacteria | DeepOcean4_ |
| Proteobacteria | DeepOcean4_ |
| Proteobacteria | DeepOcean4_ |
| Patescibacteria | DeepOcean4_ |
| Proteobacteria | DeepOcean4_ |
| Proteobacteria | DeepOcean4_ |
| Proteobacteria | DeepOcean4_ |
| Marinisomatota | DeepOcean4_ |
| Acidobacteriota | DeepOcean4_ |
| Proteobacteria | DeepOcean4_ |
| Proteobacteria | DeepOcean4_ |
| Marinisomatota | DeepOcean4_ |
| Proteobacteria | DeepOcean4_ |
| SAR324 | DeepOcean4_ |
| Bacteroidota | DeepOcean4_ |
| Proteobacteria | DeepOcean4_ |
| Proteobacteria | DeepOcean4_ |
| Proteobacteria | DeepOcean4_ |
| Proteobacteria | DeepOcean4_ |
| Proteobacteria | DeepOcean4_ |
| Proteobacteria | DeepOcean4_ |
| Proteobacteria | DeepOcean4_ |
| Proteobacteria | DeepOcean4_ |

| | |
|---|---|
| Proteobacteria | DeepOcean4_ |
| Proteobacteria | DeepOcean4_ |
| Proteobacteria | DeepOcean4_ |
| Bdellovibrionota | DeepOcean4_ |
| Patescibacteria | DeepOcean4_ |
| Myxococcota | DeepOcean4_ |
| Thermoplasmatota | DeepOcean4_ |
| Proteobacteria | DeepOcean4_ |
| Proteobacteria | DeepOcean4_ |
| Actinobacteriota | DeepOcean4_ |
| Verrucomicrobiota | DeepOcean4_ |
| Patescibacteria | DeepOcean4_ |
| Proteobacteria | DeepOcean4_ |
| Proteobacteria | DeepOcean4_ |
| Proteobacteria | DeepOcean4_ |
| Proteobacteria | DeepOcean4_ |
| Actinobacteriota | DeepOcean4_ |
| Chloroflexota | DeepOcean4_ |
| Myxococcota | DeepOcean4_ |
| Proteobacteria | DeepOcean4_ |
| Proteobacteria | DeepOcean4_ |
| Proteobacteria | DeepOcean4_ |
| Proteobacteria | DeepOcean4_ |
| Gemmatimonadota | DeepOcean4_ |
| Chloroflexota | DeepOcean4_ |
| Proteobacteria | DeepOcean4_ |
| Proteobacteria | DeepOcean4_ |
| Proteobacteria | DeepOcean4_ |
| Proteobacteria | DeepOcean4_ |
| Bdellovibrionota | DeepOcean4_ |
| Proteobacteria | DeepOcean4_ |
| Proteobacteria | DeepOcean4_ |
| Proteobacteria | DeepOcean4_ |
| Actinobacteriota | DeepOcean4_ |
| Proteobacteria | DeepOcean4_ |
| Actinobacteriota | DeepOcean4_ |
| Patescibacteria | DeepOcean4_ |
| Proteobacteria | DeepOcean4_ |
| Proteobacteria | DeepOcean4_ |
| Proteobacteria | DeepOcean4_ |
| Proteobacteria | DeepOcean4_ |
| Chloroflexota | DeepOcean4_ |
| Proteobacteria | DeepOcean4_ |

| | |
|---|---|
| Proteobacteria | DeepOcean4_ |
| Actinobacteriota | DeepOcean4_ |
| Proteobacteria | DeepOcean4_ |
| Proteobacteria | DeepOcean4_ |
| Actinobacteriota | DeepOcean4_ |
| Proteobacteria | DeepOcean4_ |
| Actinobacteriota | DeepOcean4_ |
| Proteobacteria | DeepOcean4_ |
| Proteobacteria | DeepOcean4_ |
| Proteobacteria | DeepOcean4_ |
| Chloroflexota | DeepOcean4_ |
| Proteobacteria | DeepOcean4_ |
| Actinobacteriota | DeepOcean4_ |
| Proteobacteria | DeepOcean4_ |
| Chloroflexota | DeepOcean4_ |
| Proteobacteria | DeepOcean4_ |
| Proteobacteria | DeepOcean4_ |
| Bacteroidota | DeepOcean4_ |
| Actinobacteriota | DeepOcean4_ |
| Proteobacteria | DeepOcean4_ |
| Proteobacteria | DeepOcean4_ |
| Bacteroidota | DeepOcean4_ |
| Chloroflexota | DeepOcean4_ |
| Proteobacteria | DeepOcean4_ |
| Proteobacteria | DeepOcean4_ |
| Proteobacteria | DeepOcean4_ |
| Proteobacteria | DeepOcean4_ |
| Proteobacteria | DeepOcean4_ |
| Proteobacteria | DeepOcean4_ |
| Proteobacteria | DeepOcean4_ |
| Chloroflexota | DeepOcean4_ |
| Proteobacteria | DeepOcean4_ |
| Proteobacteria | DeepOcean4_ |
| Proteobacteria | DeepOcean4_ |
| Proteobacteria | DeepOcean4_ |
| Chloroflexota | DeepOcean4_ |
| Proteobacteria | DeepOcean4_ |
| Proteobacteria | DeepOcean4_ |
| Bacteroidota | DeepOcean4_ |
| Proteobacteria | DeepOcean4_ |
| Proteobacteria | DeepOcean4_ |
| Dadabacteria | DeepOcean4_ |
| Proteobacteria | DeepOcean4_ |

| | |
|---|---|
| Proteobacteria | DeepOcean4_ |
| Proteobacteria | DeepOcean4_ |
| Chloroflexota | DeepOcean4_ |
| Proteobacteria | DeepOcean4_ |
| Proteobacteria | DeepOcean4_ |
| Proteobacteria | DeepOcean4_ |
| Actinobacteriota | DeepOcean4_ |
| Proteobacteria | DeepOcean4_ |
| Proteobacteria | DeepOcean4_ |
| Proteobacteria | DeepOcean4_ |
| Proteobacteria | DeepOcean4_ |
| Chloroflexota | DeepOcean4_ |
| Proteobacteria | DeepOcean4_ |
| Proteobacteria | DeepOcean4_ |
| Chloroflexota | DeepOcean4_ |
| Proteobacteria | DeepOcean4_ |
| Chloroflexota | DeepOcean4_ |
| SAR324 | DeepOcean4_ |
| Proteobacteria | DeepOcean4_ |
| Chloroflexota | DeepOcean4_ |
| Bacteroidota | DeepOcean4_ |
| Myxococcota | DeepOcean4_ |
| Proteobacteria | DeepOcean4_ |
| Verrucomicrobiota | DeepOcean4_ |
| Proteobacteria | DeepOcean4_ |
| Proteobacteria | DeepOcean4_ |
| Thermoplasmatota | DeepOcean4_ |
| Bdellovibrionota | DeepOcean4_ |
| Proteobacteria | DeepOcean4_ |
| Patescibacteria | DeepOcean4_ |
| Verrucomicrobiota_A | DeepOcean4_ |
| SAR324 | DeepOcean4_ |
| Proteobacteria | DeepOcean4_ |
| Proteobacteria | DeepOcean4_ |
| Gemmatimonadota | DeepOcean4_ |
| Proteobacteria | DeepOcean4_ |
| Bacteroidota | DeepOcean4_ |
| Proteobacteria | DeepOcean4_ |
| Patescibacteria | DeepOcean4_ |
| Proteobacteria | DeepOcean4_ |
| Patescibacteria | DeepOcean4_ |
| Actinobacteriota | DeepOcean4_ |
| Proteobacteria | DeepOcean4_ |

| | |
|---|---|
| Proteobacteria | DeepOcean4_ |
| Proteobacteria | DeepOcean4_ |
| Proteobacteria | DeepOcean4_ |
| Proteobacteria | DeepOcean4_ |
| Crenarchaeota | DeepOcean4_ |
| Bacteroidota | DeepOcean4_ |
| Proteobacteria | DeepOcean4_ |
| Bacteroidota | DeepOcean4_ |
| Proteobacteria | DeepOcean4_ |
| Proteobacteria | DeepOcean4_ |
| Proteobacteria | DeepOcean4_ |
| Proteobacteria | DeepOcean4_ |
| Proteobacteria | DeepOcean4_ |
| Proteobacteria | DeepOcean4_ |
| Proteobacteria | DeepOcean4_ |
| Proteobacteria | DeepOcean4_ |
| Proteobacteria | DeepOcean4_ |
| Proteobacteria | DeepOcean4_ |
| Proteobacteria | DeepOcean4_ |
| Proteobacteria | DeepOcean4_ |
| Bacteroidota | DeepOcean4_ |
| Proteobacteria | DeepOcean4_ |
| UBA8248 | DeepOcean4_ |
| Proteobacteria | DeepOcean4_ |
| Proteobacteria | DeepOcean4_ |
| Proteobacteria | DeepOcean4_ |
| Proteobacteria | DeepOcean4_ |
| Marinisomatota | DeepOcean4_ |
| Proteobacteria | DeepOcean4_ |
| Chloroflexota | DeepOcean4_ |
| Proteobacteria | DeepOcean4_ |
| Proteobacteria | DeepOcean4_ |
| Bacteroidota | DeepOcean4_ |
| Proteobacteria | DeepOcean4_ |
| Proteobacteria | DeepOcean4_ |
| Actinobacteriota | DeepOcean4_ |
| Proteobacteria | DeepOcean4_ |
| Proteobacteria | DeepOcean4_ |
| Bacteroidota | DeepOcean4_ |
| Verrucomicrobiota | DeepOcean4_ |
| Crenarchaeota | DeepOcean4_ |
| Proteobacteria | DeepOcean4_ |
| Proteobacteria | DeepOcean4_ |

| | |
|---|---|
| Proteobacteria | DeepOcean4_ |
| Chloroflexota | DeepOcean4_ |
| Halobacterota | DeepOcean4_ |
| Proteobacteria | DeepOcean4_ |

| Radiosensitiv | Missing Pfam | Dataset | Source |
|---|---|---|---|
| Yes | 0 | NIH_ATCC | https://doi.org/10.1186/s40168-019-0627-4 |
| Yes | 0 | NIH_ATCC | https://doi.org/10.1186/s40168-019-0627-4 |
| Yes | 0 | NIH_ATCC | https://doi.org/10.1186/s40168-019-0627-4 |
| Yes | 0 | NIH_ATCC | https://doi.org/10.1186/s40168-019-0627-4 |
| Yes | 0 | NIH_ATCC | https://doi.org/10.1186/s40168-019-0627-4 |
| Yes | 0 | NIH_ATCC | https://doi.org/10.1186/s40168-019-0627-4 |
| Yes | 1 | NIH_ATCC | https://doi.org/10.1186/s40168-019-0627-4 |
| Yes | 0 | NIH_ATCC | https://doi.org/10.1186/s40168-019-0627-4 |
| Yes | 0 | NIH_ATCC | https://doi.org/10.1186/s40168-019-0627-4 |
| Yes | 0 | NIH_ATCC | https://doi.org/10.1186/s40168-019-0627-4 |
| No | 0 | NIH_ATCC | https://doi.org/10.1186/s40168-019-0627-4 |
| No | 0 | NIH_ATCC | https://doi.org/10.1186/s40168-019-0627-4 |
| No | 0 | NIH_ATCC | https://doi.org/10.1186/s40168-019-0627-4 |
| No | 0 | NIH_ATCC | https://doi.org/10.1186/s40168-019-0627-4 |
| No | 1 | NIH_ATCC | https://doi.org/10.1186/s40168-019-0627-4 |
| No | 0 | NIH_ATCC | https://doi.org/10.1186/s40168-019-0627-4 |
| No | 0 | NIH_ATCC | https://doi.org/10.1186/s40168-019-0627-4 |
| No | 0 | NIH_ATCC | https://doi.org/10.1186/s40168-019-0627-4 |
| No | 0 | NIH_ATCC | https://doi.org/10.1186/s40168-019-0627-4 |
| No | 0 | NIH_ATCC | https://doi.org/10.1186/s40168-019-0627-4 |
| No | 0 | NIH_ATCC | https://doi.org/10.1186/s40168-019-0627-4 |
| No | 0 | NIH_ATCC | https://doi.org/10.1186/s40168-019-0627-4 |
| No | 0 | NIH_ATCC | https://doi.org/10.1186/s40168-019-0627-4 |
| No | 0 | NIH_ATCC | https://doi.org/10.1186/s40168-019-0627-4 |
| No | 0 | NIH_ATCC | https://doi.org/10.1186/s40168-019-0627-4 |
| No | 0 | NIH_ATCC | https://doi.org/10.1186/s40168-019-0627-4 |
| No | 0 | NIH_ATCC | https://doi.org/10.1186/s40168-019-0627-4 |
| No | 0 | NIH_ATCC | https://doi.org/10.1186/s40168-019-0627-4 |
| No | 1 | NIH_ATCC | https://doi.org/10.1186/s40168-019-0627-4 |
| No | 1 | NIH_ATCC | https://doi.org/10.1186/s40168-019-0627-4 |
| No | 0 | NIH_ATCC | https://doi.org/10.1186/s40168-019-0627-4 |
| No | 0 | NIH_ATCC | https://doi.org/10.1186/s40168-019-0627-4 |
| No | 1 | NIH_ATCC | https://doi.org/10.1186/s40168-019-0627-4 |
| No | 0 | NIH_ATCC | https://doi.org/10.1186/s40168-019-0627-4 |
| No | 0 | NIH_ATCC | https://doi.org/10.1186/s40168-019-0627-4 |
| No | 0 | NIH_ATCC | https://doi.org/10.1186/s40168-019-0627-4 |
| No | 0 | NIH_ATCC | https://doi.org/10.1186/s40168-019-0627-4 |
| Yes | 0 | Human_Oral | 10.1038/ismej.2010.30 |
| Yes | 0 | Human_Oral | 10.1038/ismej.2010.30 |

| | | | |
|---|---|---|---|
| Yes | 0 | Human_Oral | 10.1038/ismej.2010.30 |
| Yes | 1 | Human_Oral | 10.1038/ismej.2010.30 |
| Yes | 1 | Human_Oral | 10.1038/ismej.2010.30 |
| Yes | 0 | Human_Oral | 10.1038/ismej.2010.30 |
| Yes | 0 | Human_Oral | 10.1038/ismej.2010.30 |
| Yes | 0 | Human_Oral | 10.1038/ismej.2010.30 |
| Yes | 0 | Human_Oral | 10.1038/ismej.2010.30 |
| No | 2 | Human_Oral | 10.1038/ismej.2010.30 |
| No | 0 | Human_Oral | 10.1038/ismej.2010.30 |
| No | 0 | Human_Oral | 10.1038/ismej.2010.30 |
| No | 0 | Human_Oral | 10.1038/ismej.2010.30 |
| No | 0 | Human_Oral | 10.1038/ismej.2010.30 |
| No | 0 | Human_Oral | 10.1038/ismej.2010.30 |
| No | 0 | Human_Oral | 10.1038/ismej.2010.30 |
| No | 0 | Human_Oral | 10.1038/ismej.2010.30 |
| No | 0 | Human_Oral | 10.1038/ismej.2010.30 |
| No | 0 | Human_Oral | 10.1038/ismej.2010.30 |
| No | 0 | Human_Oral | 10.1038/ismej.2010.30 |
| No | 0 | Human_Oral | 10.1038/ismej.2010.30 |
| No | 0 | Human_Oral | 10.1038/ismej.2010.30 |
| No | 0 | Human_Oral | 10.1038/ismej.2010.30 |
| No | 1 | Human_Oral | 10.1038/ismej.2010.30 |
| No | 0 | Human_Oral | 10.1038/ismej.2010.30 |
| No | 0 | Human_Oral | 10.1038/ismej.2010.30 |
| No | 0 | Human_Oral | 10.1038/ismej.2010.30 |
| No | 0 | Human_Oral | 10.1038/ismej.2010.30 |
| No | 1 | Human_Oral | 10.1038/ismej.2010.30 |
| No | 0 | Human_Oral | 10.1038/ismej.2010.30 |
| No | 1 | Human_Oral | 10.1038/ismej.2010.30 |
| No | 2 | Human_Oral | 10.1038/ismej.2010.30 |
| No | 0 | Human_Oral | 10.1038/ismej.2010.30 |
| No | 0 | Human_Oral | 10.1038/ismej.2010.30 |
| No | 0 | Human_Oral | 10.1038/ismej.2010.30 |
| No | 0 | Human_Oral | 10.1038/ismej.2010.30 |
| No | 0 | Human_Oral | 10.1038/ismej.2010.30 |
| No | 0 | Human_Oral | 10.1038/ismej.2010.30 |
| No | 1 | Human_Oral | 10.1038/ismej.2010.30 |
| No | 1 | Human_Oral | 10.1038/ismej.2010.30 |
| No | 0 | Human_Oral | 10.1038/ismej.2010.30 |
| No | 0 | Human_Oral | 10.1038/ismej.2010.30 |
| No | 0 | Human_Oral | 10.1038/ismej.2010.30 |
| No | 1 | Human_Oral | 10.1038/ismej.2010.30 |
| No | 0 | Human_Oral | 10.1038/ismej.2010.30 |

| | | | |
|---|---|---|---|
| No | 0 | Human_Oral | 10.1038/ismej.2010.30 |
| Yes | 0 | Skin | 10.1038/nature13786 |
| Yes | 1 | Skin | 10.1038/nature13786 |
| Yes | 0 | Skin | 10.1038/nature13786 |
| Yes | 0 | Skin | https://doi.org/10.1038/nrmicro.2017.157 |
| No | 0 | Skin | 10.1038/nature13786 |
| No | 0 | Skin | https://doi.org/10.1038/nrmicro.2017.157 |
| No | 0 | Skin | https://doi.org/10.1038/nrmicro.2017.157 |
| No | 0 | Skin | 10.1038/nature13786 |
| No | 0 | Skin | https://doi.org/10.1038/nrmicro.2017.157 |
| No | 0 | Skin | https://doi.org/10.1038/nrmicro.2017.157 |
| No | 0 | Skin | https://doi.org/10.1038/nrmicro.2017.157 |
| No | 0 | Skin | https://doi.org/10.1038/nrmicro.2017.157 |
| No | 0 | Skin | https://doi.org/10.1038/nrmicro.2017.157 |
| No | 0 | Skin | 10.1038/nature13786 |
| No | 0 | Skin | 10.1038/nature13786 |
| No | 0 | Skin | 10.1038/nature13786 |
| No | 0 | Skin | https://doi.org/10.1038/nrmicro.2017.157 |
| No | 1 | Skin | 10.1038/nature13786 |
| No | 0 | Skin | https://doi.org/10.1038/nrmicro.2017.157 |
| No | 0 | Skin | https://doi.org/10.1038/nrmicro.2017.157 |
| No | 0 | Skin | https://doi.org/10.1038/nrmicro.2017.157 |
| No | 0 | Skin | https://doi.org/10.1038/nrmicro.2017.157 |
| No | 0 | Skin | https://doi.org/10.1038/nrmicro.2017.157 |
| No | 0 | Skin | https://doi.org/10.1038/nrmicro.2017.157 |
| No | 0 | Skin | 10.1038/nature13786 |
| No | 0 | Skin | 10.1038/nature13786 |
| No | 0 | Skin | https://doi.org/10.1038/nrmicro.2017.157 |
| No | 0 | Skin | 10.1038/nature13786 |
| Yes | 0 | Gut | https://doi.org/10.1038/s41587-018-0009-7 |
| Yes | 0 | Gut | https://doi.org/10.1038/s41587-018-0009-7 |
| Yes | 0 | Gut | https://doi.org/10.1038/s41587-018-0009-7 |
| Yes | 0 | Gut | https://doi.org/10.1038/s41587-018-0009-7 |
| Yes | 0 | Gut | https://doi.org/10.1038/s41587-018-0009-7 |
| Yes | 0 | Gut | https://doi.org/10.1038/s41587-018-0009-7 |
| Yes | 2 | Gut | https://doi.org/10.1038/s41587-018-0009-7 |
| Yes | 0 | Gut | https://doi.org/10.1038/s41587-018-0009-7 |
| Yes | 0 | Gut | https://doi.org/10.1038/s41587-018-0009-7 |
| Yes | 0 | Gut | https://doi.org/10.1038/s41587-018-0009-7 |
| Yes | 0 | Gut | https://doi.org/10.1038/s41587-018-0009-7 |
| No | 0 | Gut | https://doi.org/10.1038/s41587-018-0009-7 |

| | | | |
|---|---|---|---|
| No | 0 | Gut | https://doi.org/10.1038/s41587-018-0009-7 |
| No | 0 | Gut | https://doi.org/10.1038/s41587-018-0009-7 |
| No | 0 | Gut | https://doi.org/10.1038/s41587-018-0009-7 |
| No | 0 | Gut | https://doi.org/10.1038/s41587-018-0009-7 |
| No | 0 | Gut | https://doi.org/10.1038/s41587-018-0009-7 |
| No | 0 | Gut | https://doi.org/10.1038/s41587-018-0009-7 |
| No | 0 | Gut | https://doi.org/10.1038/s41587-018-0009-7 |
| No | 0 | Gut | https://doi.org/10.1038/s41587-018-0009-7 |
| No | 0 | Gut | https://doi.org/10.1038/s41587-018-0009-7 |
| No | 0 | Gut | https://doi.org/10.1038/s41587-018-0009-7 |
| No | 0 | Gut | https://doi.org/10.1038/s41587-018-0009-7 |
| No | 0 | Gut | https://doi.org/10.1038/s41587-018-0009-7 |
| No | 0 | Gut | https://doi.org/10.1038/s41587-018-0009-7 |
| No | 0 | Gut | https://doi.org/10.1038/s41587-018-0009-7 |
| No | 1 | Gut | https://doi.org/10.1038/s41587-018-0009-7 |
| No | 0 | Gut | https://doi.org/10.1038/s41587-018-0009-7 |
| No | 0 | Gut | https://doi.org/10.1038/s41587-018-0009-7 |
| No | 0 | Gut | https://doi.org/10.1038/s41587-018-0009-7 |
| No | 0 | Gut | https://doi.org/10.1038/s41587-018-0009-7 |
| No | 0 | Gut | https://doi.org/10.1038/s41587-018-0009-7 |
| No | 0 | Gut | https://doi.org/10.1038/s41587-018-0009-7 |
| No | 0 | Gut | https://doi.org/10.1038/s41587-018-0009-7 |
| No | 0 | Gut | https://doi.org/10.1038/s41587-018-0009-7 |
| No | 0 | Gut | https://doi.org/10.1038/s41587-018-0009-7 |
| No | 0 | Gut | https://doi.org/10.1038/s41587-018-0009-7 |
| No | 0 | Gut | https://doi.org/10.1038/s41587-018-0009-7 |
| No | 0 | Gut | https://doi.org/10.1038/s41587-018-0009-7 |
| No | 0 | Gut | https://doi.org/10.1038/s41587-018-0009-7 |
| No | 0 | Gut | https://doi.org/10.1038/s41587-018-0009-7 |

| **Radiosensitiv** | **Missing Pfam** | **Dataset** | **Source** | **Completeness** | **MAG_ID** |
|---|---|---|---|---|---|
| Yes | 0 | CHA | 10.1128/mSy | 77.54 | MAG_16_RE.TSV |
| Yes | 0 | CHA | 10.1128/mSy | 98.28 | MAG_4_RE.TSV |
| No | 0 | CHA | 10.1128/mSy | 89.02 | MAG_10_RE.TSV |
| No | 0 | CHA | 10.1128/mSy | 87.86 | MAG_11_RE.TSV |
| No | 0 | CHA | 10.1128/mSy | 84.94 | MAG_12_RE.TSV |
| No | 0 | CHA | 10.1128/mSy | 80.28 | MAG_13_RE.TSV |
| No | 0 | CHA | 10.1128/mSy | 79.43 | MAG_14_RE.TSV |
| No | 0 | CHA | 10.1128/mSy | 78.56 | MAG_15_RE.TSV |
| No | 0 | CHA | 10.1128/mSy | 74.65 | MAG_17_RE.TSV |
| No | 0 | CHA | 10.1128/mSy | 73.33 | MAG_18_RE.TSV |
| No | 0 | CHA | 10.1128/mSy | 72.78 | MAG_19_RE.TSV |

| | | | | | |
|---|---|---|---|---|---|
| No | 0 | CHA | 10.1128/mSy | 99.45 | MAG_2_RE.TSV |
| No | 0 | CHA | 10.1128/mSy | 72.38 | MAG_20_RE.TSV |
| No | 0 | CHA | 10.1128/mSy | 69.53 | MAG_21_RE.TSV |
| No | 1 | CHA | 10.1128/mSy | 69.41 | MAG_22_RE.TSV |
| No | 0 | CHA | 10.1128/mSy | 68.46 | MAG_23_RE.TSV |
| No | 0 | CHA | 10.1128/mSy | 67.43 | MAG_24_RE.TSV |
| No | 0 | CHA | 10.1128/mSy | 62.24 | MAG_25_RE.TSV |
| No | 2 | CHA | 10.1128/mSy | 61.92 | MAG_26_RE.TSV |
| No | 0 | CHA | 10.1128/mSy | 61.79 | MAG_27_RE.TSV |
| No | 0 | CHA | 10.1128/mSy | 59.34 | MAG_28_RE.TSV |
| No | 0 | CHA | 10.1128/mSy | 55.17 | MAG_29_RE.TSV |
| No | 0 | CHA | 10.1128/mSy | 99.15 | MAG_3_RE.TSV |
| No | 1 | CHA | 10.1128/mSy | 51.75 | MAG_30_RE.TSV |
| No | 0 | CHA | 10.1128/mSy | 51.45 | MAG_31_RE.TSV |
| No | 0 | CHA | 10.1128/mSy | 97.39 | MAG_5_RE.TSV |
| No | 0 | CHA | 10.1128/mSy | 93.54 | MAG_6_RE.TSV |
| No | 0 | CHA | 10.1128/mSy | 92.84 | MAG_7_RE.TSV |
| No | 0 | CHA | 10.1128/mSy | 92.5 | MAG_8_RE.TSV |
| No | 1 | CHA | 10.1128/mSy | 92.21 | MAG_9_RE.TSV |
| Yes | 0 | HMB | 10.1186/s401 | 99.37 | gut3-00014 |
| Yes | 0 | HMB | 10.1186/s401 | 98.73 | gut3-00011 |
| Yes | 0 | HMB | 10.1186/s401 | 76.27 | gut2-00025 |
| Yes | 0 | HMB | 10.1186/s401 | 99.26 | gut2-00023 |
| Yes | 0 | HMB | 10.1186/s401 | 97.99 | gut2-00018 |
| Yes | 0 | HMB | 10.1186/s401 | 97.4 | gut1-00017 |
| Yes | 0 | HMB | 10.1186/s401 | 100 | gut1-00015 |
| No | 0 | HMB | 10.1186/s401 | 98.9 | skin3-00007 |
| No | 0 | HMB | 10.1186/s401 | 99.71 | skin3-00008 |
| No | 0 | HMB | 10.1186/s401 | 99.45 | skin3-00006 |
| No | 0 | HMB | 10.1186/s401 | 99.75 | skin3-00005 |
| No | 0 | HMB | 10.1186/s401 | 92.15 | skin3-00002 |
| No | 0 | HMB | 10.1186/s401 | 99.81 | skin3-00004 |
| No | 0 | HMB | 10.1186/s401 | 99.96 | skin3-00003 |
| No | 0 | HMB | 10.1186/s401 | 92.5 | skin3-00001 |
| No | 0 | HMB | 10.1186/s401 | 98.13 | skin2-00015 |
| No | 0 | HMB | 10.1186/s401 | 98.8 | skin2-00013 |
| No | 0 | HMB | 10.1186/s401 | 99.71 | skin2-00012 |
| No | 0 | HMB | 10.1186/s401 | 97.99 | skin2-00014 |
| No | 0 | HMB | 10.1186/s401 | 95.75 | skin2-00011 |
| No | 0 | HMB | 10.1186/s401 | 99.81 | skin2-00010 |
| No | 0 | HMB | 10.1186/s401 | 97.99 | skin2-00007 |
| No | 0 | HMB | 10.1186/s401 | 97.13 | skin2-00009 |

| | | | | | |
|---|---|---|---|---|---|
| No | 0 | HMB | 10.1186/s401 | 99.63 | skin2-00008 |
| No | 0 | HMB | 10.1186/s401 | 94.72 | skin2-00005 |
| No | 0 | HMB | 10.1186/s401 | 95.98 | skin2-00004 |
| No | 0 | HMB | 10.1186/s401 | 92.57 | skin2-00001 |
| No | 0 | HMB | 10.1186/s401 | 98.47 | skin1-00009 |
| No | 0 | HMB | 10.1186/s401 | 89.51 | skin2-00003 |
| No | 0 | HMB | 10.1186/s401 | 96.55 | skin2-00002 |
| No | 0 | HMB | 10.1186/s401 | 89.86 | skin1-00008 |
| No | 0 | HMB | 10.1186/s401 | 99.45 | skin1-00006 |
| No | 0 | HMB | 10.1186/s401 | 99.45 | skin1-00005 |
| No | 0 | HMB | 10.1186/s401 | 99.71 | skin1-00007 |
| No | 0 | HMB | 10.1186/s401 | 98.61 | skin1-00002 |
| No | 0 | HMB | 10.1186/s401 | 99.14 | skin1-00004 |
| No | 0 | HMB | 10.1186/s401 | 97.7 | skin1-00001 |
| No | 0 | HMB | 10.1186/s401 | 99.37 | gut3-00019 |
| No | 0 | HMB | 10.1186/s401 | 93.25 | gut3-00018 |
| No | 0 | HMB | 10.1186/s401 | 91.33 | gut3-00016 |
| No | 0 | HMB | 10.1186/s401 | 95.58 | gut3-00017 |
| No | 0 | HMB | 10.1186/s401 | 99.37 | gut3-00013 |
| No | 0 | HMB | 10.1186/s401 | 99.42 | gut3-00012 |
| No | 0 | HMB | 10.1186/s401 | 52.16 | gut3-00009 |
| No | 0 | HMB | 10.1186/s401 | 96.41 | gut3-00010 |
| No | 0 | HMB | 10.1186/s401 | 99.96 | gut3-00008 |
| No | 0 | HMB | 10.1186/s401 | 60.78 | gut3-00006 |
| No | 1 | HMB | 10.1186/s401 | 54.96 | gut3-00007 |
| No | 0 | HMB | 10.1186/s401 | 77.27 | gut3-00003 |
| No | 0 | HMB | 10.1186/s401 | 82.91 | gut3-00005 |
| No | 0 | HMB | 10.1186/s401 | 96.84 | gut3-00002 |
| No | 0 | HMB | 10.1186/s401 | 98.95 | gut2-00029 |
| No | 0 | HMB | 10.1186/s401 | 99.12 | gut2-00028 |
| No | 1 | HMB | 10.1186/s401 | 58.01 | gut3-00001 |
| No | 0 | HMB | 10.1186/s401 | 93.26 | gut2-00026 |
| No | 0 | HMB | 10.1186/s401 | 100 | gut2-00022 |
| No | 0 | HMB | 10.1186/s401 | 99.33 | gut2-00019 |
| No | 0 | HMB | 10.1186/s401 | 99.33 | gut2-00021 |
| No | 0 | HMB | 10.1186/s401 | 99.37 | gut2-00020 |
| No | 0 | HMB | 10.1186/s401 | 99.42 | gut2-00017 |
| No | 0 | HMB | 10.1186/s401 | 99.33 | gut2-00013 |
| No | 0 | HMB | 10.1186/s401 | 99.52 | gut2-00015 |
| No | 0 | HMB | 10.1186/s401 | 97.57 | gut2-00014 |
| No | 0 | HMB | 10.1186/s401 | 95.02 | gut2-00016 |
| No | 0 | HMB | 10.1186/s401 | 58.77 | gut2-00012 |
| No | 0 | HMB | 10.1186/s401 | 97.64 | gut2-00010 |

| | | | | | |
|---|---|---|---|---|---|
| No | 0 | HMB | 10.1186/s401 | 88.22 | gut2-00009 |
| No | 0 | HMB | 10.1186/s401 | 73.91 | gut2-00007 |
| No | 0 | HMB | 10.1186/s401 | 96.54 | gut2-00008 |
| No | 0 | HMB | 10.1186/s401 | 79.31 | gut2-00006 |
| No | 0 | HMB | 10.1186/s401 | 99.7 | gut2-00005 |
| No | 0 | HMB | 10.1186/s401 | 89.26 | gut2-00003 |
| No | 0 | HMB | 10.1186/s401 | 93.22 | gut2-00004 |
| No | 0 | HMB | 10.1186/s401 | 89.18 | gut2-00002 |
| No | 0 | HMB | 10.1186/s401 | 99.33 | gut2-00001 |
| No | 0 | HMB | 10.1186/s401 | 99.35 | gut1-00020 |
| No | 0 | HMB | 10.1186/s401 | 99.37 | gut1-00019 |
| No | 0 | HMB | 10.1186/s401 | 97.27 | gut1-00018 |
| No | 0 | HMB | 10.1186/s401 | 99.33 | gut1-00016 |
| No | 0 | HMB | 10.1186/s401 | 99.37 | gut1-00014 |
| No | 0 | HMB | 10.1186/s401 | 99.42 | gut1-00013 |
| No | 0 | HMB | 10.1186/s401 | 96.55 | gut1-00011 |
| No | 0 | HMB | 10.1186/s401 | 98.95 | gut1-00012 |
| No | 0 | HMB | 10.1186/s401 | 99.25 | gut1-00010 |
| No | 0 | HMB | 10.1186/s401 | 99.67 | gut1-00009 |
| No | 0 | HMB | 10.1186/s401 | 77.85 | gut1-00006 |
| No | 0 | HMB | 10.1186/s401 | 97.08 | gut1-00008 |
| No | 0 | HMB | 10.1186/s401 | 80.06 | gut1-00007 |
| No | 0 | HMB | 10.1186/s401 | 95.32 | gut1-00005 |
| No | 0 | HMB | 10.1186/s401 | 76.69 | gut1-00004 |
| No | 0 | HMB | 10.1186/s401 | 72.52 | gut1-00003 |
| No | 0 | HMB | 10.1186/s401 | 77.99 | gut1-00001 |
| No | 1 | HMB | 10.1186/s401 | 84.19 | gut1-00002 |
| No | 0 | HMB | 10.1186/s401 | 99.94 | skin2-00006 |
| No | 0 | HMB | 10.1186/s401 | 99.96 | gut2-00011 |
| No | 0 | HMB | 10.1186/s401 | 99.49 | skin1-00003 |
| Yes | 0 | Deep)cean | https://doi.or | 95.91 | MP-DEEP_MAG-0011.ENHA |
| Yes | 0 | Deep)cean | https://doi.or | 92.68 | MP-DEEP_MAG-0013.ENHA |
| Yes | 0 | Deep)cean | https://doi.or | 92.47 | MP-DEEP_MAG-0019.ENHA |
| Yes | 2 | Deep)cean | https://doi.or | 63.12 | MP-DEEP_MAG-0020.ENHA |
| Yes | 0 | Deep)cean | https://doi.or | 53.76 | MP-DEEP_MAG-0023.ENHA |
| Yes | 0 | Deep)cean | https://doi.or | 99.05 | MP-DEEP_MAG-0027.ENHA |
| Yes | 1 | Deep)cean | https://doi.or | 92.08 | MP-DEEP_MAG-0036.ENHA |
| Yes | 0 | Deep)cean | https://doi.or | 94.35 | MP-DEEP_MAG-0037.ENHA |
| Yes | 0 | Deep)cean | https://doi.or | 78.25 | MP-DEEP_MAG-0038.ENHA |
| Yes | 0 | Deep)cean | https://doi.or | 99.29 | MP-DEEP_MAG-0053.ENHA |
| Yes | 1 | Deep)cean | https://doi.or | 69.07 | MP-DEEP_MAG-0055.ENHA |
| Yes | 0 | Deep)cean | https://doi.or | 93.53 | MP-DEEP_MAG-0058.ENHA |

| | | | | | |
|---|---|---|---|---|---|
| Yes | 1 | Deep)cean | https://doi.or | 96.43 | MP-DEEP_MAG-0070.ENHA |
| Yes | 1 | Deep)cean | https://doi.or | 85.16 | MP-DEEP_MAG-0077.ENHA |
| Yes | 0 | Deep)cean | https://doi.or | 84.56 | MP-DEEP_MAG-0082.ENHA |
| Yes | 2 | Deep)cean | https://doi.or | 94.17 | MP-DEEP_MAG-0093.ENHA |
| Yes | 1 | Deep)cean | https://doi.or | 91.21 | MP-DEEP_MAG-0102.ENHA |
| Yes | 0 | Deep)cean | https://doi.or | 93.55 | MP-DEEP_MAG-0108.ENHA |
| Yes | 1 | Deep)cean | https://doi.or | 88.07 | MP-DEEP_MAG-0122.ENHA |
| Yes | 2 | Deep)cean | https://doi.or | 58.1 | MP-DEEP_MAG-0123.ENHA |
| Yes | 0 | Deep)cean | https://doi.or | 96.58 | MP-DEEP_MAG-0132.ENHA |
| Yes | 3 | Deep)cean | https://doi.or | 67.37 | MP-DEEP_MAG-0142.ENHA |
| Yes | 3 | Deep)cean | https://doi.or | 61.79 | MP-DEEP_MAG-0155.ENHA |
| Yes | 0 | Deep)cean | https://doi.or | 90.79 | MP-DEEP_MAG-0165.ENHA |
| Yes | 0 | Deep)cean | https://doi.or | 93.9 | MP-DEEP_MAG-0173.ENHA |
| Yes | 1 | Deep)cean | https://doi.or | 82.43 | MP-DEEP_MAG-0176.ENHA |
| Yes | 0 | Deep)cean | https://doi.or | 99.42 | MP-DEEP_MAG-0177.ENHA |
| Yes | 2 | Deep)cean | https://doi.or | 100 | MP-DEEP_MAG-0183.ENHA |
| Yes | 2 | Deep)cean | https://doi.or | 82.97 | MP-DEEP_MAG-0191.ENHA |
| Yes | 3 | Deep)cean | https://doi.or | 77.44 | MP-DEEP_MAG-0194.ENHA |
| Yes | 0 | Deep)cean | https://doi.or | 97.26 | MP-DEEP_MAG-0195.ENHA |
| Yes | 3 | Deep)cean | https://doi.or | 72.58 | MP-DEEP_MAG-0196.ENHA |
| Yes | 1 | Deep)cean | https://doi.or | 87.91 | MP-DEEP_MAG-0200.ENHA |
| Yes | 1 | Deep)cean | https://doi.or | 91.21 | MP-DEEP_MAG-0203.ENHA |
| Yes | 0 | Deep)cean | https://doi.or | 54.48 | MP-DEEP_MAG-0208.ENHA |
| Yes | 0 | Deep)cean | https://doi.or | 87.02 | MP-DEEP_MAG-0209.ENHA |
| Yes | 0 | Deep)cean | https://doi.or | 92.79 | MP-DEEP_MAG-0213.ENHA |
| Yes | 1 | Deep)cean | https://doi.or | 88.24 | MP-DEEP_MAG-0214.ENHA |
| Yes | 1 | Deep)cean | https://doi.or | 95.6 | MP-DEEP_MAG-0218.ENHA |
| Yes | 0 | Deep)cean | https://doi.or | 92.68 | MP-DEEP_MAG-0223.ENHA |
| Yes | 2 | Deep)cean | https://doi.or | 66.47 | MP-DEEP_MAG-0231.ENHA |
| Yes | 3 | Deep)cean | https://doi.or | 82.99 | MP-DEEP_MAG-0238.ENHA |
| Yes | 2 | Deep)cean | https://doi.or | 79.21 | MP-DEEP_MAG-0241.ENHA |
| Yes | 4 | Deep)cean | https://doi.or | 78.74 | MP-DEEP_MAG-0246.ENHA |
| Yes | 1 | Deep)cean | https://doi.or | 92.01 | MP-DEEP_MAG-0255.ENHA |
| Yes | 3 | Deep)cean | https://doi.or | 57.09 | MP-DEEP_MAG-0273.ENHA |
| Yes | 0 | Deep)cean | https://doi.or | 91.42 | MP-DEEP_MAG-0283.ENHA |
| Yes | 0 | Deep)cean | https://doi.or | 90.05 | MP-DEEP_MAG-0285.ENHA |
| Yes | 3 | Deep)cean | https://doi.or | 52.81 | MP-DEEP_MAG-0294.ENHA |
| Yes | 3 | Deep)cean | https://doi.or | 75.77 | MP-DEEP_MAG-0307.ENHA |
| Yes | 1 | Deep)cean | https://doi.or | 81.75 | MP-DEEP_MAG-0318.ENHA |
| Yes | 2 | Deep)cean | https://doi.or | 65.56 | MP-DEEP_MAG-0319.ENHA |
| Yes | 0 | Deep)cean | https://doi.or | 93.96 | MP-DEEP_MAG-0321.ENHA |
| Yes | 0 | Deep)cean | https://doi.or | 91.32 | MP-DEEP_MAG-0323.ENHA |
| Yes | 0 | Deep)cean | https://doi.or | 88.7 | MP-DEEP_MAG-0330.ENHA |

| | | | | | |
|---|---|---|---|---|---|
| Yes | 0 | Deep)cean | https://doi.or | 70.52 | MP-DEEP_MAG-0335.ENHA |
| Yes | 2 | Deep)cean | https://doi.or | 90.11 | MP-DEEP_MAG-0339.ENHA |
| Yes | 1 | Deep)cean | https://doi.or | 72.81 | MP-DEEP_MAG-0340.ENHA |
| Yes | 0 | Deep)cean | https://doi.or | 86.42 | MP-DEEP_MAG-0349.ENHA |
| Yes | 0 | Deep)cean | https://doi.or | 94.17 | MP-DEEP_MAG-0351.ENHA |
| Yes | 2 | Deep)cean | https://doi.or | 57.58 | MP-DEEP_MAG-0354.ENHA |
| Yes | 0 | Deep)cean | https://doi.or | 82.65 | MP-DEEP_MAG-0355.ENHA |
| Yes | 0 | Deep)cean | https://doi.or | 86.04 | MP-DEEP_MAG-0356.ENHA |
| Yes | 1 | Deep)cean | https://doi.or | 84.07 | MP-DEEP_MAG-0363.ENHA |
| Yes | 2 | Deep)cean | https://doi.or | 80.49 | MP-DEEP_MAG-0368.ENHA |
| Yes | 0 | Deep)cean | https://doi.or | 76.92 | MP-DEEP_MAG-0373.ENHA |
| Yes | 0 | Deep)cean | https://doi.or | 93.6 | MP-DEEP_MAG-0376.ENHA |
| Yes | 0 | Deep)cean | https://doi.or | 97.53 | MP-DEEP_MAG-0378.ENHA |
| Yes | 1 | Deep)cean | https://doi.or | 81.35 | MP-DEEP_MAG-0383.ENHA |
| Yes | 2 | Deep)cean | https://doi.or | 84.49 | MP-DEEP_MAG-0391.ENHA |
| Yes | 1 | Deep)cean | https://doi.or | 66.99 | MP-DEEP_MAG-0397.ENHA |
| Yes | 1 | Deep)cean | https://doi.or | 67.26 | MP-DEEP_MAG-0401.ENHA |
| Yes | 1 | Deep)cean | https://doi.or | 71.68 | MP-DEEP_MAG-0404.ENHA |
| Yes | 0 | Deep)cean | https://doi.or | 96.77 | MP-DEEP_MAG-0410.ENHA |
| Yes | 0 | Deep)cean | https://doi.or | 93.11 | MP-DEEP_MAG-0416.ENHA |
| Yes | 0 | Deep)cean | https://doi.or | 91.03 | MP-DEEP_MAG-0434.ENHA |
| Yes | 1 | Deep)cean | https://doi.or | 72.97 | MP-DEEP_MAG-0438.ENHA |
| Yes | 1 | Deep)cean | https://doi.or | 87.91 | MP-DEEP_MAG-0447.ENHA |
| Yes | 3 | Deep)cean | https://doi.or | 85.19 | MP-DEEP_MAG-0465.ENHA |
| Yes | 1 | Deep)cean | https://doi.or | 79.06 | MP-DEEP_MAG-0471.ENHA |
| Yes | 3 | Deep)cean | https://doi.or | 51.16 | MP-DEEP_MAG-0474.ENHA |
| Yes | 0 | Deep)cean | https://doi.or | 94.44 | MP-DEEP_MAG-0478.ENHA |
| Yes | 4 | Deep)cean | https://doi.or | 50.59 | MP-DEEP_MAG-0485.ENHA |
| Yes | 0 | Deep)cean | https://doi.or | 65.67 | MP-DEEP_MAG-0488.ENHA |
| Yes | 1 | Deep)cean | https://doi.or | 98.84 | MP-DEEP_MAG-0490.ENHA |
| Yes | 0 | Deep)cean | https://doi.or | 93.55 | MP-DEEP_MAG-0491.ENHA |
| Yes | 0 | Deep)cean | https://doi.or | 99.67 | MP-DEEP_MAG-0494.ENHA |
| Yes | 0 | Deep)cean | https://doi.or | 85.2 | MP-DEEP_MAG-0496.ENHA |
| Yes | 0 | Deep)cean | https://doi.or | 71.68 | MP-DEEP_MAG-0503.ENHA |
| Yes | 2 | Deep)cean | https://doi.or | 90.32 | MP-DEEP_MAG-0514.ENHA |
| Yes | 0 | Deep)cean | https://doi.or | 96.14 | MP-DEEP_MAG-0518.ENHA |
| Yes | 1 | Deep)cean | https://doi.or | 70.53 | MP-DEEP_MAG-0519.ENHA |
| Yes | 1 | Deep)cean | https://doi.or | 50.43 | MP-DEEP_MAG-0524.ENHA |
| Yes | 3 | Deep)cean | https://doi.or | 52.61 | MP-DEEP_MAG-0526.ENHA |
| Yes | 2 | Deep)cean | https://doi.or | 78.12 | MP-DEEP_MAG-0534.ENHA |
| Yes | 0 | Deep)cean | https://doi.or | 93.87 | MP-DEEP_MAG-0539.ENHA |
| Yes | 0 | Deep)cean | https://doi.or | 95.3 | MP-DEEP_MAG-0548.ENHA |
| Yes | 3 | Deep)cean | https://doi.or | 76.34 | MP-DEEP_MAG-0549.ENHA |

| | | | | | |
|---|---|---|---|---|---|
| Yes | 3 | Deep)cean | https://doi.or | 59.01 | MP-DEEP_MAG-0569.ENHA |
| Yes | 0 | Deep)cean | https://doi.or | 86.21 | MP-DEEP_MAG-0573.ENHA |
| Yes | 3 | Deep)cean | https://doi.or | 60.06 | MP-DEEP_MAG-0576.ENHA |
| Yes | 1 | Deep)cean | https://doi.or | 53.26 | MP-DEEP_MAG-0577.ENHA |
| Yes | 0 | Deep)cean | https://doi.or | 99.46 | MP-DEEP_MAG-0578.ENHA |
| Yes | 0 | Deep)cean | https://doi.or | 98.35 | MP-DEEP_MAG-0580.ENHA |
| Yes | 1 | Deep)cean | https://doi.or | 87.94 | MP-DEEP_MAG-0585.ENHA |
| Yes | 1 | Deep)cean | https://doi.or | 96.64 | MP-DEEP_MAG-0586.ENHA |
| Yes | 2 | Deep)cean | https://doi.or | 95.54 | MP-DEEP_MAG-0590.ENHA |
| Yes | 0 | Deep)cean | https://doi.or | 99.67 | MP-DEEP_MAG-0593.ENHA |
| Yes | 0 | Deep)cean | https://doi.or | 98.88 | MP-DEEP_MAG-0603.ENHA |
| Yes | 0 | Deep)cean | https://doi.or | 99.46 | MP-DEEP_MAG-0606.ENHA |
| Yes | 0 | Deep)cean | https://doi.or | 95.5 | MP-DEEP_MAG-0608.ENHA |
| Yes | 2 | Deep)cean | https://doi.or | 86.81 | MP-DEEP_MAG-0609.ENHA |
| Yes | 0 | Deep)cean | https://doi.or | 98.58 | MP-DEEP_MAG-0612.ENHA |
| Yes | 1 | Deep)cean | https://doi.or | 87.91 | MP-DEEP_MAG-0619.ENHA |
| No | 0 | Deep)cean | https://doi.or | 65.08 | MP-DEEP_MAG-0002.ENHA |
| No | 0 | Deep)cean | https://doi.or | 76.19 | MP-DEEP_MAG-0008.ENHA |
| No | 2 | Deep)cean | https://doi.or | 55.3 | MP-DEEP_MAG-0015.ENHA |
| No | 0 | Deep)cean | https://doi.or | 72.98 | MP-DEEP_MAG-0018.ENHA |
| No | 0 | Deep)cean | https://doi.or | 96.12 | MP-DEEP_MAG-0021.ENHA |
| No | 0 | Deep)cean | https://doi.or | 97.78 | MP-DEEP_MAG-0028.ENHA |
| No | 0 | Deep)cean | https://doi.or | 95.93 | MP-DEEP_MAG-0029.ENHA |
| No | 2 | Deep)cean | https://doi.or | 64.25 | MP-DEEP_MAG-0031.ENHA |
| No | 0 | Deep)cean | https://doi.or | 91.75 | MP-DEEP_MAG-0035.ENHA |
| No | 0 | Deep)cean | https://doi.or | 52.33 | MP-DEEP_MAG-0039.ENHA |
| No | 0 | Deep)cean | https://doi.or | 92.35 | MP-DEEP_MAG-0040.ENHA |
| No | 2 | Deep)cean | https://doi.or | 57.23 | MP-DEEP_MAG-0042.ENHA |
| No | 0 | Deep)cean | https://doi.or | 90.99 | MP-DEEP_MAG-0044.ENHA |
| No | 0 | Deep)cean | https://doi.or | 81.52 | MP-DEEP_MAG-0045.ENHA |
| No | 0 | Deep)cean | https://doi.or | 86.74 | MP-DEEP_MAG-0047.ENHA |
| No | 1 | Deep)cean | https://doi.or | 86.81 | MP-DEEP_MAG-0048.ENHA |
| No | 1 | Deep)cean | https://doi.or | 84.33 | MP-DEEP_MAG-0050.ENHA |
| No | 0 | Deep)cean | https://doi.or | 91.26 | MP-DEEP_MAG-0052.ENHA |
| No | 0 | Deep)cean | https://doi.or | 95.33 | MP-DEEP_MAG-0060.ENHA |
| No | 0 | Deep)cean | https://doi.or | 85.41 | MP-DEEP_MAG-0064.ENHA |
| No | 0 | Deep)cean | https://doi.or | 88.96 | MP-DEEP_MAG-0067.ENHA |
| No | 0 | Deep)cean | https://doi.or | 98.64 | MP-DEEP_MAG-0075.ENHA |
| No | 1 | Deep)cean | https://doi.or | 56.72 | MP-DEEP_MAG-0078.ENHA |
| No | 0 | Deep)cean | https://doi.or | 97.78 | MP-DEEP_MAG-0081.ENHA |
| No | 0 | Deep)cean | https://doi.or | 98.95 | MP-DEEP_MAG-0084.ENHA |
| No | 0 | Deep)cean | https://doi.or | 98.71 | MP-DEEP_MAG-0094.ENHA |
| No | 0 | Deep)cean | https://doi.or | 84.92 | MP-DEEP_MAG-0095.ENHA |

| | | | | | |
|---|---|---|---|---|---|
| No | 0 | Deep)cean | https://doi.oı | 82.83 | MP-DEEP_MAG-0096.ENHA |
| No | 0 | Deep)cean | https://doi.oı | 84.74 | MP-DEEP_MAG-0098.ENHA |
| No | 1 | Deep)cean | https://doi.oı | 89.16 | MP-DEEP_MAG-0101.ENHA |
| No | 1 | Deep)cean | https://doi.oı | 92.56 | MP-DEEP_MAG-0103.ENHA |
| No | 3 | Deep)cean | https://doi.oı | 51.29 | MP-DEEP_MAG-0105.ENHA |
| No | 0 | Deep)cean | https://doi.oı | 68.77 | MP-DEEP_MAG-0111.ENHA |
| No | 2 | Deep)cean | https://doi.oı | 69.87 | MP-DEEP_MAG-0112.ENHA |
| No | 0 | Deep)cean | https://doi.oı | 95.99 | MP-DEEP_MAG-0118.ENHA |
| No | 0 | Deep)cean | https://doi.oı | 77.24 | MP-DEEP_MAG-0126.ENHA |
| No | 0 | Deep)cean | https://doi.oı | 87.93 | MP-DEEP_MAG-0127.ENHA |
| No | 1 | Deep)cean | https://doi.oı | 86.66 | MP-DEEP_MAG-0128.ENHA |
| No | 2 | Deep)cean | https://doi.oı | 61.05 | MP-DEEP_MAG-0130.ENHA |
| No | 2 | Deep)cean | https://doi.oı | 79.57 | MP-DEEP_MAG-0131.ENHA |
| No | 0 | Deep)cean | https://doi.oı | 95.26 | MP-DEEP_MAG-0134.ENHA |
| No | 0 | Deep)cean | https://doi.oı | 98.24 | MP-DEEP_MAG-0137.ENHA |
| No | 0 | Deep)cean | https://doi.oı | 98.99 | MP-DEEP_MAG-0140.ENHA |
| No | 0 | Deep)cean | https://doi.oı | 86.53 | MP-DEEP_MAG-0144.ENHA |
| No | 0 | Deep)cean | https://doi.oı | 89.32 | MP-DEEP_MAG-0146.ENHA |
| No | 1 | Deep)cean | https://doi.oı | 92.9 | MP-DEEP_MAG-0149.ENHA |
| No | 1 | Deep)cean | https://doi.oı | 59.74 | MP-DEEP_MAG-0150.ENHA |
| No | 0 | Deep)cean | https://doi.oı | 91.06 | MP-DEEP_MAG-0152.ENHA |
| No | 0 | Deep)cean | https://doi.oı | 98.6 | MP-DEEP_MAG-0153.ENHA |
| No | 0 | Deep)cean | https://doi.oı | 64.5 | MP-DEEP_MAG-0156.ENHA |
| No | 1 | Deep)cean | https://doi.oı | 97.74 | MP-DEEP_MAG-0161.ENHA |
| No | 0 | Deep)cean | https://doi.oı | 88.45 | MP-DEEP_MAG-0162.ENHA |
| No | 0 | Deep)cean | https://doi.oı | 99.57 | MP-DEEP_MAG-0166.ENHA |
| No | 0 | Deep)cean | https://doi.oı | 76.64 | MP-DEEP_MAG-0167.ENHA |
| No | 0 | Deep)cean | https://doi.oı | 84.5 | MP-DEEP_MAG-0169.ENHA |
| No | 0 | Deep)cean | https://doi.oı | 92.75 | MP-DEEP_MAG-0175.ENHA |
| No | 0 | Deep)cean | https://doi.oı | 88.99 | MP-DEEP_MAG-0178.ENHA |
| No | 2 | Deep)cean | https://doi.oı | 82.23 | MP-DEEP_MAG-0181.ENHA |
| No | 0 | Deep)cean | https://doi.oı | 88.7 | MP-DEEP_MAG-0182.ENHA |
| No | 0 | Deep)cean | https://doi.oı | 92.92 | MP-DEEP_MAG-0184.ENHA |
| No | 0 | Deep)cean | https://doi.oı | 94.2 | MP-DEEP_MAG-0185.ENHA |
| No | 0 | Deep)cean | https://doi.oı | 95.38 | MP-DEEP_MAG-0186.ENHA |
| No | 0 | Deep)cean | https://doi.oı | 69.17 | MP-DEEP_MAG-0188.ENHA |
| No | 2 | Deep)cean | https://doi.oı | 67.87 | MP-DEEP_MAG-0192.ENHA |
| No | 0 | Deep)cean | https://doi.oı | 72.41 | MP-DEEP_MAG-0193.ENHA |
| No | 0 | Deep)cean | https://doi.oı | 99.51 | MP-DEEP_MAG-0202.ENHA |
| No | 0 | Deep)cean | https://doi.oı | 92.09 | MP-DEEP_MAG-0210.ENHA |
| No | 0 | Deep)cean | https://doi.oı | 92.02 | MP-DEEP_MAG-0212.ENHA |
| No | 1 | Deep)cean | https://doi.oı | 95.71 | MP-DEEP_MAG-0222.ENHA |
| No | 0 | Deep)cean | https://doi.oı | 76.6 | MP-DEEP_MAG-0226.ENHA |

| | | | | | |
|---|---|---|---|---|---|
| No | 0 | Deep)cean | https://doi.or | 98.28 | MP-DEEP_MAG-0228.ENHA |
| No | 0 | Deep)cean | https://doi.or | 87.61 | MP-DEEP_MAG-0229.ENHA |
| No | 0 | Deep)cean | https://doi.or | 92.74 | MP-DEEP_MAG-0233.ENHA |
| No | 0 | Deep)cean | https://doi.or | 95.78 | MP-DEEP_MAG-0239.ENHA |
| No | 0 | Deep)cean | https://doi.or | 86.88 | MP-DEEP_MAG-0242.ENHA |
| No | 0 | Deep)cean | https://doi.or | 72.96 | MP-DEEP_MAG-0250.ENHA |
| No | 0 | Deep)cean | https://doi.or | 97.01 | MP-DEEP_MAG-0251.ENHA |
| No | 0 | Deep)cean | https://doi.or | 85.2 | MP-DEEP_MAG-0258.ENHA |
| No | 0 | Deep)cean | https://doi.or | 96.47 | MP-DEEP_MAG-0262.ENHA |
| No | 0 | Deep)cean | https://doi.or | 96.14 | MP-DEEP_MAG-0263.ENHA |
| No | 0 | Deep)cean | https://doi.or | 55.32 | MP-DEEP_MAG-0265.ENHA |
| No | 0 | Deep)cean | https://doi.or | 89.05 | MP-DEEP_MAG-0267.ENHA |
| No | 0 | Deep)cean | https://doi.or | 99.47 | MP-DEEP_MAG-0269.ENHA |
| No | 0 | Deep)cean | https://doi.or | 98.89 | MP-DEEP_MAG-0274.ENHA |
| No | 0 | Deep)cean | https://doi.or | 91.59 | MP-DEEP_MAG-0275.ENHA |
| No | 0 | Deep)cean | https://doi.or | 94.59 | MP-DEEP_MAG-0280.ENHA |
| No | 2 | Deep)cean | https://doi.or | 88.84 | MP-DEEP_MAG-0282.ENHA |
| No | 1 | Deep)cean | https://doi.or | 90.02 | MP-DEEP_MAG-0284.ENHA |
| No | 0 | Deep)cean | https://doi.or | 93.16 | MP-DEEP_MAG-0286.ENHA |
| No | 0 | Deep)cean | https://doi.or | 62.93 | MP-DEEP_MAG-0289.ENHA |
| No | 1 | Deep)cean | https://doi.or | 65.52 | MP-DEEP_MAG-0290.ENHA |
| No | 0 | Deep)cean | https://doi.or | 92.88 | MP-DEEP_MAG-0291.ENHA |
| No | 0 | Deep)cean | https://doi.or | 91.85 | MP-DEEP_MAG-0293.ENHA |
| No | 0 | Deep)cean | https://doi.or | 87.19 | MP-DEEP_MAG-0295.ENHA |
| No | 0 | Deep)cean | https://doi.or | 97.43 | MP-DEEP_MAG-0296.ENHA |
| No | 0 | Deep)cean | https://doi.or | 68.97 | MP-DEEP_MAG-0298.ENHA |
| No | 0 | Deep)cean | https://doi.or | 70.6 | MP-DEEP_MAG-0300.ENHA |
| No | 0 | Deep)cean | https://doi.or | 99.31 | MP-DEEP_MAG-0303.ENHA |
| No | 0 | Deep)cean | https://doi.or | 66.38 | MP-DEEP_MAG-0304.ENHA |
| No | 0 | Deep)cean | https://doi.or | 97.38 | MP-DEEP_MAG-0305.ENHA |
| No | 1 | Deep)cean | https://doi.or | 94.22 | MP-DEEP_MAG-0309.ENHA |
| No | 0 | Deep)cean | https://doi.or | 88.23 | MP-DEEP_MAG-0314.ENHA |
| No | 0 | Deep)cean | https://doi.or | 85.11 | MP-DEEP_MAG-0315.ENHA |
| No | 0 | Deep)cean | https://doi.or | 85.66 | MP-DEEP_MAG-0317.ENHA |
| No | 0 | Deep)cean | https://doi.or | 79.32 | MP-DEEP_MAG-0320.ENHA |
| No | 0 | Deep)cean | https://doi.or | 96.2 | MP-DEEP_MAG-0322.ENHA |
| No | 0 | Deep)cean | https://doi.or | 71.82 | MP-DEEP_MAG-0325.ENHA |
| No | 0 | Deep)cean | https://doi.or | 89.26 | MP-DEEP_MAG-0327.ENHA |
| No | 0 | Deep)cean | https://doi.or | 98.79 | MP-DEEP_MAG-0328.ENHA |
| No | 0 | Deep)cean | https://doi.or | 55.39 | MP-DEEP_MAG-0329.ENHA |
| No | 0 | Deep)cean | https://doi.or | 92.99 | MP-DEEP_MAG-0334.ENHA |
| No | 1 | Deep)cean | https://doi.or | 82.59 | MP-DEEP_MAG-0336.ENHA |
| No | 0 | Deep)cean | https://doi.or | 84.62 | MP-DEEP_MAG-0338.ENHA |

| | | | | | |
|---|---|---|---|---|---|
| No | 0 | Deep)cean | https://doi.or | 70.7 | MP-DEEP_MAG-0342.ENHA |
| No | 0 | Deep)cean | https://doi.or | 95.81 | MP-DEEP_MAG-0343.ENHA |
| No | 0 | Deep)cean | https://doi.or | 92.74 | MP-DEEP_MAG-0344.ENHA |
| No | 0 | Deep)cean | https://doi.or | 97.81 | MP-DEEP_MAG-0345.ENHA |
| No | 0 | Deep)cean | https://doi.or | 96.73 | MP-DEEP_MAG-0347.ENHA |
| No | 1 | Deep)cean | https://doi.or | 71.5 | MP-DEEP_MAG-0353.ENHA |
| No | 0 | Deep)cean | https://doi.or | 86.87 | MP-DEEP_MAG-0358.ENHA |
| No | 0 | Deep)cean | https://doi.or | 80.17 | MP-DEEP_MAG-0361.ENHA |
| No | 0 | Deep)cean | https://doi.or | 92.05 | MP-DEEP_MAG-0362.ENHA |
| No | 0 | Deep)cean | https://doi.or | 99.57 | MP-DEEP_MAG-0366.ENHA |
| No | 0 | Deep)cean | https://doi.or | 97.04 | MP-DEEP_MAG-0367.ENHA |
| No | 0 | Deep)cean | https://doi.or | 82.21 | MP-DEEP_MAG-0375.ENHA |
| No | 0 | Deep)cean | https://doi.or | 59.64 | MP-DEEP_MAG-0380.ENHA |
| No | 0 | Deep)cean | https://doi.or | 98.91 | MP-DEEP_MAG-0386.ENHA |
| No | 0 | Deep)cean | https://doi.or | 57.24 | MP-DEEP_MAG-0387.ENHA |
| No | 0 | Deep)cean | https://doi.or | 85.44 | MP-DEEP_MAG-0390.ENHA |
| No | 1 | Deep)cean | https://doi.or | 91.75 | MP-DEEP_MAG-0399.ENHA |
| No | 1 | Deep)cean | https://doi.or | 85.33 | MP-DEEP_MAG-0400.ENHA |
| No | 1 | Deep)cean | https://doi.or | 78.66 | MP-DEEP_MAG-0405.ENHA |
| No | 0 | Deep)cean | https://doi.or | 53.03 | MP-DEEP_MAG-0419.ENHA |
| No | 0 | Deep)cean | https://doi.or | 99.01 | MP-DEEP_MAG-0420.ENHA |
| No | 0 | Deep)cean | https://doi.or | 95.48 | MP-DEEP_MAG-0421.ENHA |
| No | 0 | Deep)cean | https://doi.or | 95.12 | MP-DEEP_MAG-0422.ENHA |
| No | 2 | Deep)cean | https://doi.or | 65.94 | MP-DEEP_MAG-0427.ENHA |
| No | 0 | Deep)cean | https://doi.or | 91.66 | MP-DEEP_MAG-0430.ENHA |
| No | 0 | Deep)cean | https://doi.or | 95.29 | MP-DEEP_MAG-0435.ENHA |
| No | 2 | Deep)cean | https://doi.or | 71.47 | MP-DEEP_MAG-0437.ENHA |
| No | 0 | Deep)cean | https://doi.or | 93.45 | MP-DEEP_MAG-0439.ENHA |
| No | 1 | Deep)cean | https://doi.or | 96.87 | MP-DEEP_MAG-0441.ENHA |
| No | 2 | Deep)cean | https://doi.or | 63.14 | MP-DEEP_MAG-0449.ENHA |
| No | 1 | Deep)cean | https://doi.or | 92.57 | MP-DEEP_MAG-0452.ENHA |
| No | 1 | Deep)cean | https://doi.or | 89.58 | MP-DEEP_MAG-0455.ENHA |
| No | 0 | Deep)cean | https://doi.or | 94 | MP-DEEP_MAG-0457.ENHA |
| No | 0 | Deep)cean | https://doi.or | 84.31 | MP-DEEP_MAG-0459.ENHA |
| No | 1 | Deep)cean | https://doi.or | 96.7 | MP-DEEP_MAG-0461.ENHA |
| No | 0 | Deep)cean | https://doi.or | 86.91 | MP-DEEP_MAG-0466.ENHA |
| No | 0 | Deep)cean | https://doi.or | 99.34 | MP-DEEP_MAG-0467.ENHA |
| No | 0 | Deep)cean | https://doi.or | 60.53 | MP-DEEP_MAG-0469.ENHA |
| No | 2 | Deep)cean | https://doi.or | 63.14 | MP-DEEP_MAG-0470.ENHA |
| No | 1 | Deep)cean | https://doi.or | 69.03 | MP-DEEP_MAG-0472.ENHA |
| No | 2 | Deep)cean | https://doi.or | 80.3 | MP-DEEP_MAG-0477.ENHA |
| No | 0 | Deep)cean | https://doi.or | 92.73 | MP-DEEP_MAG-0480.ENHA |
| No | 0 | Deep)cean | https://doi.or | 97.8 | MP-DEEP_MAG-0483.ENHA |

| | | | | | |
|---|---|---|---|---|---|
| No | 0 | Deep)cean | https://doi.or | 93.79 | MP-DEEP_MAG-0484.ENHA |
| No | 0 | Deep)cean | https://doi.or | 92.39 | MP-DEEP_MAG-0489.ENHA |
| No | 0 | Deep)cean | https://doi.or | 93.35 | MP-DEEP_MAG-0502.ENHA |
| No | 0 | Deep)cean | https://doi.or | 79.75 | MP-DEEP_MAG-0505.ENHA |
| No | 2 | Deep)cean | https://doi.or | 100 | MP-DEEP_MAG-0507.ENHA |
| No | 0 | Deep)cean | https://doi.or | 65.52 | MP-DEEP_MAG-0508.ENHA |
| No | 0 | Deep)cean | https://doi.or | 94.83 | MP-DEEP_MAG-0509.ENHA |
| No | 1 | Deep)cean | https://doi.or | 75.98 | MP-DEEP_MAG-0510.ENHA |
| No | 0 | Deep)cean | https://doi.or | 97.41 | MP-DEEP_MAG-0511.ENHA |
| No | 0 | Deep)cean | https://doi.or | 97.75 | MP-DEEP_MAG-0512.ENHA |
| No | 1 | Deep)cean | https://doi.or | 88.4 | MP-DEEP_MAG-0523.ENHA |
| No | 0 | Deep)cean | https://doi.or | 99.46 | MP-DEEP_MAG-0527.ENHA |
| No | 0 | Deep)cean | https://doi.or | 99.7 | MP-DEEP_MAG-0528.ENHA |
| No | 0 | Deep)cean | https://doi.or | 98.38 | MP-DEEP_MAG-0531.ENHA |
| No | 0 | Deep)cean | https://doi.or | 96.97 | MP-DEEP_MAG-0533.ENHA |
| No | 0 | Deep)cean | https://doi.or | 91.46 | MP-DEEP_MAG-0535.ENHA |
| No | 0 | Deep)cean | https://doi.or | 92.07 | MP-DEEP_MAG-0542.ENHA |
| No | 0 | Deep)cean | https://doi.or | 94.46 | MP-DEEP_MAG-0545.ENHA |
| No | 0 | Deep)cean | https://doi.or | 98.15 | MP-DEEP_MAG-0553.ENHA |
| No | 0 | Deep)cean | https://doi.or | 67.05 | MP-DEEP_MAG-0554.ENHA |
| No | 0 | Deep)cean | https://doi.or | 95.73 | MP-DEEP_MAG-0555.ENHA |
| No | 0 | Deep)cean | https://doi.or | 76.57 | MP-DEEP_MAG-0557.ENHA |
| No | 1 | Deep)cean | https://doi.or | 56.41 | MP-DEEP_MAG-0559.ENHA |
| No | 0 | Deep)cean | https://doi.or | 91.09 | MP-DEEP_MAG-0560.ENHA |
| No | 0 | Deep)cean | https://doi.or | 90.96 | MP-DEEP_MAG-0562.ENHA |
| No | 0 | Deep)cean | https://doi.or | 93.74 | MP-DEEP_MAG-0564.ENHA |
| No | 0 | Deep)cean | https://doi.or | 62.47 | MP-DEEP_MAG-0566.ENHA |
| No | 3 | Deep)cean | https://doi.or | 57.48 | MP-DEEP_MAG-0568.ENHA |
| No | 1 | Deep)cean | https://doi.or | 77.49 | MP-DEEP_MAG-0570.ENHA |
| No | 1 | Deep)cean | https://doi.or | 91.75 | MP-DEEP_MAG-0571.ENHA |
| No | 0 | Deep)cean | https://doi.or | 95.62 | MP-DEEP_MAG-0572.ENHA |
| No | 0 | Deep)cean | https://doi.or | 68.03 | MP-DEEP_MAG-0582.ENHA |
| No | 0 | Deep)cean | https://doi.or | 56.8 | MP-DEEP_MAG-0584.ENHA |
| No | 0 | Deep)cean | https://doi.or | 67.47 | MP-DEEP_MAG-0589.ENHA |
| No | 0 | Deep)cean | https://doi.or | 85.25 | MP-DEEP_MAG-0591.ENHA |
| No | 0 | Deep)cean | https://doi.or | 74.53 | MP-DEEP_MAG-0592.ENHA |
| No | 0 | Deep)cean | https://doi.or | 84.15 | MP-DEEP_MAG-0596.ENHA |
| No | 1 | Deep)cean | https://doi.or | 66.12 | MP-DEEP_MAG-0598.ENHA |
| No | 1 | Deep)cean | https://doi.or | 87.63 | MP-DEEP_MAG-0600.ENHA |
| No | 3 | Deep)cean | https://doi.or | 69.97 | MP-DEEP_MAG-0604.ENHA |
| No | 2 | Deep)cean | https://doi.or | 95.07 | MP-DEEP_MAG-0605.ENHA |
| No | 0 | Deep)cean | https://doi.or | 97.8 | MP-DEEP_MAG-0607.ENHA |
| No | 0 | Deep)cean | https://doi.or | 57.36 | MP-DEEP_MAG-0610.ENHA |

| | | | | | |
|---|---|---|---|---|---|
| No | 0 | Deep)cean | https://doi.or | 68.26 | MP-DEEP_MAG-0611.ENHA |
| No | 0 | Deep)cean | https://doi.or | 96.7 | MP-DEEP_MAG-0614.ENHA |
| No | 3 | Deep)cean | https://doi.or | 88.4 | MP-DEEP_MAG-0616.ENHA |
| No | 0 | Deep)cean | https://doi.or | 86.43 | MP-DEEP_MAG-0617.ENHA |

ANCED.TSV
ANCED.TSV
ANCED.TSV
ANCED.TSV
ANCED.TSV
ANCED.TSV
ANCED.TSV
ANCED.TSV
ANCED.TSV
ANCED.TSV
ANCED.TSV
ANCED.TSV

ANCED.TSV
ANCED.TSV
ANCED.TSV
ANCED.TSV
ANCED.TSV
ANCED.TSV
ANCED.TSV
ANCED.TSV
ANCED.TSV
ANCED.TSV
ANCED.TSV
ANCED.TSV
ANCED.TSV
ANCED.TSV
ANCED.TSV
ANCED.TSV
ANCED.TSV
ANCED.TSV
ANCED.TSV
ANCED.TSV
ANCED.TSV
ANCED.TSV
ANCED.TSV
ANCED.TSV
ANCED.TSV
ANCED.TSV
ANCED.TSV
ANCED.TSV
ANCED.TSV
ANCED.TSV
ANCED.TSV
ANCED.TSV
ANCED.TSV
ANCED.TSV
ANCED.TSV
ANCED.TSV
ANCED.TSV
ANCED.TSV
ANCED.TSV
ANCED.TSV
ANCED.TSV
ANCED.TSV
ANCED.TSV

ANCED.TSV
ANCED.TSV
ANCED.TSV
ANCED.TSV
ANCED.TSV
ANCED.TSV
ANCED.TSV
ANCED.TSV
ANCED.TSV
ANCED.TSV
ANCED.TSV
ANCED.TSV
ANCED.TSV
ANCED.TSV
ANCED.TSV
ANCED.TSV
ANCED.TSV
ANCED.TSV
ANCED.TSV
ANCED.TSV
ANCED.TSV
ANCED.TSV
ANCED.TSV
ANCED.TSV
ANCED.TSV
ANCED.TSV
ANCED.TSV
ANCED.TSV
ANCED.TSV
ANCED.TSV
ANCED.TSV
ANCED.TSV
ANCED.TSV
ANCED.TSV
ANCED.TSV
ANCED.TSV
ANCED.TSV
ANCED.TSV
ANCED.TSV
ANCED.TSV
ANCED.TSV
ANCED.TSV
ANCED.TSV

ANCED.TSV
ANCED.TSV
ANCED.TSV
ANCED.TSV
ANCED.TSV
ANCED.TSV
ANCED.TSV
ANCED.TSV
ANCED.TSV
ANCED.TSV
ANCED.TSV
ANCED.TSV
ANCED.TSV
ANCED.TSV
ANCED.TSV
ANCED.TSV
ANCED.TSV
ANCED.TSV
ANCED.TSV
ANCED.TSV
ANCED.TSV
ANCED.TSV
ANCED.TSV
ANCED.TSV
ANCED.TSV
ANCED.TSV
ANCED.TSV
ANCED.TSV
ANCED.TSV
ANCED.TSV
ANCED.TSV
ANCED.TSV
ANCED.TSV
ANCED.TSV
ANCED.TSV
ANCED.TSV
ANCED.TSV
ANCED.TSV
ANCED.TSV
ANCED.TSV
ANCED.TSV
ANCED.TSV
ANCED.TSV

ANCED.TSV
ANCED.TSV
ANCED.TSV
ANCED.TSV
ANCED.TSV
ANCED.TSV
ANCED.TSV
ANCED.TSV
ANCED.TSV
ANCED.TSV
ANCED.TSV
ANCED.TSV
ANCED.TSV
ANCED.TSV
ANCED.TSV
ANCED.TSV
ANCED.TSV
ANCED.TSV
ANCED.TSV
ANCED.TSV
ANCED.TSV
ANCED.TSV
ANCED.TSV
ANCED.TSV
ANCED.TSV
ANCED.TSV
ANCED.TSV
ANCED.TSV
ANCED.TSV
ANCED.TSV
ANCED.TSV
ANCED.TSV
ANCED.TSV
ANCED.TSV
ANCED.TSV
ANCED.TSV
ANCED.TSV
ANCED.TSV
ANCED.TSV
ANCED.TSV
ANCED.TSV
ANCED.TSV
ANCED.TSV

ANCED.TSV
ANCED.TSV
ANCED.TSV
ANCED.TSV
ANCED.TSV
ANCED.TSV
ANCED.TSV
ANCED.TSV
ANCED.TSV
ANCED.TSV
ANCED.TSV
ANCED.TSV
ANCED.TSV
ANCED.TSV
ANCED.TSV
ANCED.TSV
ANCED.TSV
ANCED.TSV
ANCED.TSV
ANCED.TSV
ANCED.TSV
ANCED.TSV
ANCED.TSV
ANCED.TSV
ANCED.TSV
ANCED.TSV
ANCED.TSV
ANCED.TSV
ANCED.TSV
ANCED.TSV
ANCED.TSV
ANCED.TSV
ANCED.TSV
ANCED.TSV
ANCED.TSV
ANCED.TSV
ANCED.TSV
ANCED.TSV
ANCED.TSV
ANCED.TSV
ANCED.TSV
ANCED.TSV
ANCED.TSV

ANCED.TSV
ANCED.TSV
ANCED.TSV
ANCED.TSV
ANCED.TSV
ANCED.TSV
ANCED.TSV
ANCED.TSV
ANCED.TSV
ANCED.TSV
ANCED.TSV
ANCED.TSV
ANCED.TSV
ANCED.TSV
ANCED.TSV
ANCED.TSV
ANCED.TSV
ANCED.TSV
ANCED.TSV
ANCED.TSV
ANCED.TSV
ANCED.TSV
ANCED.TSV
ANCED.TSV
ANCED.TSV
ANCED.TSV
ANCED.TSV
ANCED.TSV
ANCED.TSV
ANCED.TSV
ANCED.TSV
ANCED.TSV
ANCED.TSV
ANCED.TSV
ANCED.TSV
ANCED.TSV
ANCED.TSV
ANCED.TSV
ANCED.TSV
ANCED.TSV
ANCED.TSV
ANCED.TSV
ANCED.TSV

ANCED.TSV
ANCED.TSV
ANCED.TSV
ANCED.TSV
ANCED.TSV
ANCED.TSV
ANCED.TSV
ANCED.TSV
ANCED.TSV
ANCED.TSV
ANCED.TSV
ANCED.TSV
ANCED.TSV
ANCED.TSV
ANCED.TSV
ANCED.TSV
ANCED.TSV
ANCED.TSV
ANCED.TSV
ANCED.TSV
ANCED.TSV
ANCED.TSV
ANCED.TSV
ANCED.TSV
ANCED.TSV
ANCED.TSV
ANCED.TSV
ANCED.TSV
ANCED.TSV
ANCED.TSV
ANCED.TSV
ANCED.TSV
ANCED.TSV
ANCED.TSV
ANCED.TSV
ANCED.TSV
ANCED.TSV
ANCED.TSV
ANCED.TSV
ANCED.TSV
ANCED.TSV
ANCED.TSV
ANCED.TSV

\NCED.TSV
\NCED.TSV
\NCED.TSV
\NCED.TSV

**Sup. Table 5: Putative radiosensitive species from the HMB**

**Putative Radiosensitive Species from the HMB**

| Species | Phyla |
|---|---|
| Alcaligenes xylosoxydans xylosoxydans (Achromobacter xylosoxidans) | Proteobacteria |
| Alistipes shahii WAL 8301 | Bacteroidetes |
| Bacteroides fragilis (strain ATCC 25285 / DSM 2151 / CCUG 4856 / JCM 1101[illegible] | Bacteroidetes |
| Bacteroides stercoris | Bacteroidetes |
| Bacteroides thetaiotaomicron (strain ATCC 29148 / DSM 2079 / JCM 5827 / C | Bacteroidetes |
| Gordonibacter pamelaeae 7-10-1-b | Actinobacteria |
| Helicobacter pylori (strain ATCC 700392 / 26695) (Campylobacter pylori) | Proteobacteria |
| Klebsiella pneumoniae subsp. pneumoniae (strain HS11286) | Proteobacteria |
| Odoribacter splanchnicus (strain ATCC 29572 / DSM 20712 / CIP 104287 / JC[illegible] | Bacteroidetes |
| Parabacteroides merdae ATCC 43184 | Bacteroidetes |
| Prevotella copri DSM 18205 | Bacteroidetes |
| Neisseria cinerea | Proteobacteria |
| Neisseria flavescens NRL30031/H210 | Proteobacteria |
| Pseudomonas aeruginosa (strain ATCC 15692 / DSM 22644 / CIP 104116 / JC[illegible] | Proteobacteria |
| Veillonella parvula (strain ATCC 10790 / DSM 2008 / CCUG 5123 / JCM 12972 | Firmicutes |
| Campylobacter gracilis RM3268 | Proteobacteria |
| Capnocytophaga granulosa | Bacteroidetes |
| Capnocytophaga granulosa (strain ATCC 51502 / DSM 11449 / JCM 8566 / L[illegible] | Bacteroidetes |
| Cryptobacterium curtum (strain ATCC 700683 / DSM 15641 / CCUG 43107 / 1 | Actinobacteria |
| Eikenella corrodens CC92I | Proteobacteria |
| Prevotella denticola | Bacteroidetes |
| Prevotella nigrescens ATCC 33563 | Bacteroidetes |
| Tannerella forsythia (strain ATCC 43037 / JCM 10827 / CCUG 21028 A / KCTC | Bacteroidetes |
| Veillonella parvula ATCC 17745 | Firmicutes |
| Alistipes putredinis CAG:67 | Bacteroidetes |
| Bacteroides caccae | Bacteroidetes |
| Bacteroides ovatus (strain ATCC 8483 / DSM 1896 / JCM 5824 / BCRC 10623 / | Bacteroidetes |
| Bacteroides stercoris | Bacteroidetes |
| Bacteroides uniformis | Bacteroidetes |
| Capnocytophaga sputigena | Bacteroidetes |
| Eikenella corrodens | Pseudomonadota |
| Eubacterium siraeum CAG:80 | Bacillota |
| Helicobacter canadensis MIT 98-5491 | Pseudomonadota |
| Parabacteroides merdae ATCC 43184 | Bacteroidetes |

| Location |
| --- |
| Gut |
| Gut |
| Gut |
| Gut |
| Gut |
| Gut |
| Gut |
| Gut |
| Gut |
| Gut |
| Gut |
| Skin |
| Skin |
| Skin |
| Skin |
| Oral |
| Oral |
| Oral |
| Oral |
| Oral |
| Oral |
| Oral |
| Oral |
| Oral |
| NIH_ATCC |
| NIH_ATCC |
| NIH_ATCC |
| NIH_ATCC |
| NIH_ATCC |
| NIH_ATCC |
| NIH_ATCC |
| NIH_ATCC |
| NIH_ATCC |
| NIH_ATCC |
